# Literature-scaled immunological gene set annotation using AI-powered immune cell knowledge graph (ICKG)

**DOI:** 10.1101/2025.02.19.639172

**Authors:** Shan He, Yukun Tan, Qing Ye, Matthew Gubin, Hind Rafei, Weiyi Peng, Katayoun Rezvani, Vakul Mohanty, Ken Chen

## Abstract

Large scale application of single-cell and spatial omics in models and patient samples has led to the discovery of many novel gene sets, particularly those from an immunotherapeutic context. However, the biological meaning of those gene sets has been interpreted anecdotally through over-representation analysis against canonical annotation databases of limited complexity, granularity, and accuracy. Rich functional descriptions of individual genes in an immunological context exist in the literature but are not semantically summarized to perform gene set analysis. To overcome this limitation, we constructed immune cell knowledge graphs (ICKGs) by integrating over 24,000 published abstracts from recent literature using large language models (LLMs). ICKGs effectively integrate knowledge across individual, peer-reviewed studies, enabling accurate, verifiable graph-based reasoning. We validated the quality of ICKGs using functional omics data obtained independently from cytokine stimulation, CRISPR gene knock-out, and protein-protein interaction experiments. Using ICKGs, we achieved rich, holistic, and accurate annotation of immunological gene sets, including those that were unannotated by existing approaches and those that are in use for clinical applications. We created an interactive website (https://kchen-lab.github.io/immune-knowledgegraph.github.io/) to perform ICKG-based gene set annotations and visualize the supporting rationale.

## Introduction

Immune cells are central to human health. In cancer, the interaction between immune and tumor cells influences tumor progression and treatment outcome^1–3^. The complex transcriptional programs in individual immune cell-types ultimately govern their function and consequently their roles within the tumor microenvironment (TME). To prevent and effectively treat diseases, it is imperative to develop methods that can systematically grow knowledge and deliver a precise, comprehensive understanding of the functional programs in diverse immune cell-types.

The burgeoning of omics studies applying single-cell and spatial omics technologies in human specimens has revealed unprecedented complexity and heterogeneity of signaling, interaction, and regulatory patterns in human tissues and disease states. For example, in immuno-oncological research, several pan-cancer cell atlases have been published that catalogue condition-specific cell types, cell states and associated molecular signatures^4–12^. Multiple databases were curated to annotate such signatures. For example, RNA profiling of immune-perturbed bulk specimens has been performed to catalogue immune-related cell states (MSigDB C7^13^). Functional annotation databases such as Hallmarks and KEGG have been widely used to annotate gene sets. Although valuable, these databases lack the granularity and richness to power the interpretation of gene sets observed in rich, single-cell resolution omics studies. They appear particularly limited in immunological knowledge, thus annotations obtained using these databases often lack the necessary granularity or are biased towards certain aspects of biology (e.g., metabolism)^13,14,15^. This lack of capacity is a major bottleneck in data-driven discovery in immunology. As a result, the meaning of the discovered gene sets is laboriously reasoned based on researchers’ empirical knowledge, making annotations often arbitrary, incomplete, and/or biased.

On the other hand, a large body of knowledge about individual genes and their functions in specific biological contexts exist in the literature, produced largely via hypothesis-driven research. These knowledge are often of high quality, validated via orthogonal experiments, and described in rich language, making them valuable for annotating and interpreting novel gene sets. However, this approach requires labor-intensive, domain-specific curation to consolidate annotations as a standardized, authoritative knowledgebase for systematic use. Breakthroughs in AI technologies such as large language models (LLMs) have made it possible to overcome the challenge in leveraging literature for gene set annotation. We hypothesize that by applying LLMs, we can create a well-curated knowledgebase specifically designed to describe human immunity and to infer rich molecular-cellular-function relationships that are not available in current databases. Several recent approaches^16,17^ have been developed that directly apply LLM as a knowledgebase and language model to interpret gene sets. However, these approaches suffer from the inherent limitations of LLMs, including a lack of transparency and accountability in reasoning, often resulting in hallucinations.

As an alternative approach, we have constructed immune cell knowledge graphs (ICKGs) on over 24,000 PubMed-indexed abstracts using various AI techniques including LLMs. These ICKGs cover major human immune cell types, containing entities such as genes, protein complexes, diseases, cell types, pathways, and relevant biomedical concepts, as well as relations such as activation and inhibition, all of which can be visualized and traced back to the supporting publications. With ICKGs, knowledge graph reasoning (KGR)^18–21^ can be performed to uncover hidden immunological insights and perform gene set interpretations. We rigorously validated each step of ICKG construction and evaluated the KGR performance using independent cell-type specific, functional perturbation data. We conclude that performing KGR on ICKGs can enhance gene set annotations with rich and highly specific immunological languages that can substantially advance the immunological knowledge acquisition, interpretation, and translation.

## Results

### Construction of ICKGs to represent immune systems complexity

The overall schematic for ICKG construction follows **Figure 1a**. To enhance context-specificity and achieve cellular granularity, we constructed ICKGs for specific cell types including T cells (including CD4/CD8^+^ T cells and regulatory T cells), natural killer (NK) cells, B cells and macrophages. To obtain the most up-to-date publications on immune cells in the setting of cancer, we searched PubMed and retained abstracts published between 2020 and 2024 using “cell type” and “cancer immunotherapy” as the keywords (**Figure 1b**). Genes, diseases, and cell types are important entities to be extracted from abstracts, as they provide both the molecular and functional basis to characterize immune cells. To this end, we finetuned respective named-entity-recognition (NER) tasks using a pre-trained BioBERT model with corresponding training datasets and selected the best one based on highest precision (**Figure 1c**, **Methods**). Ultimately, BioBERT models finetuned on BioNLP13PC^22^, NCBI^23^, and GENIA were selected to perform the NER task for gene, disease, and cell type recognition, respectively, with a precision of 87.87%, 85.23%, and 61.77% (**Table 1**).

**Figure 1.**
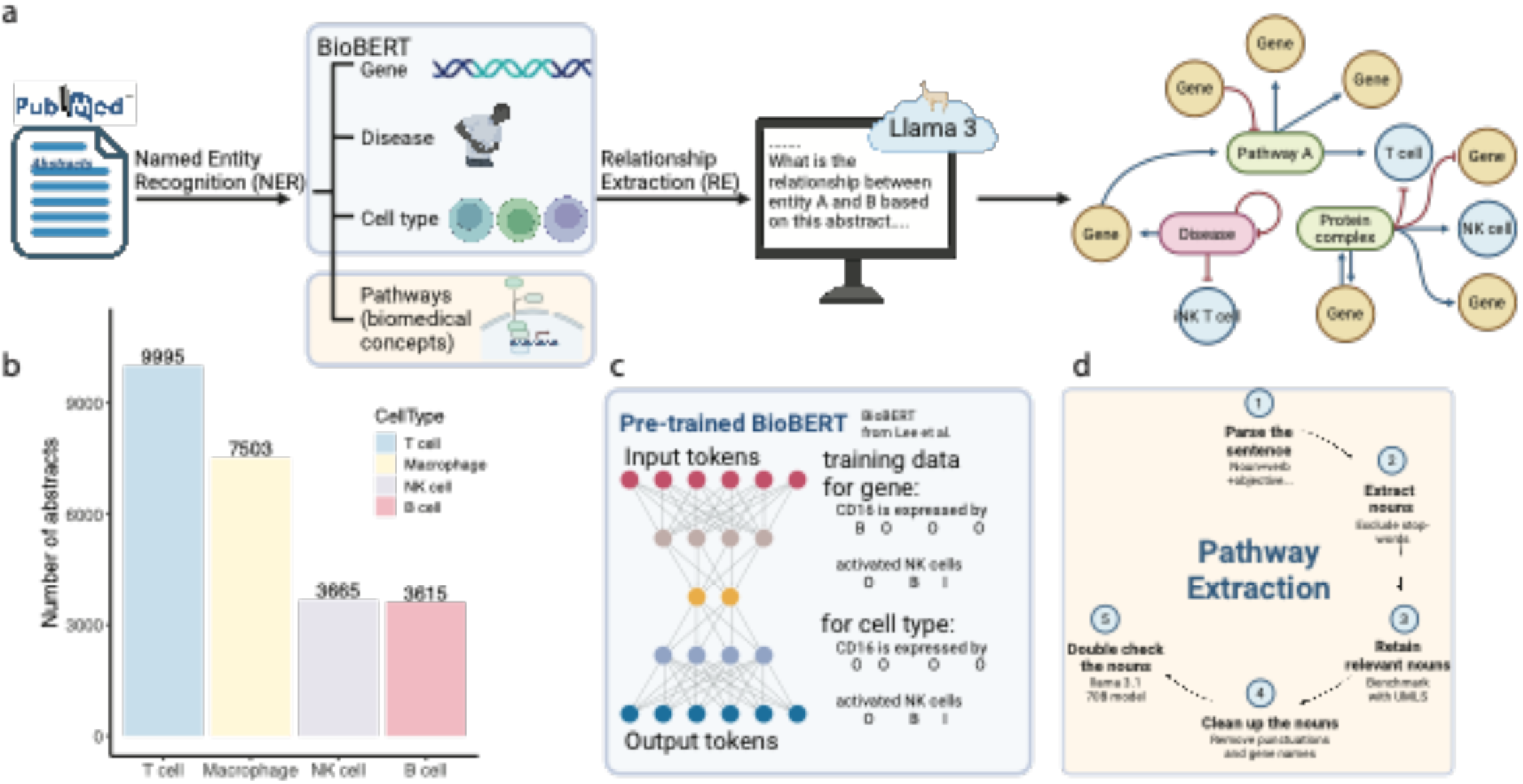
Schematic for Immune Cell Knowledge Graph Construction. (a) Overall workflow: abstracts download based on keywords. Named entity recognition (NER) based on finetuned BioBERT model. Relationship extraction based on prompt engineering. Network assembly based on relations (nodes are colored based on node types and edges are colored based on relationship). (b) Number of abstracts extracted for each immune cell type of interest. (c) BioBERT methodology demonstration. (d) Step-by-step explanation for pathway extractions from abstracts.

**Table 1.** Performance Evaluation for Named Entity Recognition (NER) across di=erent entity types and training datasets.

| Training data | No. of words | Precision | Recall | F1 Score |
| --- | --- | --- | --- | --- |
| Gene name NER |  |  |  |  |
| BC2GM<sup>24</sup> | 148,529 | 82.85 | 84.21 | 83.52 |
| JNLPBA | 118,582 | 70.78 | 83.07 | 76.43 |
| BioNLP13PC* | 47,896 | **87.87** | **90.09** | **88.97** |
| Disease name NER |  |  |  |  |
| BC5CDR<sup>25</sup> | 129,562 | 85.00 | 87.07 | 86.02 |
| **NCBI\*** | 25,438 | **85.23** | **88.96** | **87.05** |
| Cell type name NER |  |  |  |  |
| **GENIA\*** | 50,031 | 61.77 | 73.42 | 67.09 |
| Pathways/Biomedical phrases NER |  |  |  |  |
| GENIA | 50,031 | 52.63 | 60.58 | 56.33 |
| **Rule-based + prompt\*** | 48,222 | **72.81** | **80.64** | **76.53** |

Pathways (or biological concepts) are also important entities that need to be extracted from abstracts as they provide the basis for new vocabularies that we can use to annotate a gene set. However, definitions of pathways are vague, challenging our entity extraction approach. In addition, since data with pathway terms pre-labeled are rare with only GENIA providing limited pathway labels, ground truth is lacking to validate any pathway extraction approach. To resolve these issues, we established a lenient but effective definition of “pathways” as any informative biomedical noun phrase in an abstract. Since abstracts are usually written in succinct sentences, we designed a rule-based approach based on the grammatical structure of each sentence to extract biomedical phrases followed by prompt engineering to retain the biomedically relevant nouns (**Figure 1d, Methods**). We also manually annotated a database with 48,000 words based on our definition and found that this combined method was able to extract concise and informative biomedical phrases from abstracts with a precision of 72.81% (**Table 1**).

KGs constructed for other purpose, e.g., drug repurposing^26–31^, often contain complex edge properties^28,30^, which may hamper effective reasoning and interpretations. Conversely, those with simple edge properties, extracted from relationship databases^27,29,31^, can limit novel discoveries. To achieve a tradeoff between functionality and practicality, we defined the edges with two biologically interpretable properties: activation versus inhibition. We asked the LLM to infer the relationship between the entities based on the abstract that they were extracted from^32^.

We employed a zero-shot prompt engineering approach using the open-source Llama 3.1 model (70B parameters) with a carefully designed prompt (**Methods**). We requested the model to infer the directional relationship (i.e., activation, inhibition, or no association) between pairwise entities base only on the abstract. We evaluated Relationship Extraction (RE) performance by benchmarking our inferred relationships against gene-gene and gene-cell type databases using KEGG^15^ and the Immune Cell Atlas^33^, respectively. For gene-gene relationships, we test if ICKG can capture KEGG-defined activation and inhibition interactions between gene pairs. For gene-cell type relationships, we test if the genes enriched in major immune cell types are captured by ICKG as genes that directly activate the cell type node. We extracted 58,456 pairwise directional relationships from 361 KEGG pathways and simplified the relationships into either activation or inhibition based on interaction types (**Methods**). The Immune Cell Atlas was profiled from more than 300,000 individual immune cells extracted from 16 different heathy tissues, and we compared our gene-cell type relationship with the cell type DEGs from this atlas (**Methods**). **Table 2** includes a summarized performance across the different RE tasks, with precision ranging from 77.18% to 87.66%.

**Table 2.** Performance evaluation for relationship extraction for gene-gene and gene-cell type across the di=erent immune cell-specific knowledge graphs.

| Cell type | Precision | Recall | F1 Score |
| --- | --- | --- | --- |
| Gene-Gene relationship extraction |  |  |  |
| T | 83.08% | 95.16% | 88.70% |
| B | 85.65% | 93.43% | 89.40% |
| Macrophage | 80.57% | 90.36% | 85.19% |
| NK | 84.47% | 96.80% | 90.22% |
| Gene-Cell type relationship extraction |  |  |  |
| T | 77.18% | 75.43% | 76.29% |
| B | 81.08% | 86.77% | 83.83% |
| Macrophage | 82.67% | 84.26% | 83.46% |
| NK | 87.66% | 82.79% | 85.16% |

The constructed knowledge graphs have nodes of various types (i.e., gene, disease, cell type, and biomedical concepts) and edges of either activating or inhibiting relationships with weights representing the number of abstracts that mention such relationships. Each count on an edge corresponds to a real PubMed publication that was identified to support the relationship and is manually verifiable to ensure accuracy and verifiability. **Table 3** summarizes basic node and edge information for the constructed knowledge graphs. For visualization purposes, we have presented the core section of the T and the NK cell graphs with thick edges, representing relationships with more literature supports, in **Figure 2a** and **Figure 2b**, respectively. Subgraphs for the B cell and the macrophages are presented in **Figure S1a-b**. To summarize some of the constructed relationships between genes and phenotypes, we leveraged the PageRank algorithm and extracted genes that are significantly associated with major cell types and TCGA cancer subtypes in respective cell type graphs (**Figure 2c-f, Figure S1c-f, Methods**).

**Figure 2.**
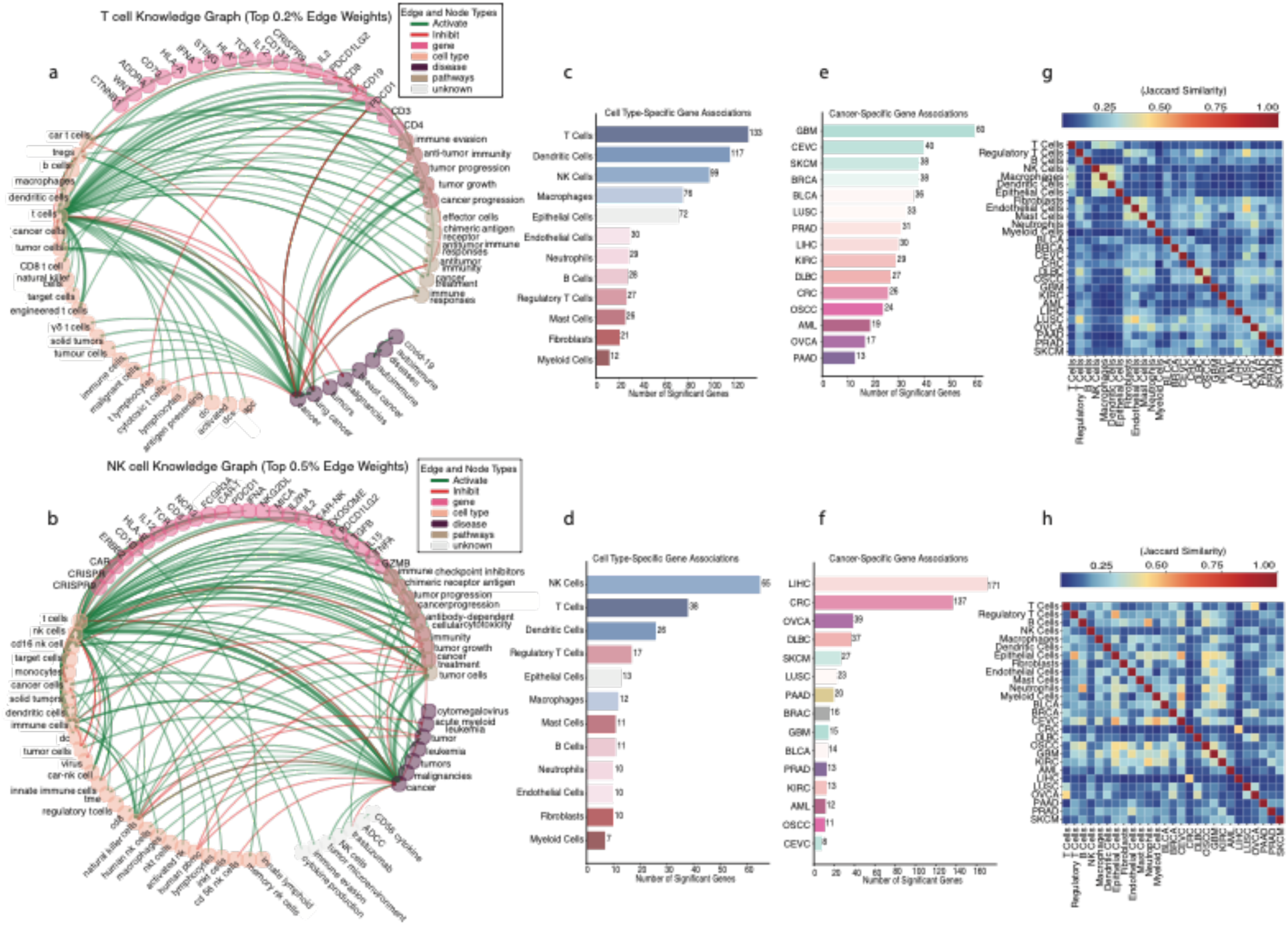
(a-b) T and NK cell specific knowledge graph subset containing only edges with top 0.5% weight. Nodes are colored based on node types (gene, disease, cell type, pathways and others). Edges are colored based on activation or inhibition between the two connecting nodes. The number on the edge represents the number of published abstracts that support such relationship (c-d). Number of significantly associated genes with each major cell types derived from T and NK cell specific knowledge graph (e-f). Number of significantly associated genes with each major TCGA cancer type according to PageRank scores derived from T-and NK-cell specific knowledge graphs. (g-h) Pairwise jaccard distance (measure for the number of overlapping genes) across cancer types and cell types derived respectively from T-and NK-cell specific knowledge graphs.

**Table 3.**
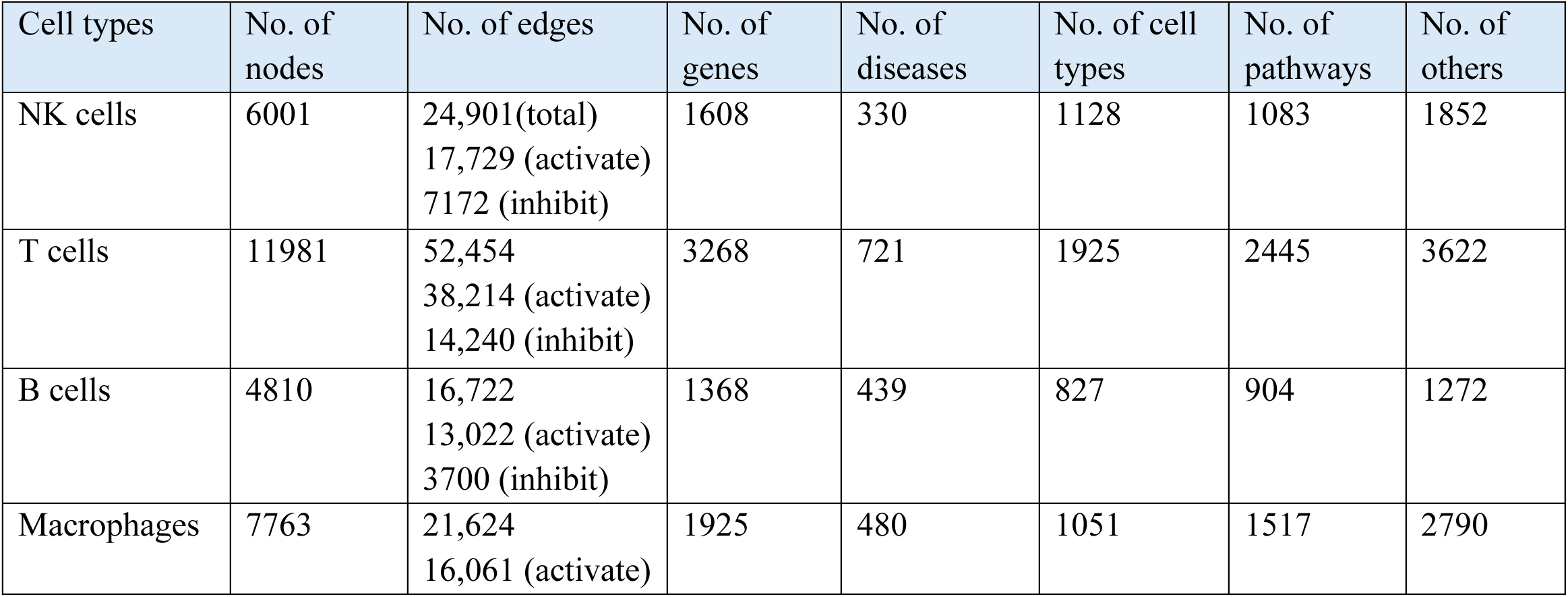
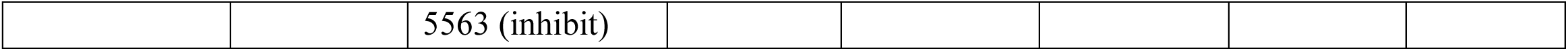
Basic information for the constructed Knowledge Graphs.

| Cell types | No. of nodes | No. of edges | No. of genes | No. of diseases | No. of cell types | No. of pathways | No. of others |
| --- | --- | --- | --- | --- | --- | --- | --- |
| NK cells | 6001 | 24,901(total)<br>17,729 (activate)<br>7172 (inhibit) | 1608 | 330 | 1128 | 1083 | 1852 |
| T cells | 11981 | 52,454<br>38,214 (activate)<br>14,240 (inhibit) | 3268 | 721 | 1925 | 2445 | 3622 |
| B cells | 4810 | 16,722<br>13,022 (activate)<br>3700 (inhibit) | 1368 | 439 | 827 | 904 | 1272 |
| Macrophages | 7763 | 21,624<br>16,061 (activate) | 1925 | 480 | 1051 | 1517 | 2790 |

|  |  |  |
| --- | --- | --- |
|  |  | 5563 (inhibit) |

Our knowledge graphs effectively capture major cell and cancer types. As expected, we found that more genes are significantly associated with T cells in the T cell graph, and more genes are significantly associated with NK cells in the NK cell graph. Interestingly, cancer types are associated with variable numbers of genes in the T cell graph vs those in the NK cell graph. For example, Glioblastoma (GBM) is associated with the highest number of genes in the T cell graph, whereas in the NK cell graphs, Liver Hepatocellular Carcinoma (LIHC) appears to be the most associated cancer type. By analyzing these gene associations, we gain insight into the research landscape—identifying which genes researchers are studying in relation to various immune cell types and cancer types. **Figures 2g-h and Figure S1g-h** summarize disease and cell type similarities based on shared genes in the corresponding graphs, highlighting genetic commonality across cell types and cancer types.

### ICKGs enable verifiable, graph-based reasoning

With the knowledge graphs constructed, reasoning can be performed based on the connections and attributes in the graph to infer new information or predict outcomes. The performance of a knowledge graph reasoning (KGR) task, however, can vary depending on the quality (e.g., accuracy, completeness) of the graph and the efficacy of the inference method. Methods for KGR range from rule-based logical reasoning to graph neural networks^21^. Among the available methods, PageRank has been widely used to perform graph-based reasoning and to quantify the influence of each node within a network, thereby enabling more informed inferences based on the relative importance and connectivity of nodes in the complex systems.

To validate the quality of ICKGs and identify a suitable KGR method, we compared the results of predicting cytokine stimulation or gene knockout effects using PageRank on ICKGs, adjusted random walk on ICKGs, and PageRank on randomly shuffled graphs. For cytokine stimulation, we took the cytokine dictionary data^34^, in which the authors profiled the transcriptomic changes of 17 different immune cell types when they were stimulated with 86 different cytokines. For gene perturbations, we took a scCRISPR screening of T cells^35^, in which the authors performed single-cell knockout screen of 180 transcription factors (TFs) in primary CD8+ T cells. To establish the ground truth, we performed differential gene expression analysis between cells with and without experimental manipulations and identified differentially expressed genes (DEGs) in both datasets (**Methods**). We then locate each perturbation, whether cytokine or gene, on the respective ICKG and reason through each method respectively to find the activated and inhibited genes (**Methods**).

Figure 3a summarizes the distribution of AUC values across perturbations tested. The results demonstrate that PageRank, when applied to the respective ICKG, consistently yields significantly higher AUC values than the control experiments using either random walk or random graphs. To further understand the suboptimal performance for some perturbation predictions, we dichotomized the perturbations into low and high AUC groups using a bimodal gaussian mixture model (**Methods**) and found that those perturbations achieving high AUC values involve genes with a significantly higher degree centrality in ICKGs (Figure 3b), reflecting the number of pairwise relationships involving this gene captured by the literature. Given the impact of degree centrality on reasoning performance, we can infer that enhancing reasoning performance might be feasible through the continual integration of additional information into the ICKGs. Taken together, the results verify that we have effectively integrated textual information into ICKGs and that PageRank is a reasonable approach to enable ICKG-based gene set annotation.

**Figure 3.**
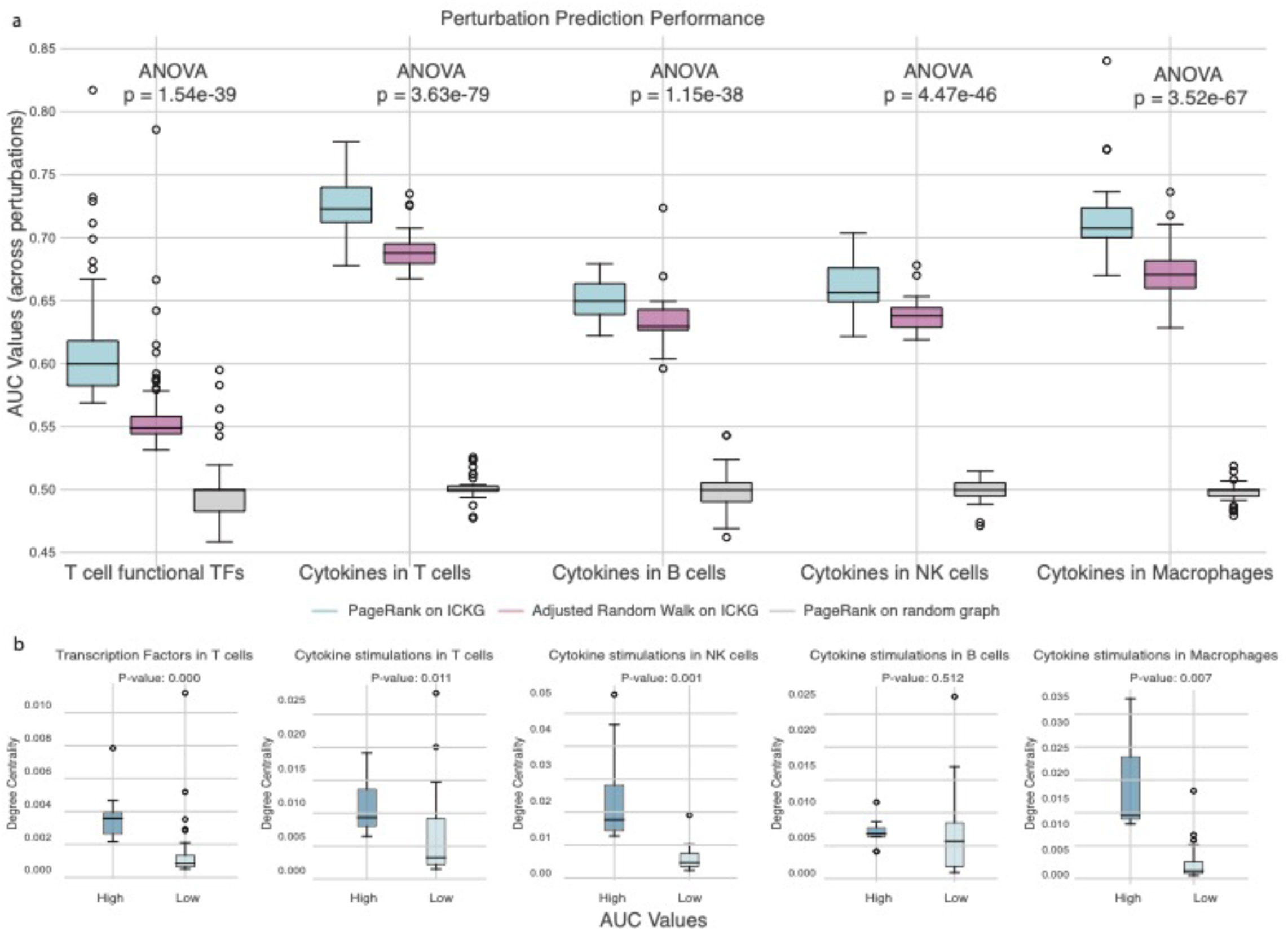
(a) Distribution of AUC values measured by three di=erent reasoning approaches. Blue represents PageRank methods on respective ICKGs. Pink represents adjusted random walk on the respective ICKG. Grey represents PageRank on a randomly shu=led graph. (b) The degree centrality of the perturbed genes in ICKGs for experiments achieving high vs those achieving low AUC values.

### ICKG-based gene set annotation outperforms traditional methods

Having established the authenticity of ICKGs and the reasoning method, we now return our focus to our goal of enhancing gene set annotation.

The constructed ICKGs contain rich and specific vocabularies to describe biomedical concepts and pathways that allow researchers to easily conceptualize gene set functions. Since ICKGs feature relationships among genes, pathways, and biomedical concepts, we can effectively leverage their interconnectivities to perform gene set annotation using PageRank^36^ (**Methods**). For evaluation, we obtained gene sets from the meta programs (MPs) of four immune cell types (12 B, 10 1CD4^+^ T cell, 12 CD8^+^ T cell, and 13 macrophage gene sets) constructed by Gavish et al^37^, four tumor-reactive T cells signature^38–41^, and four NK gene sets related to cytotoxicity, inhibitory, stimulatory, and tumor-induced stress functions^42^. These gene sets reflect important molecular programs in immunology: Gavish’s MPs were derived by integrating 77 studies covering 24 tumor types, presenting conserved transcriptomic programs in different immune cell types; the tumor-reactive T cells signatures were derived experimentally to characterize clonally expanded T cells; and NK genes sets were derived from 716 patients with 24 cancer types, thus accurately annotating the functions of these gene sets in terms of rich and relevant immune languages would indicate the broad applicability of our approach. For comparison, we first annotated the gene sets using the traditional over-representation analysis (ORA) based on Hallmarks, REACTOME, KEGG, and Wiki-Pathways from the MSigDB database^43^, which sum up to 2515 pathways, comparable to the number of pathway nodes in ICKGs. We then annotated the same gene sets (N = 55) based on the respective ICKGs. **Table S1-S2** summarizes the enriched terms for each gene set from ICKGs and ORA, respectively.

To quantify the capability of ICKGs to annotate gene sets into relevant biological functions, we projected the functional annotations obtained via individual genes, ORA and ICKGs onto the BERT embedding space and assessed their similarities (**Methods**). It has been shown that LLMs such as BERT can meaningfully represent biological terms in their embeddings^44^. Compared with ORA annotations, we found that ICKG annotations are as relevant (similar distances between cluster centroids in the embedding space) to the gene set of interest but are more specific, with a significantly smaller within-cluster sum of square (Figure 4a-b**, Figure S2**), indicating more functionally coherent annotations. For example, when we map the embeddings for an antigen-specific T cell signature^39^, we see ICKG annotations are more condensed with highly relevant annotations, such as ‘vaccine-mediated antibody response’ and ‘antigen-induced exhaustion’, while ORA-annotations are more spread out with functions not immediately relevant to the core functionality of the gene set (Figure 4c). In addition, the annotations generated by ICKGs are shorter in string length (average length in characters is 25 for ICKG terms and 47 for ORA terms), suggesting that our LLM approach is able to use concise language while capturing the core functionality.

**Figure 4.**
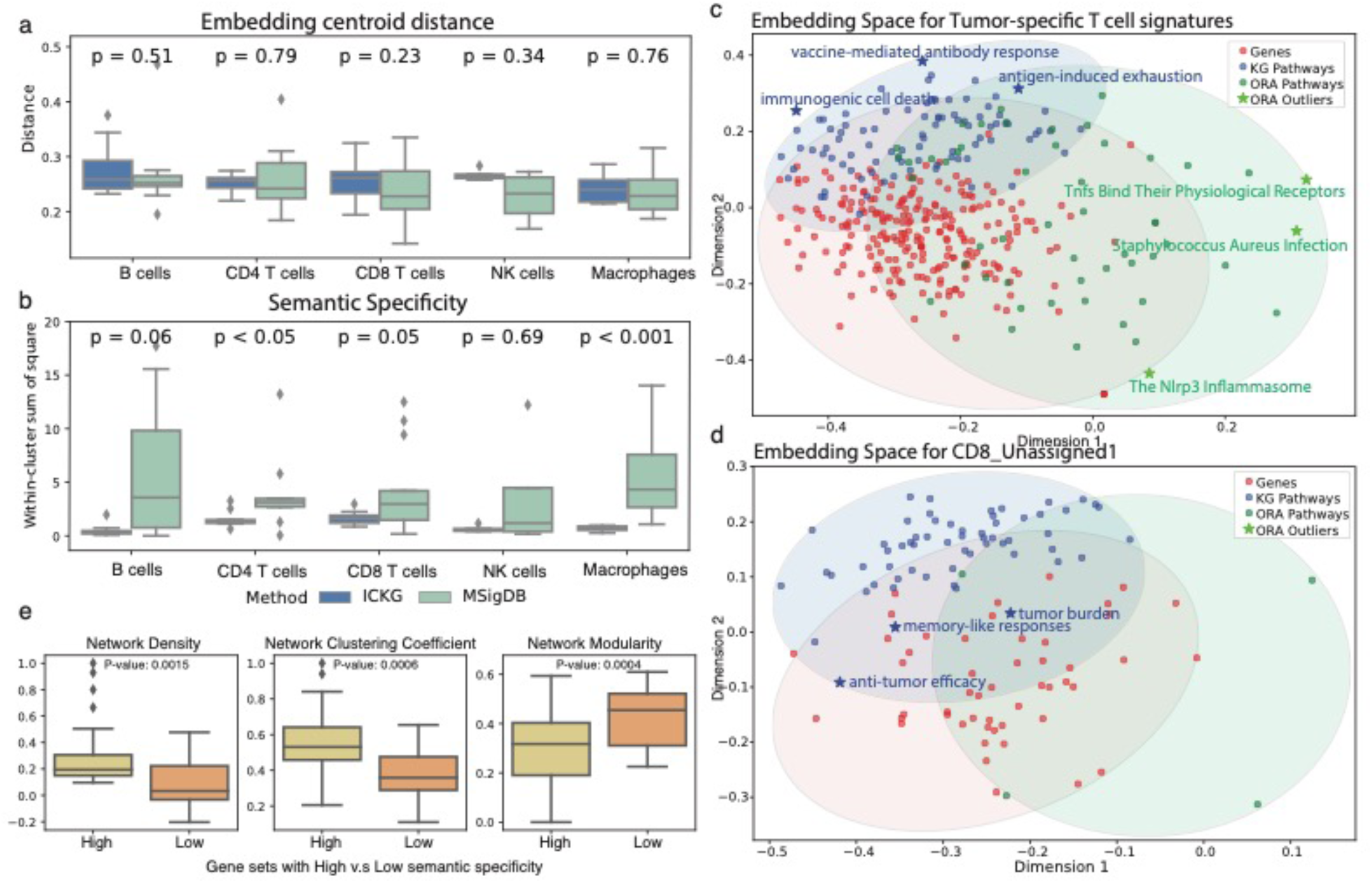
(a) Comparison of centroid distance resulting from the ICKG PageRank and MSigDB ORA method. (b) Comparison of semantic specificity resulting from the ICKG PageRank and MSigDB ORA method. Semantic specificity is approximated by within-cluster sum of square (WCSS) in the BERT embedding space, representing the degree of coherence of the functional annotation, with lower values indicating more homogeneous (more specific) annotations. Statistical significance between ICKG and MSigDB was assessed using the Mann-Whitney U test for each cell type. Metrics are shown separately for B cells, CD4 T cells, CD8 T cells, NK cells, and Macrophages. (c-d) Two-dimensional representation of genes in antigen-specific T cell gene set and CD8_unassigned1 gene set and respective enriched pathways in the BERT embedding space. Individual genes (red dots) are shown alongside two types of pathways: ICKG Pathways (blue dots) and ORA Pathways (green dots). The proximity between dots suggests semantic or functional proximity. (e) Comparison of PPI network density, average clustering coe=icient, and modularity between gene sets with high vs low semantic specificity.

Notably, ICKGs are capable of annotating gene sets when other approaches fail. Gavish et al manually summarized the enrichments into one term based on empirical knowledge, but they were unable to annotate three T cell related MPs, highlighting the challenges and limitations of human summarization when the enriched terms are very spread out in functionality, which is usually the case when we perform ORA-based annotation using MSigDB (Figure 4b). However, our approach was able to confidently annotate all three MPs. According to the T cell KG (**Table S1**), CD4_Unassigned is enriched in cytotoxicity and memory, suggesting a tissue resident T cell phenotype. When we further looked into its constituent genes, we found this gene set is enriched in genes related to cytoskeletal maintenance and mobility (ARPC4, ARPC1B, ARPC5, CAP1, CORO1A, CORO1*B*), suggesting interaction with the surrounding cells and extracellular matrix, which are critical for T cells to migrate, adhere, and maintain their positions in tissues^45^. The CD8_Unassigned1 and CD8_Unassigned2 likely represent cytotoxic CD8 T cells but differ in their functional states and roles. According to the ICKG, CD8_Unassigned1 is enriched in ‘anti-tumor efficacy’ and ‘tumor burden’ (Figure 4d), likely representing CD8 T cells in an intermediate or early stage of exhaustion with cytotoxic potential (GZMK, KLRG1) and memory-like capacity (CD27, IL7R), though facing metabolic stress (TXNIP, UCP2) as they progress towards exhaustion. In contrast, CD8_Unassigned2 is enriched in ‘cytotoxic extracellular vesicles’ and ‘tumoricidal factors’, suggesting tissue-resident memory (TRM) phenotypes with genes supporting long-term survival and immune surveillance (TXN, PARK7, ARPC1B).

To further substantiate the semantic specificity metric obtained from BERT, we dichotomized the gene sets by their semantic specificity using a Gaussian mixture model and examined the distribution of the genes in the protein-protein interaction (PPI) network for each gene set (**Methods**). We found that gene sets in the low semantic specificity group exhibit significantly lower network density and clustering coefficient, and they have higher modularity scores, indicating higher degree of heterogeneity (Figure 4e).

## Discussion

In this study, we developed a knowledge-graph based approach that performs rich, cell-type-specific, and human-verifiable gene set annotation using AI. We constructed ICKGs for four major immune cell types: T cells, B cells, NK cells, and macrophages by parsing and integrating immune cell related abstracts published on PubMed from January 1, 2020 to September 1, 2024. We demonstrated that ICKGs can systematically annotate novel gene sets using immune specific vocabularies in the literature, achieving granularity and accuracy beyond existing approaches. As new gene sets are being discovered from large-scale single-cell and spatial omics studies, our approach has the potential to broadly facilitate data interpretation, hypothesis generation and testing in immunological research and therapeutics development. For example, our work can be applied to the Immune Cell Atlas projects^46,47^ to better characterize immune cellular subtype based on respective DEGs.

Our ICKG-based annotation is transparent and verifiable, overcoming limitations in approaches that directly apply LLMs or deep learning methods for annotation. Unlike LLMs that are prone to generate non-reproducible or “hallucinated” answers, our approach ensures that all evidence involved is traceable to specific, peer-reviewed publications. This preserves scientific rigor and ensures reliable knowledge acquisition. In addition, the reasonings behind each annotation task can be visualized through query-able ICKG subgraphs, providing clear and interpretable insights.

Pathway NER is challenging as the field continues to lack systematically labeled training data. In working to advance this field, we established clear and universally generalizable criteria and manually labelled more than 48,000 words instrumental for the training and finetuning of pathway NER models. We also formulated a framework that combines rule-based methods and prompt engineering to robustly extract biomedical concepts from texts that contain phrases that are both rich and specific in immunobiology descriptions. Automating biomedical concept extraction from the literature allows for continuous expansion of our knowledgebase to facilitate data interpretation.

While ICKGs constructed in this study can be powerful tools, there are several limitations users should be cognizant of. First, the knowledge graph reflects so-called literature biases, overrepresenting frequently studied subjects and recent publications. While normalization techniques and expanded coverage of older literature could address these biases, our current approach effectively captures the research landscape, offering a good starting point for further development. Second, while limiting our analysis to abstracts instead of full texts constrained the volume of information present in the original studies, focusing on abstracts enhanced the robustness of pathway NER, avoiding complex grammatic and linguistic variability in parsing the full text. This approach also mitigated issues related to double-counting, which can occur when multiple sections of a paper redundantly report the same data, leading to overrepresentation in the analysis. Third, we tasked LLMs to dichotomize entity relationships into either activation or inhibition. Although this facilitates efficient and intuitive reasoning, the simplification can limit biological granularity, thus this approach can potentially benefit from further methodological innovation.

While the immediate focus of our work is on major immune cell types, the methodology is generalizable to other cell types or other omics settings to broadly facilitate molecular analysis, outcome prediction, genetic screening, and risk assessment.

Overall, our study highlights promising advancements in leveraging AI and graph-based reasoning to enhance gene set annotations, addressing a critical bottleneck in understanding novel molecular findings from single-cell and spatial transcriptomic studies. Future efforts should focus on integrating diverse data sources, refining algorithms for achieving accurate, efficient reasoning, and fostering collaboration between computational analysis and experimental validation to enhance utility and accuracy. To facilitate user-friendly access, we have created a web-based application with four ICKGs embedded that allows for customized cell type specific gene set annotations (https://kchen-lab.github.io/immune-knowledgegraph.github.io/). Researchers can leverage the graphs to systematically query relevant knowledge encoded in recent literature to perform cell-type specific gene set annotations.

## Methods

### Named entity recognition (NER) task

We fine-tuned the BioBERT-Large v1.1 model, which was pre-trained on 1 million PubMed articles and built on the BERT-large-Cased architecture with a custom 30,000 vocabularies, using PyTorch. The fine-tuning was performed using multiple NER datasets that were previously labeled to recognize relevant entities. For gene names, we used BC2GM, BioNLP13PC, and JNLPBA. For disease names, we used NCBI and BC5CDR. For cell type names, we used GENIA. We finetuned the BioBERT model with these datasets and selected the final finetuned model for each NER task base on precision (i.e., the proportion of correctly recognized entities out of all recognized entities), ensuring optimized performance in domain-specific NER tasks.

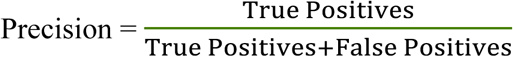

### Pathway extraction

We developed a pipeline to extract pathways and relevant biomedical phrases from PubMed abstracts using spaCy^48^ and a custom set of rules based on grammatical structures. We split each abstract into sentences, each of which is then processed using the en_core_web_sm language model in spaCy. We extracted noun phrases that are not single tokens and excluded common stop words, appending these to a set of candidate pathway terms. To refine the extracted terms, we utilized the Unified Medical Language System (UMLS) API query to identify biomedical terms, applying Levenshtein similarity to ensure close matches to known biomedical terms. Terms are further filtered using a set of criteria that prioritize biological relevance, such as specific suffixes (“ase,” “in,” “ion,” “itis,” “osis,” “oma,” “icity,” and “pathy”), substrings (“cell,” “gene,” “protein,” “receptor,” “antibody,” “immune,” “tumor,” “cancer,” “virus,” and “bacteria”) related to biology, and exclusion of non-biomedical entities. Finally, we fed the cleaned terms into the llama 3 70B-parameter model, requesting it to review the terms and output only the informative ones. This approach allows us to systematically extract and validate pathways, improving the accuracy of entity recognition in biomedical texts.

### Relationship extraction (RE)

To infer pairwise relationships between extracted entities in each abstract, we utilized a zero-shot prompt engineering approach. First, all possible pairs of terms were generated using pairwise permutations of the extracted entities. For each entity pair, we provided a detailed prompt to the Llama 3.1 model (70B parameters), instructing it to determine the directional relationship between the two terms. The model inferred whether the first term activated, inhibited, or had no association with the second term based on the content of the abstract. Clear guidelines were given to the model, specifying that relationships should only be inferred from the first term to the second, with ambiguous or biologically irrelevant relationships marked as “no association.” The model’s output was parsed and filtered to retain only relationships classified as “Activate” or “Inhibit,” which were subsequently used for downstream analyses.

To evaluate the performance of RE, we leveraged different databases. Gene-gene relationships were benchmarked with the KEGG database and Gene-cell type relationships were validated using an Immune Cell Atlas. For gene-gene relationships, we used the *KEGGREST* package in R and extracted more than 55,000 pairwise relationships between genes based on 361 KEGG pathways. For comparability, we simplified the relationships into either activation or inhibition based on interaction types: activation will encompass all of the following KEGG-defined interaction types: “activation,” “activation-indirect effect,” and “activation-indirect;” inhibition includes the following: “inhibition,” “inhibition-indirect effect,” “inhibition-repression,” and “repression-indirect effect,” which sum up to 15,832 activate and 3751 inhibit relationships. We output all the gene-gene pairs along with their directionality from ICKG and constructed a confusion matrix based on the overlapping gene-gene pairs.

|  | Activate in KG | Inhibit in KG |  |
| --- | --- | --- | --- |
| Activate in KEGG | A | B | A+B |
| Inhibit in KEGG | C | D | C+D |
|  | A+C | B+D | E = No. of gene-gene pairs that exist in both KG and KEGG |

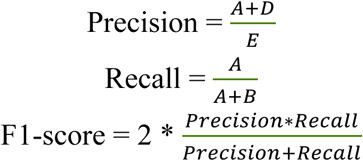

For gene-cell type relationships, we output all the gene nodes that were direct neighbors of that cell type of interest in the KG and benchmarked them with differentially expressed genes (DEGs) for the corresponding cell type of interest using the immune cell atlas dataset, then we constructed a confusion matrix.

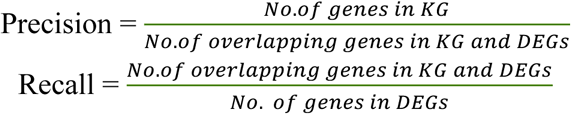

For both RE evaluations, we used the *select* function from the *AnnotationDbi* package in R to output a possible alias for each gene, and we considered a match if a gene matched any of the aliases of the genes in the benchmarking database.

### PageRank

We implemented PageRank using the NetworkX package (version 2.6) in Python (version 3.8.18). We constructed our network graph using NetworkX, and we then applied the PageRank algorithm with the following function: *nx.pagerank(G, alpha=0.85, personalization=phenotype_node)*, where G is our weighted network, alpha is the damping factor (default to 0.85), personalization is set to assign higher importance to specific nodes of interest (a list of genes if we are interested in gene set annotation, one gene if we are interested in its downstream impact). Other parameters in this function were set to default values.

### Permutation test

We implemented a permutation test to assess the significance of PageRank scores for the selected nodes in the network. We first computed personalized PageRank scores using the input gene set as personalization nodes. We then performed 1000 permutations, each time randomly selecting a gene set of the same size as the input set and computing PageRank scores. For each selected node, we compared its actual PageRank score to the distribution of scores from the permutations. P-values were calculated as n/1000, where n is the number of permutations yielding a score greater than or equal to the actual score. This approach allowed us to determine whether the PageRank scores for pathway nodes were significantly higher than expected by chance, given the network structure. Additionally, we computed 95% confidence intervals for each selected node’s PageRank score using the 2.5th and 97.5th percentiles of the permutation distribution.

### Adjusted Random Walk

We implemented adjusted random walk using the NetworkX package (version 2.6) in Python (version 3.8.18). We used *numpy.random.choices()* for weighted path selection, with transition probabilities proportional to exp(w/T), where w is the edge weight and T is a temperature parameter. For each node, 1000 random walks of length 20 steps are performed. Node importance is quantified by normalized visit frequency.

### Over-representation Analysis (ORA)

We performed ORA using the *enricher* function in the *clusterProfiler* package in R. For gene set definitions, we utilized the *msigdbr* package, which provides access to the Molecular Signatures Database (MSigDB) for Hallmark and C2 collection.

### Gene and annotation embeddings

We generated a summary paragraph for each gene and annotation. For genes, we used the NCBI summary. For MSigDB annotations, we looked for a full description (or a brief description, if a full description was not available) on the MSigDB webpage. For ICKG annotations, we created a prompt and asked Llama3-70B parameter model to generate a short summary paragraph for each term. We fed the summary paragraphs for genes and annotations into the pretrained BERT base model and obtained the first two-dimension embeddings and map genes/annotations onto a 2-dimensional map.

### Gaussian Mixture Model

We employed the Gaussian Mixture Model (GMM) to categorize within-cluster sum of squares (WCSS) data into two groups, aiming to identify clusters that indicate high and low levels of variance. We used the *GaussianMixture* function from the *sklearn.mixture* python library, specifying two components and a random seed for reproducibility *(n_components=2, random_state=0)*. After fitting GMM to the WCSS data using the *fit* method, we extracted the mean of each component from the model. The component with the higher mean was labeled as “High”, representing clusters with higher variance (lower specificity), whereas the component with the lower mean was labeled as “Low,” indicative of clusters with lower variance (higher specificity).

### Protein-protein interaction network

We fetched protein-protein interaction (PPI) data from the STRING database for each gene set. This interaction data was then employed to construct functional gene interaction networks using the *NetworkX* library. Density reflects how tightly nodes are linked. Clustering coefficient reveals local connectivity. Modularity evaluates how well a network divides into tightly-knit communities with sparse interconnections, shedding light on the network’s overall structure. Specifically, we utilized the *nx.Graph()* function to create the graph structure, populating it with nodes and weighted edges based on interaction scores. The constructed networks were analyzed to calculate key structural metrics: network density, clustering coefficient, and modularity. These metrics were computed with *nx.density(), nx.average_clustering(),* and *nx.algorithms.community.modularity()*.

### Differential gene expression

Differential expression gene (DEG) analysis was performed using R (version 4.2.3) and the Seurat package (version 4.1.4). The preprocessed and normalized single-cell RNA sequencing data were analyzed using the *FindAllMarkers* function from Seurat with the default parameter settings. Genes were considered DEGs if their adjusted p-value < 0.05. The p-values were adjusted for multiple testing using the Bonferroni correction method.

### Llama3-70B Prompt

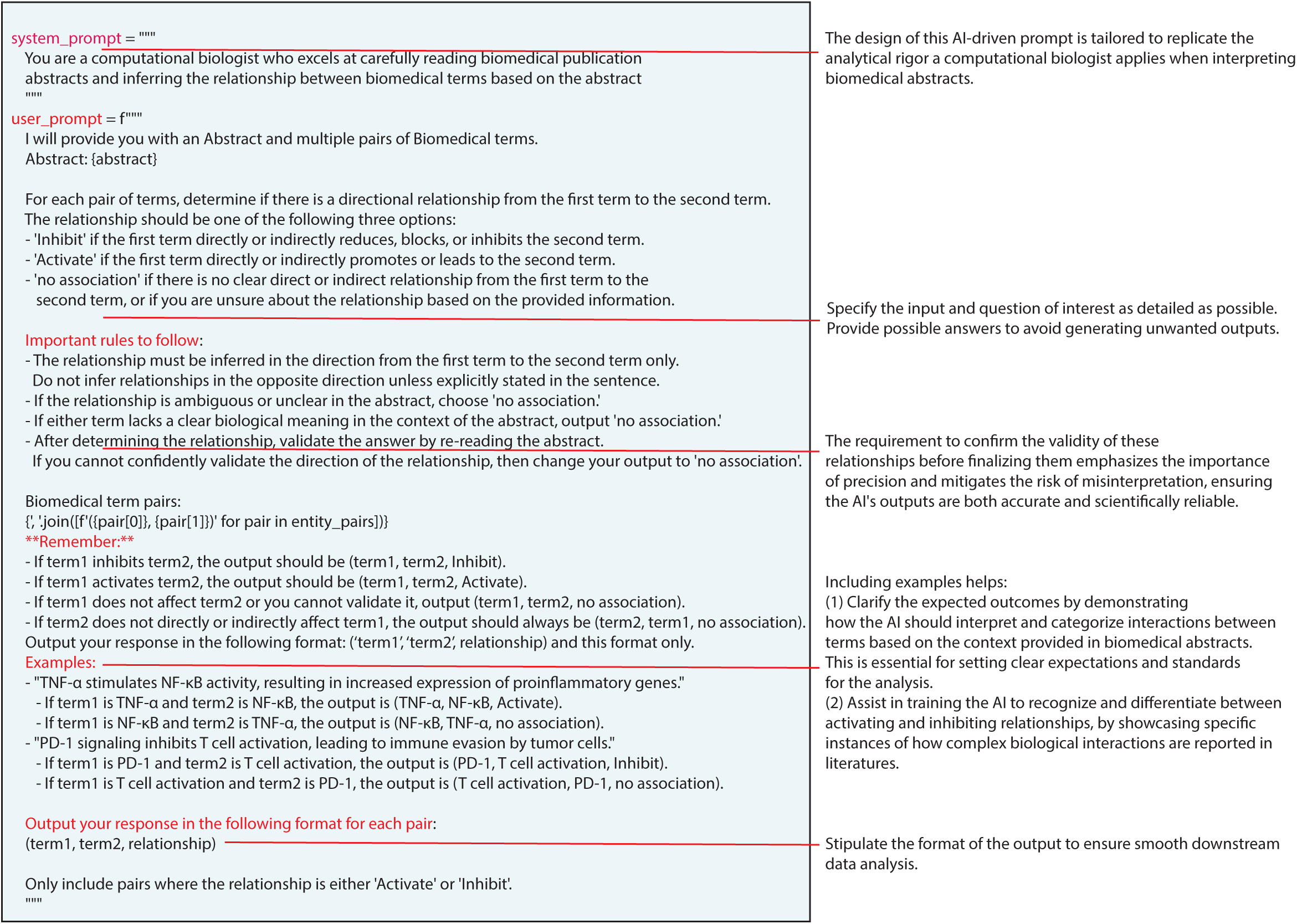

## Data and Code availability

All codes have been made available in https://github.com/KChen-lab/ICKG/tree/main/ including extraction of PubMed abstracts, biomedical entities recognition, knowledge graphs construction, and gene set annotations. In addition, finetuned NER models for genes, diseases and cell types and the manually annotated pathway database based on GENIA database have also been uploaded on https://github.com/KChen-lab/ICKG/tree/main/finetuned_NER_models. To facilitate user-friendly application, we have created a website for immune cell type specific gene set annotations: https://kchen-lab.github.io/immune-knowledgegraph.github.io/

## Supporting information

Main Tables

## Acknowledgements

This work is made possible by 2024-345892 to K.C. from the Chan Zuckerberg Initiative DAF, an advised fund of the Chan Zuckerberg Initiative Foundation, 5U01CA281902 from National Cancer Institute and The University of Texas MD Anderson Cancer Center Institute for Cell Therapy Discovery & Innovation. M.M.G. is a Cancer Prevention and Research Institute of Texas (CPRIT) Scholar in Cancer Research and is supported by CPRIT (Recruitment of First-Time Tenure-Track Faculty Members; RR190017), an Andrew Sabin Family Foundation Fellowship, and NCI R01CA282027 to M.M.G. We sincerely thank Jessica Swanton for proofreading the study. We sincerely thank Drs. Traver Hart and Christine Peterson for their feedback during this study.

