## Supplementary figures and images for "Literature-scaled immunological gene set annotation using AI-powered immune cell knowledge graph (ICKG)"

### Supplemental Figure 1

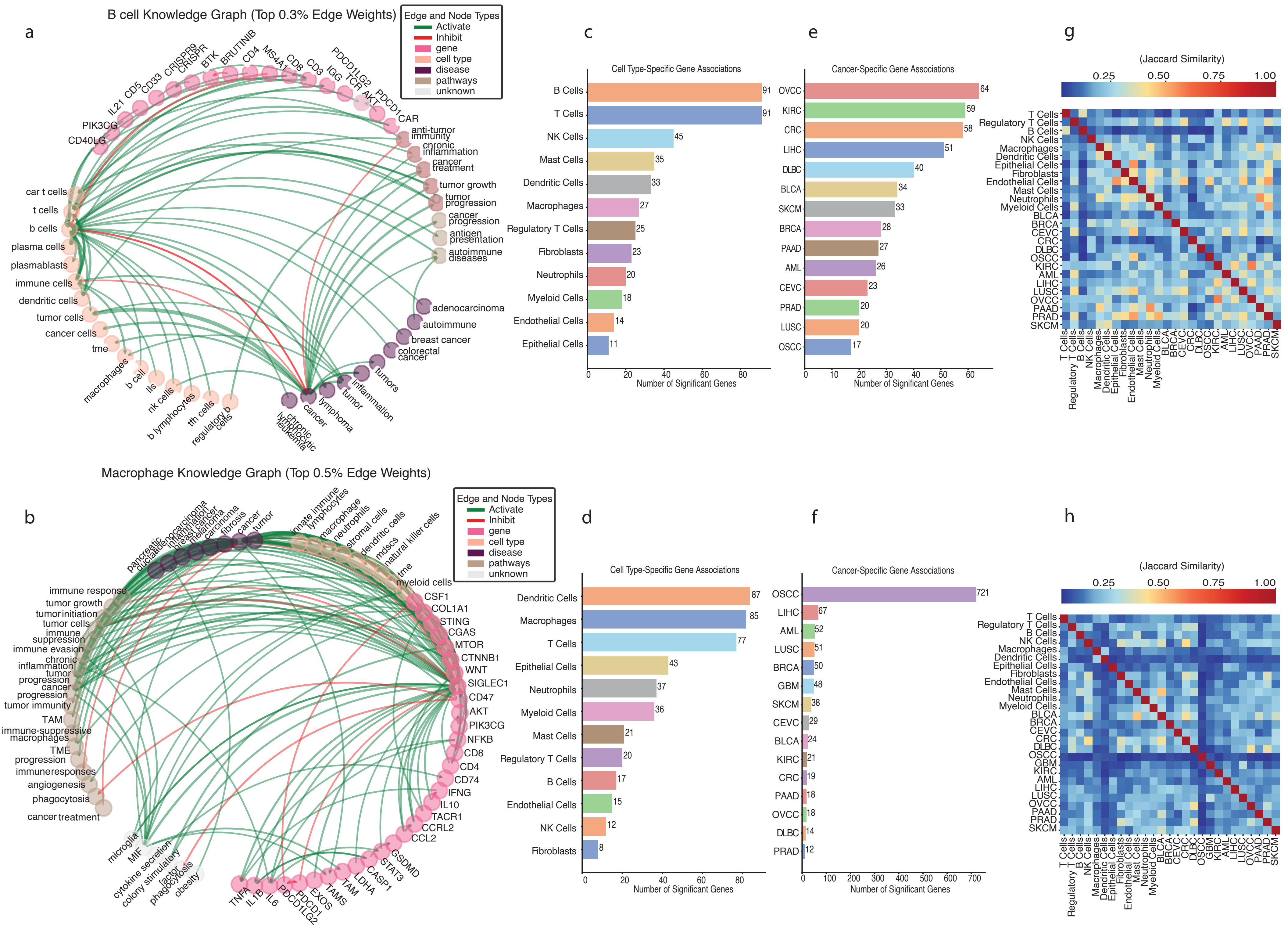
