## Supplemental Figure 2 for "Literature-scaled immunological gene set annotation using AI-powered immune cell knowledge graph (ICKG)"

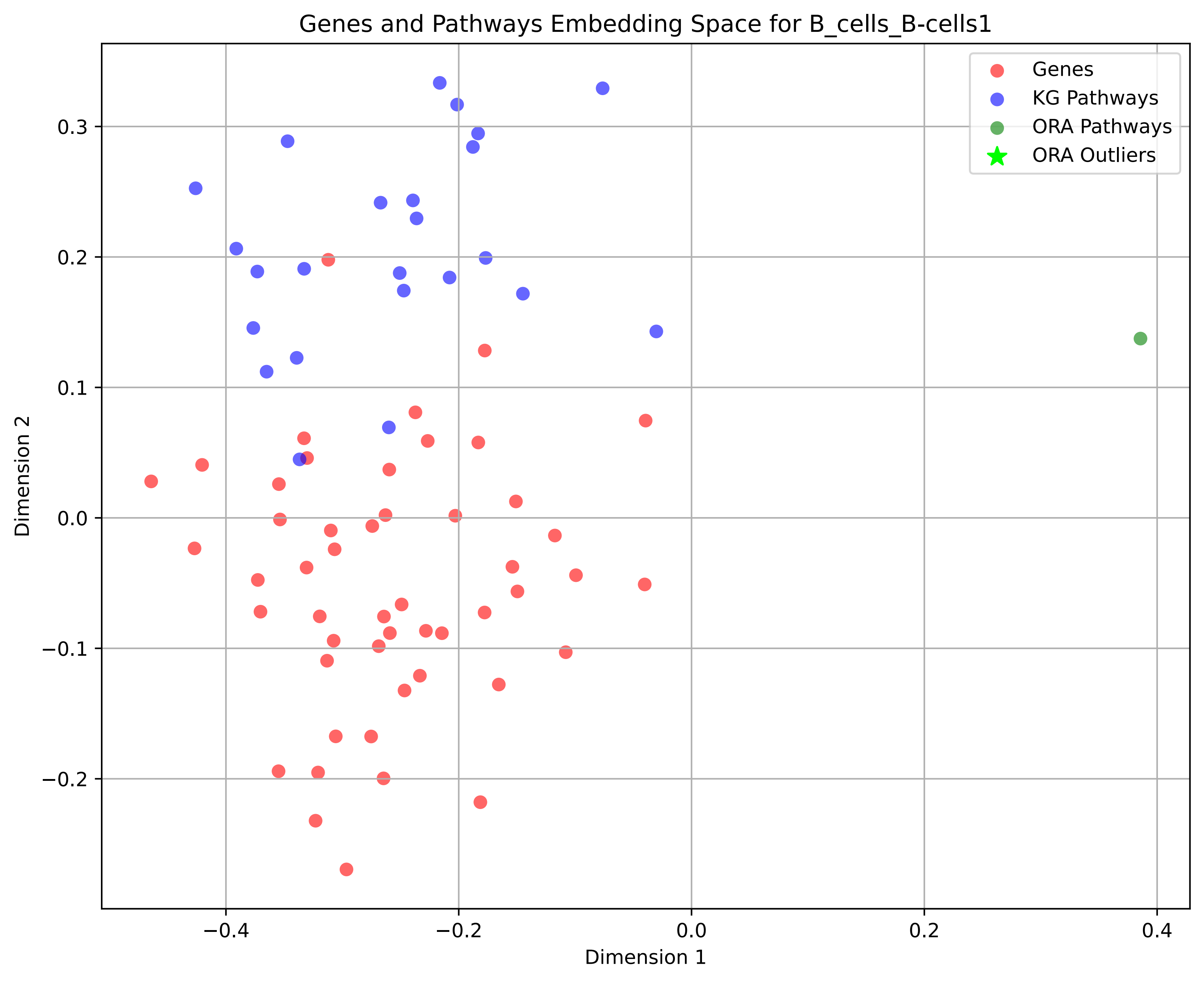
Genes and Pathways Embedding Space for B_cells_B-cells1
Genes
KG Pathways
ORA Pathways
ORA Outliers
Dimension 1
Dimension 2
−0.4
−0.2
0.0
0.2
0.4
−0.2
−0.1
0.0
0.1
0.2
0.3

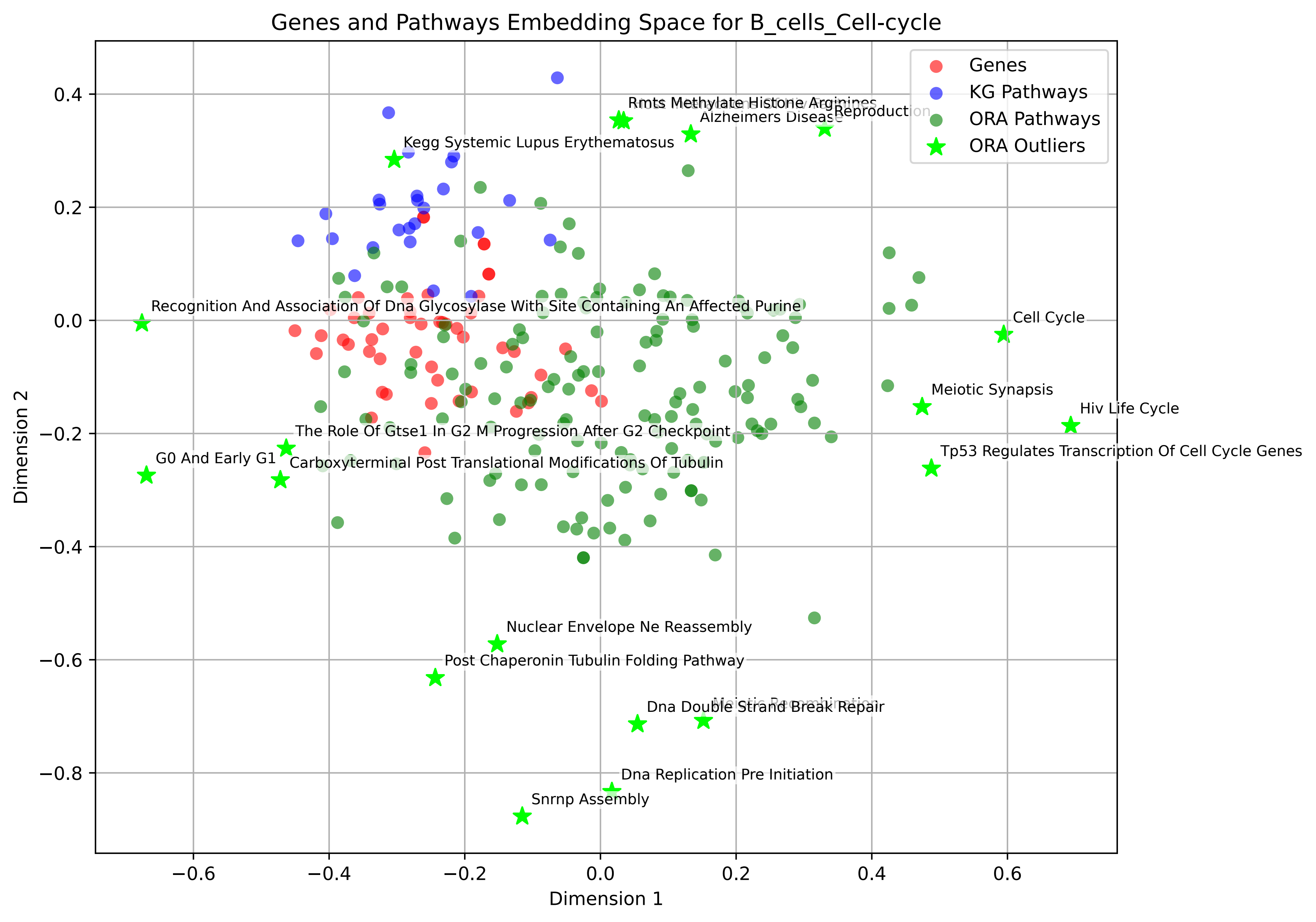

Genes and Pathways Embedding Space for B_cells_Cell-cycle
Genes
KG Pathways
ORA Pathways
ORA Outliers
Rmts Methylate Histone Arginines
Alzheimers Disease
Reproduction
Kegg Systemic Lupus Erythematosus
Recognition And Association Of Dna Glycosylase With Site Containing An Affected Purine
Cell Cycle
Meiotic Synapsis
Hiv Life Cycle
Tp53 Regulates Transcription Of Cell Cycle Genes
The Role Of Gtse1 In G2 M Progression After G2 Checkpoint
G0 And Early G1
Carboxyterminal Post Translational Modifications Of Tubulin
Nuclear Envelope Ne Reassembly
Post Chaperonin Tubulin Folding Pathway
Meiotic Recombination
Dna Double Strand Break Repair
Dna Replication Pre Initiation
Snrnp Assembly
Dimension 1
Dimension 2
0.4
0.2
0.0
−0.2
−0.4
−0.6
−0.8
−0.6
−0.4
−0.2
0.0
0.2
0.4
0.6

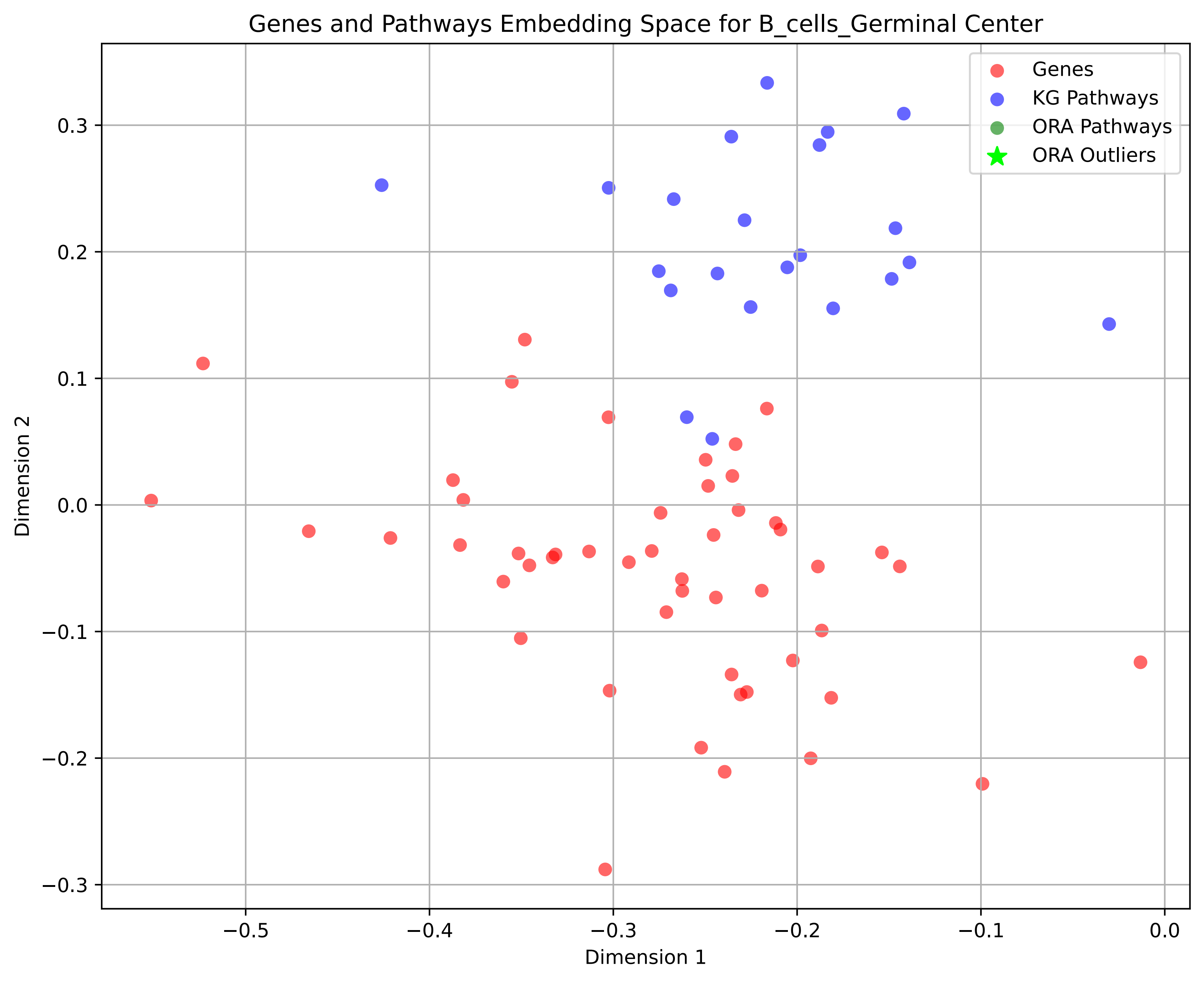
Genes and Pathways Embedding Space for B_cells_Germinal Center
Genes
KG Pathways
ORA Pathways
ORA Outliers
Dimension 1
Dimension 2
0.3
0.2
0.1
0.0
−0.1
−0.2
−0.3
−0.5
−0.4
−0.3
−0.2
−0.1
0.0

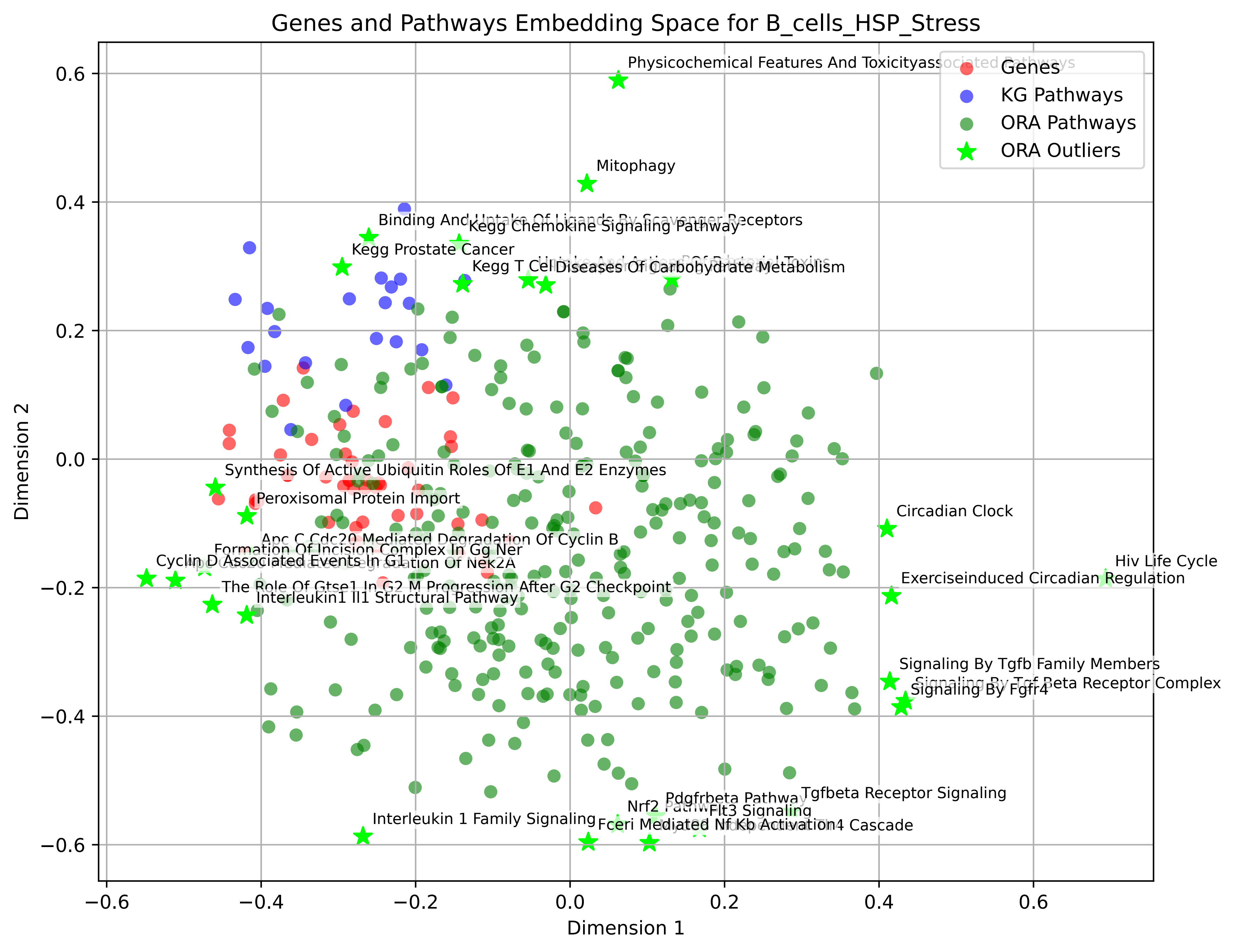

Genes and Pathways Embedding Space for B_cells_HSP_Stress
Genes
KG Pathways
ORA Pathways
ORA Outliers
Dimension 1
Dimension 2
0.6
0.4
0.2
0.0
−0.2
−0.4
−0.6
Physicochemical Features And Toxicityassociated Pathways
Mitophagy
Binding And Uptake Of Ligands By Scavenger Receptors
Kegg Chemokine Signaling Pathway
Kegg Prostate Cancer
Diseases Of Carbohydrate Metabolism
Synthesis Of Active Ubiquitin Roles Of E1 And E2 Enzymes
Peroxisomal Protein Import
Circadian Clock
Apc C Cdc20 Mediated Degradation Of Cyclin B
Formation Of Incision Complex In Gg Ner
Cyclin D Associated Events In G1
Hiv Life Cycle
Exerciseinduced Circadian Regulation
The Role Of Gtse1 In G2 M Progression After G2 Checkpoint
Interleukin1 Il1 Structural Pathway
Signaling By Tgfb Family Members
Signaling By Tgf Beta Receptor Complex
Signaling By Fgfr4
Tgfbeta Receptor Signaling
Pdgfrbeta Pathway
Nrf2
Flt3 Signaling
Interleukin 1 Family Signaling
Fceri Mediated Nf Kb Activation

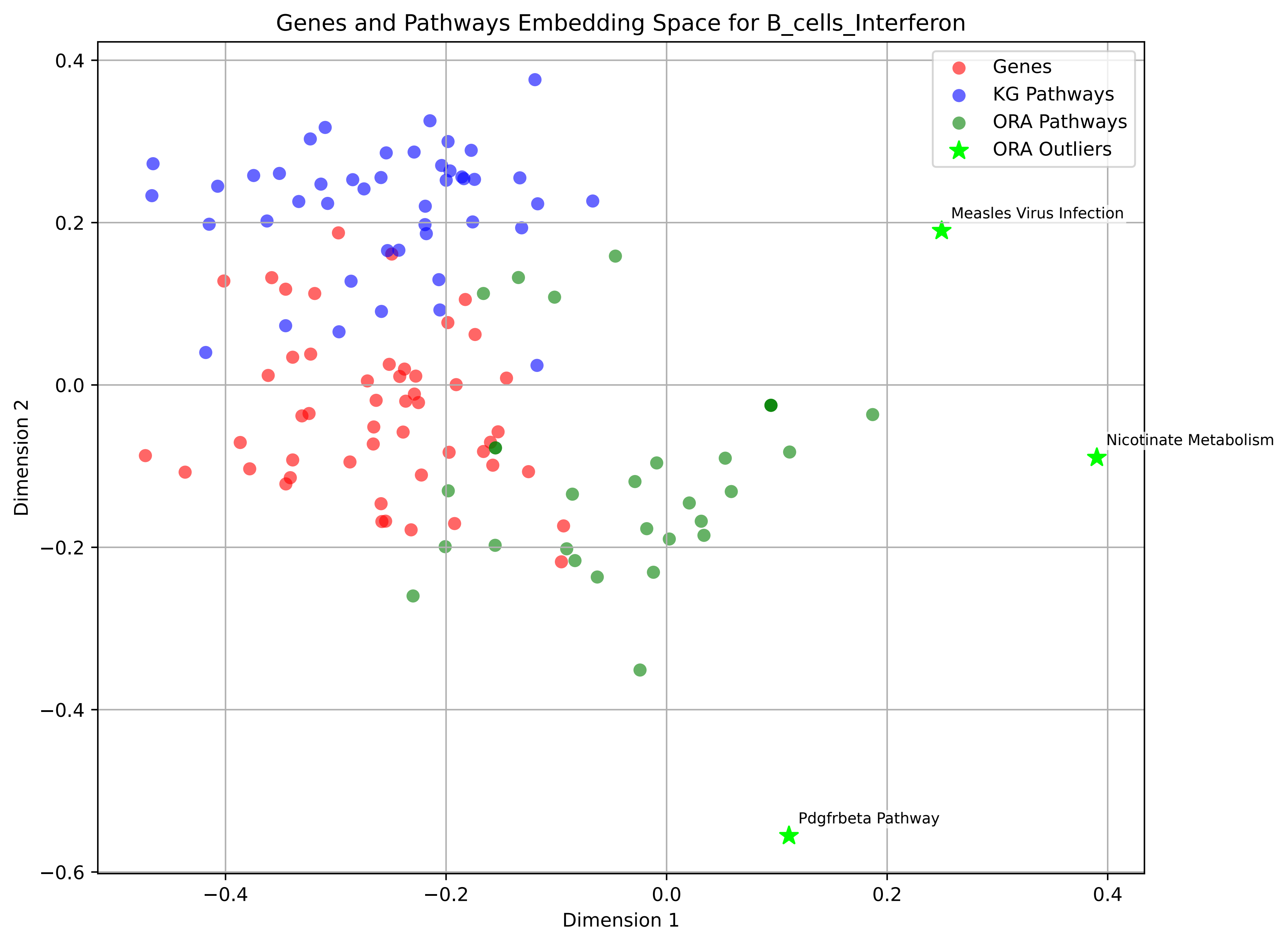
Genes and Pathways Embedding Space for B_cells_Interferon
Genes
KG Pathways
ORA Pathways
ORA Outliers
Measles Virus Infection
Nicotinate Metabolism
Pdgfrbeta Pathway
Dimension 1
Dimension 2
0.4
0.2
0.0
−0.2
−0.4
−0.6
−0.4
−0.2
0.0
0.2
0.4

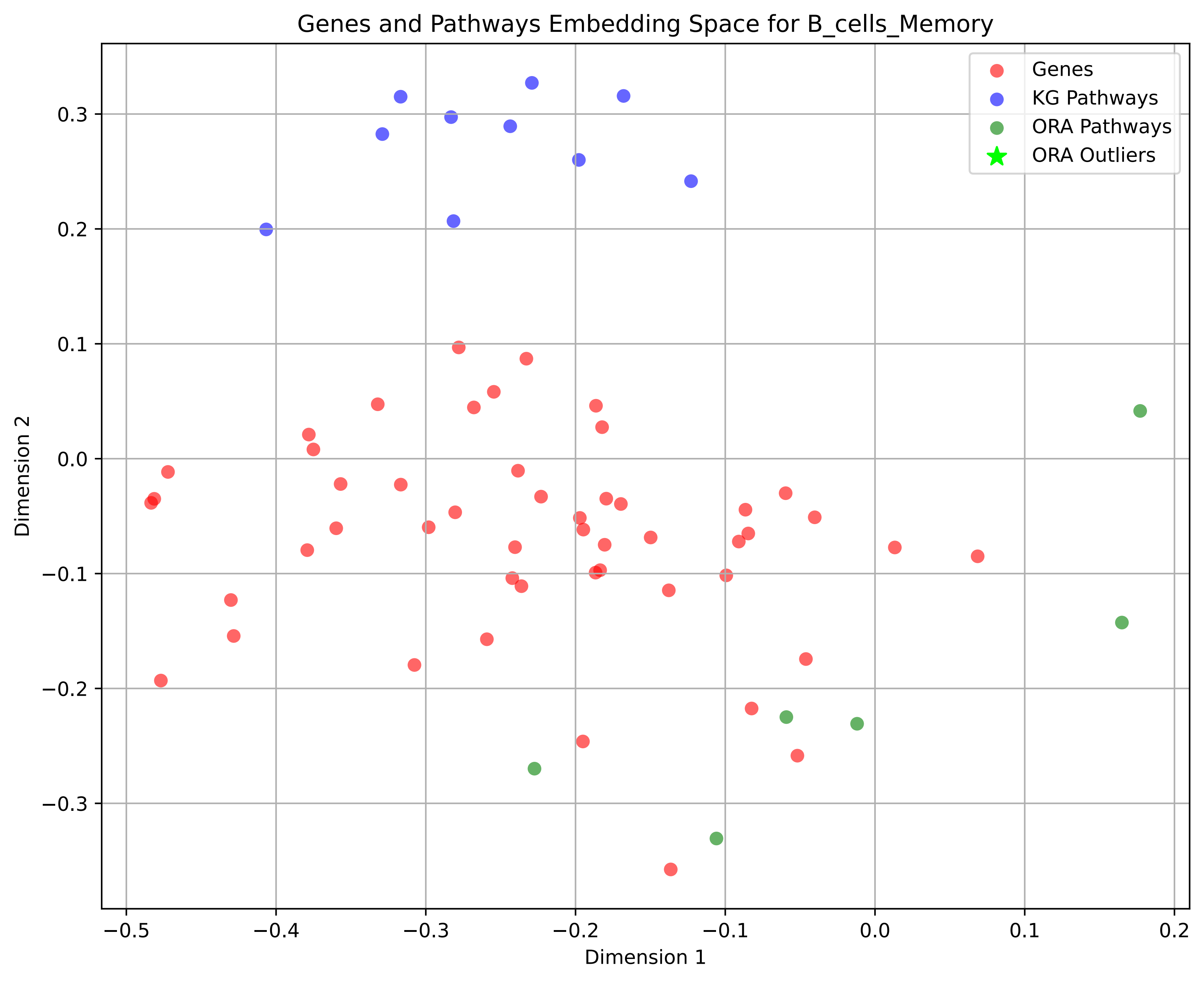
Genes and Pathways Embedding Space for B_cells_Memory
Genes
KG Pathways
ORA Pathways
ORA Outliers
Dimension 1
Dimension 2
0.3
0.2
0.1
0.0
−0.1
−0.2
−0.3
−0.5
−0.4
−0.3
−0.2
−0.1
0.0
0.1
0.2

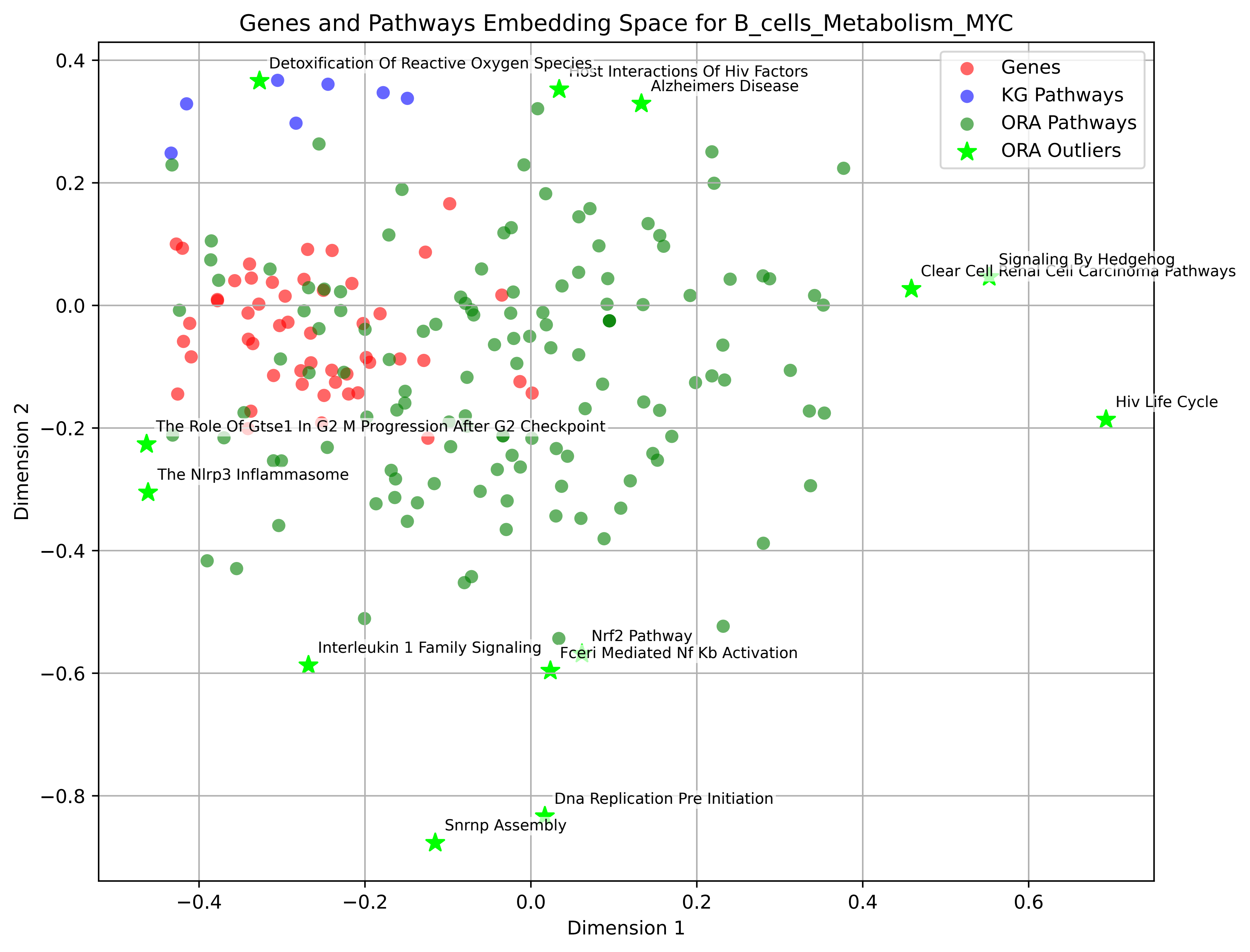
Genes and Pathways Embedding Space for B_cells_Metabolism_MYC
Genes
KG Pathways
ORA Pathways
ORA Outliers
Detoxification Of Reactive Oxygen Species
Host Interactions Of Hiv Factors
Alzheimers Disease
Signaling By Hedgehog
Clear Cell Renal Cell Carcinoma Pathways
Hiv Life Cycle
The Role Of Gtse1 In G2 M Progression After G2 Checkpoint
The Nlrp3 Inflammasome
Nrf2 Pathway
Interleukin 1 Family Signaling
Fceri Mediated Nf Kb Activation
Dna Replication Pre Initiation
Snrnp Assembly
Dimension 1
Dimension 2
0.4
0.2
0.0
−0.2
−0.4
−0.6
−0.8
−0.4
−0.2
0.0
0.2
0.4
0.6

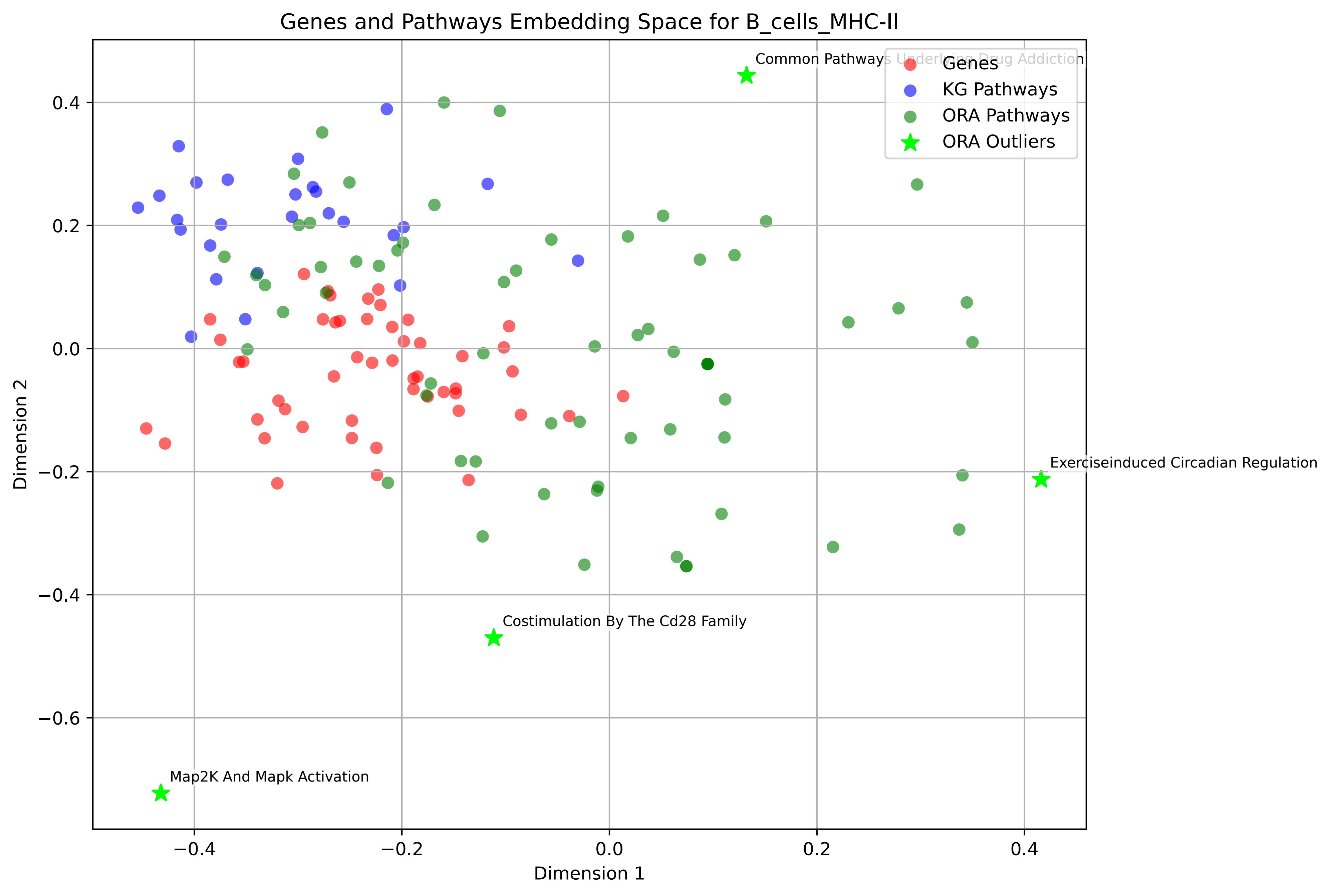

Genes and Pathways Embedding Space for B_cells_MHC-II
Genes
KG Pathways
ORA Pathways
ORA Outliers
Common Pathways Underlying Drug Addiction
Exerciseinduced Circadian Regulation
Costimulation By The Cd28 Family
Map2K And Mapk Activation
Dimension 1
Dimension 2
0.4
0.2
0.0
−0.2
−0.4
−0.6
−0.4
−0.2
0.0
0.2
0.4

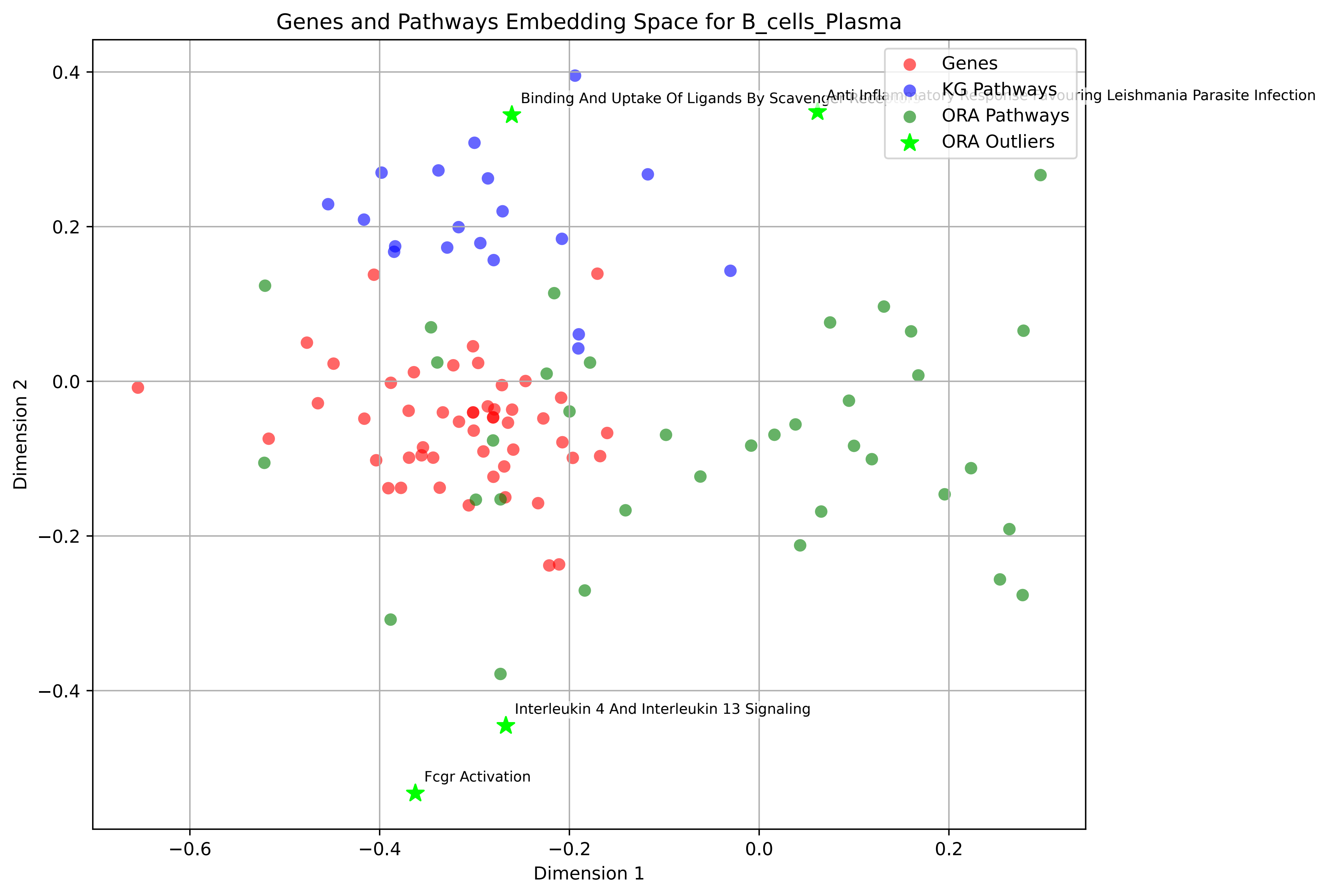

Genes and Pathways Embedding Space for B_cells_Plasma
Genes
KG Pathways
ORA Pathways
ORA Outliers
Binding And Uptake Of Ligands By Scavenger Receptors
Anti Inflammatory Response Favouring Leishmania Parasite Infection
Interleukin 4 And Interleukin 13 Signaling
Fcgr Activation
Dimension 1
Dimension 2
0.4
0.2
0.0
−0.2
−0.4
−0.6
−0.4
−0.2
0.0
0.2

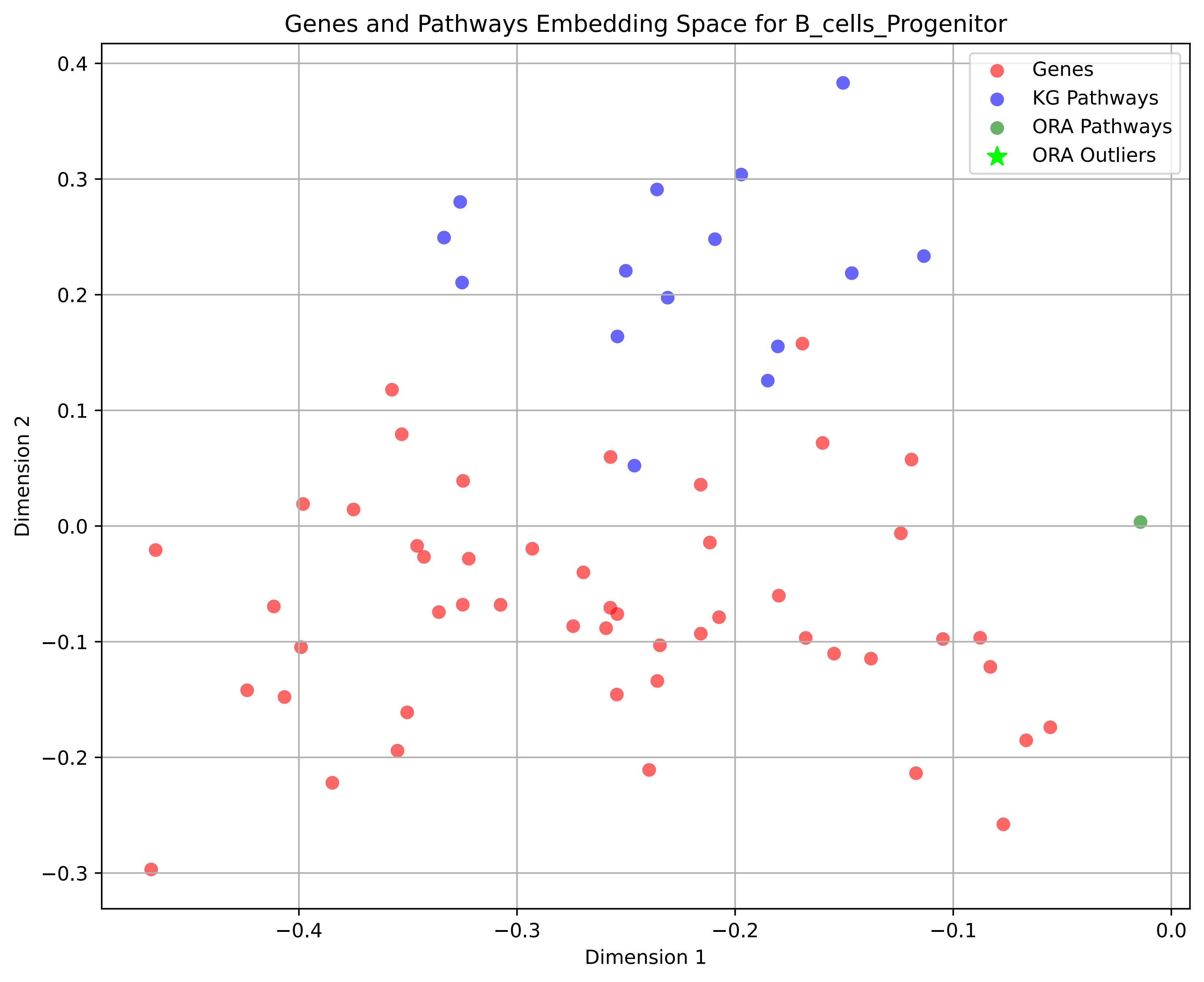

Genes and Pathways Embedding Space for B_cells_Progenitor
Genes
KG Pathways
ORA Pathways
ORA Outliers
Dimension 1
Dimension 2
0.4
0.3
0.2
0.1
0.0
−0.1
−0.2
−0.3
−0.4
−0.3
−0.2
−0.1
0.0

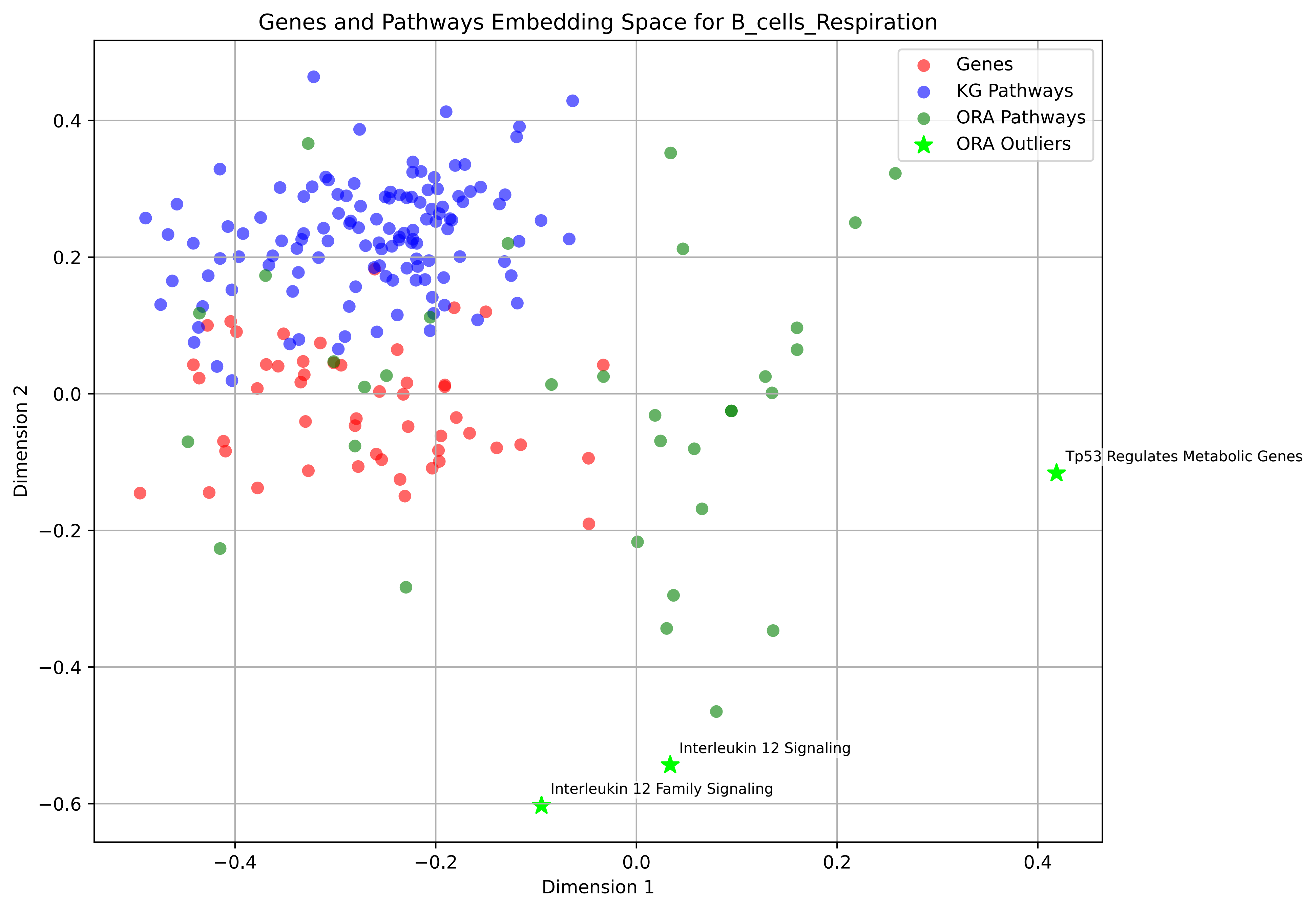

Genes and Pathways Embedding Space for B_cells_Respiration
Genes
KG Pathways
ORA Pathways
ORA Outliers
Tp53 Regulates Metabolic Genes
Interleukin 12 Signaling
Interleukin 12 Family Signaling
Dimension 1
Dimension 2
0.4
0.2
0.0
−0.2
−0.4
−0.6
0.4
0.2
0.0
−0.2
−0.4

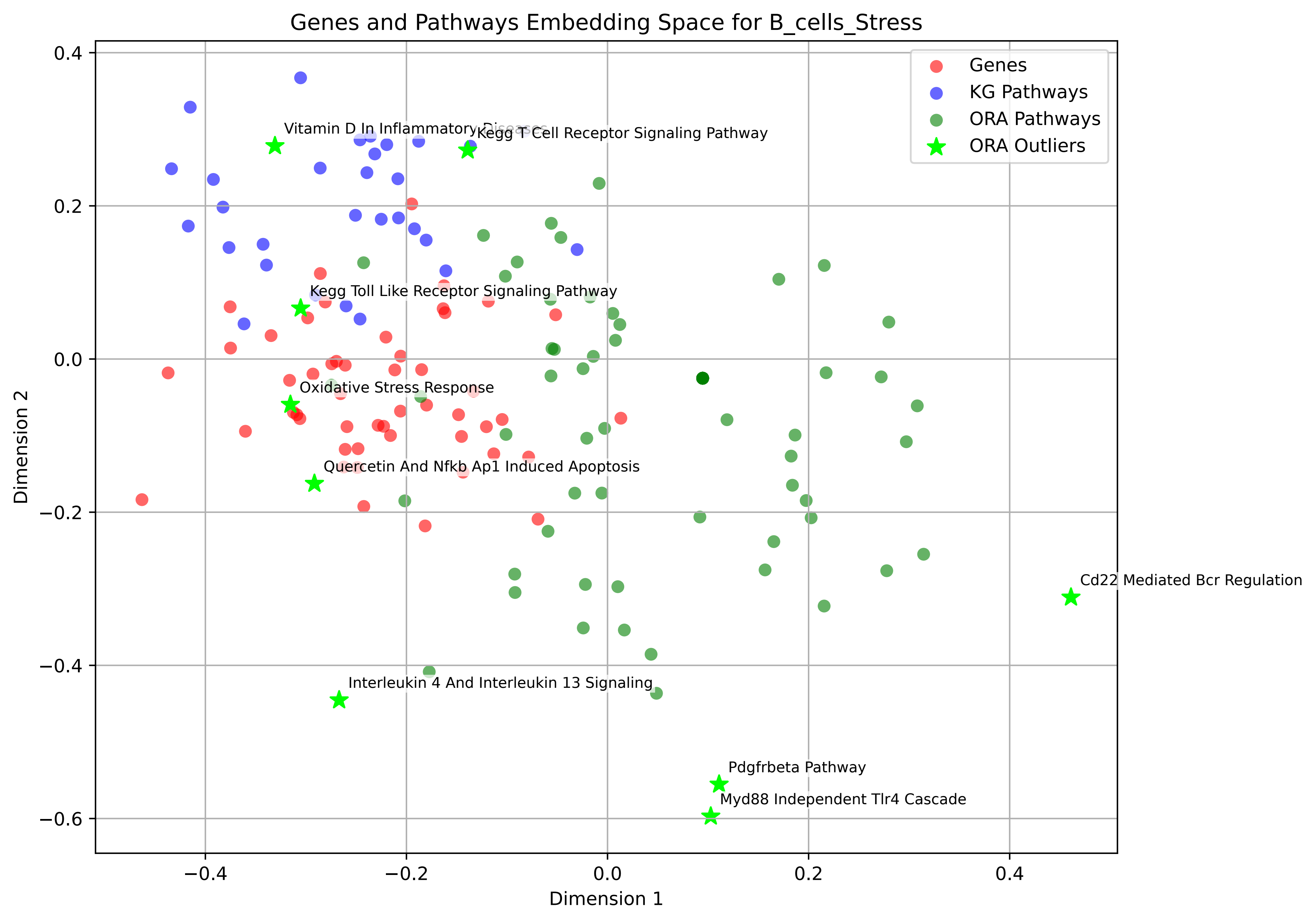
Genes and Pathways Embedding Space for B_cells_Stress
Genes
KG Pathways
ORA Pathways
ORA Outliers
Vitamin D In Inflammatory Diseases
Kegg T Cell Receptor Signaling Pathway
Kegg Toll Like Receptor Signaling Pathway
Oxidative Stress Response
Quercetin And Nfkb Ap1 Induced Apoptosis
Cd22 Mediated Bcr Regulation
Interleukin 4 And Interleukin 13 Signaling
Pdgfrbeta Pathway
Myd88 Independent Tlr4 Cascade
Dimension 1
Dimension 2
0.4
0.2
0.0
−0.2
−0.4
−0.6
−0.4
−0.2
0.0
0.2
0.4

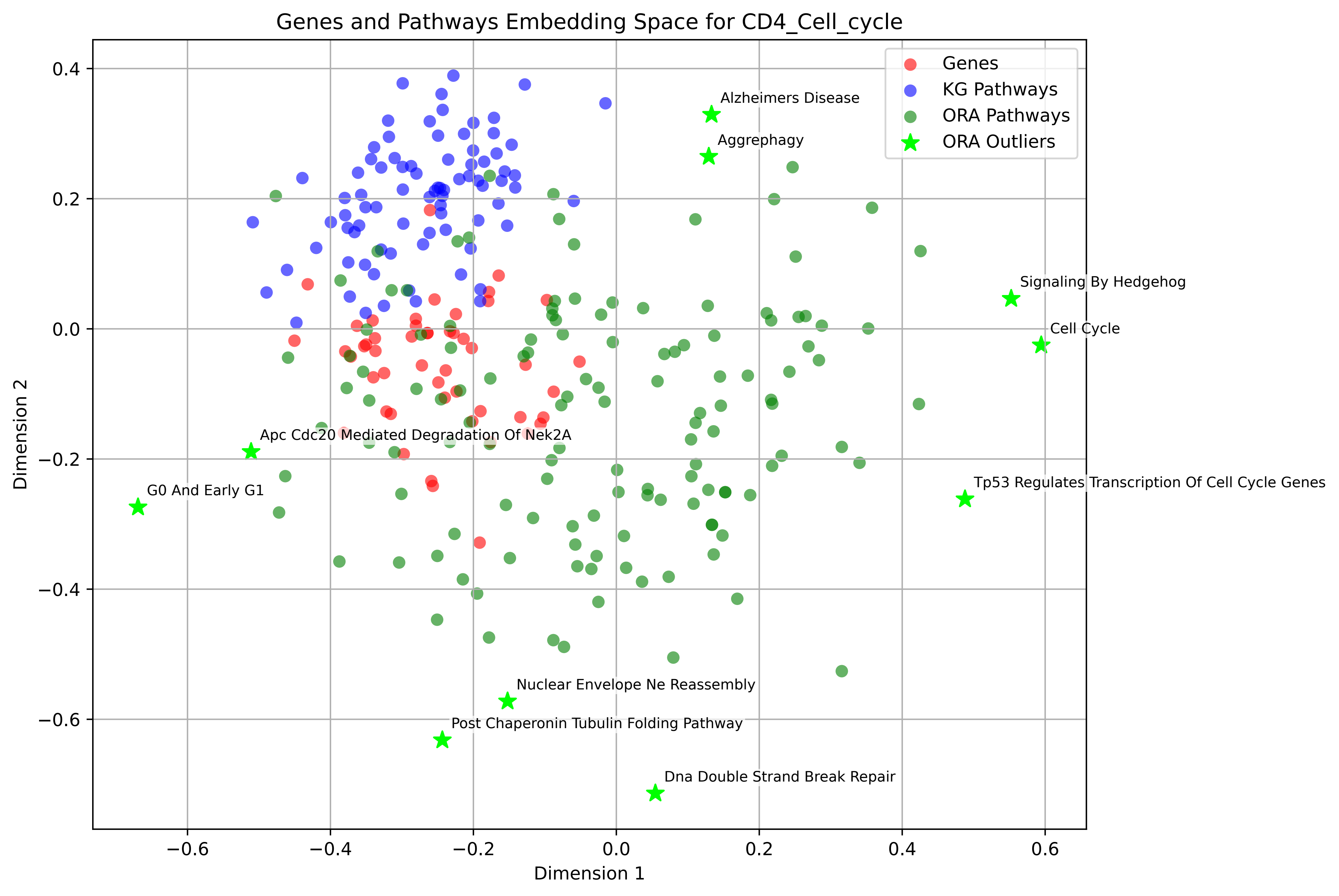

Genes and Pathways Embedding Space for CD4_Cell_cycle
Genes
KG Pathways
ORA Pathways
ORA Outliers
Alzheimers Disease
Aggrephagy
Signaling By Hedgehog
Cell Cycle
Tp53 Regulates Transcription Of Cell Cycle Genes
Apc Cdc20 Mediated Degradation Of Nek2A
G0 And Early G1
Nuclear Envelope Ne Reassembly
Post Chaperonin Tubulin Folding Pathway
Dna Double Strand Break Repair
Dimension 1
Dimension 2

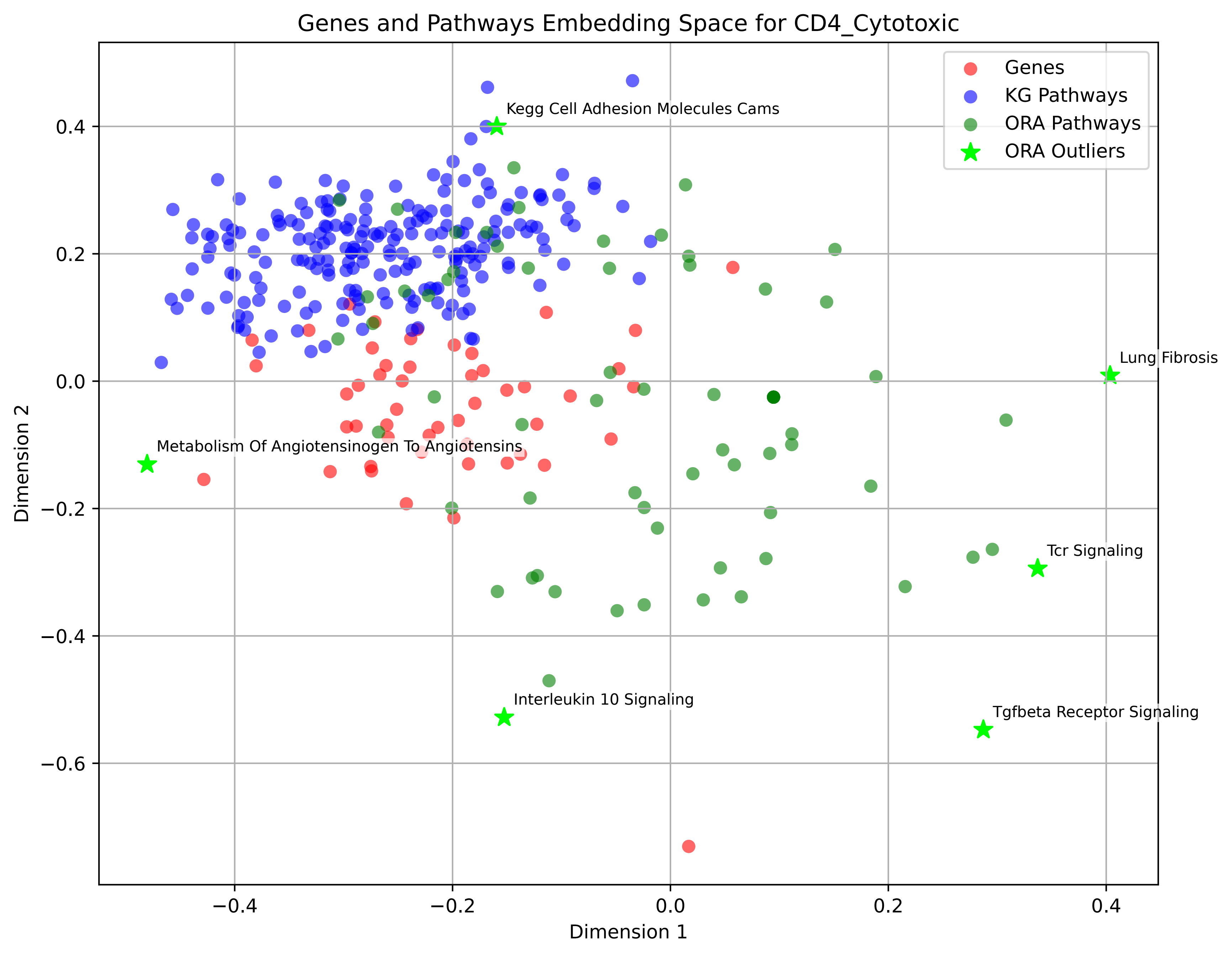

Genes and Pathways Embedding Space for CD4_Cytotoxic
Genes
KG Pathways
ORA Pathways
ORA Outliers
Kegg Cell Adhesion Molecules Cams
Lung Fibrosis
Metabolism Of Angiotensinogen To Angiotensins
Tcr Signaling
Interleukin 10 Signaling
Tgfbeta Receptor Signaling
Dimension 1
Dimension 2
0.4
0.2
0.0
−0.2
−0.4
−0.6
−0.4
−0.2
0.0
0.2
0.4

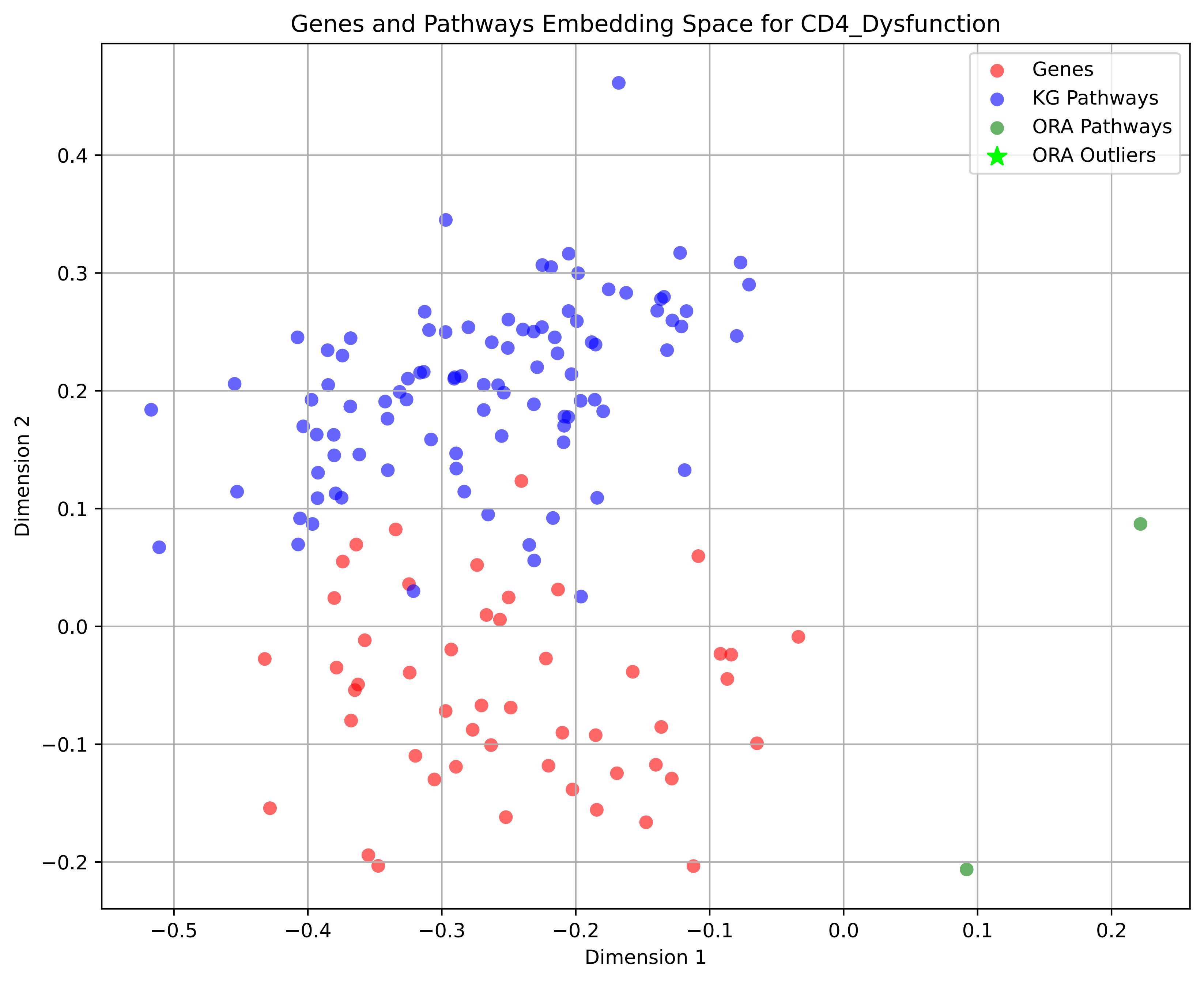

Genes and Pathways Embedding Space for CD4_Dysfunction
Genes
KG Pathways
ORA Pathways
ORA Outliers
Dimension 1
Dimension 2
−0.5
−0.4
−0.3
−0.2
−0.1
0.0
0.1
0.2
0.4
0.3
0.2
0.1
0.0
−0.1
−0.2

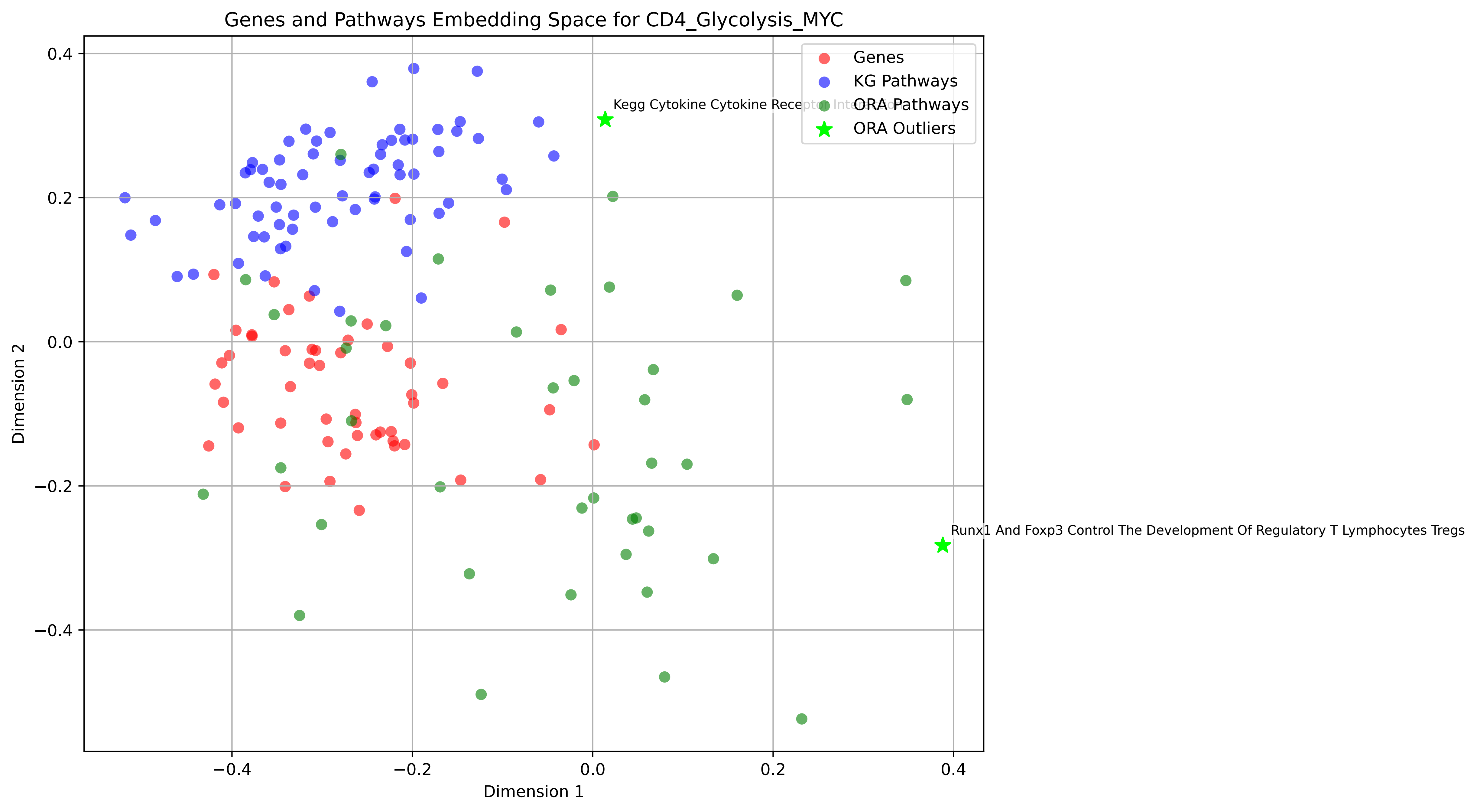

Genes and Pathways Embedding Space for CD4_Glycolysis_MYC
Genes
KG Pathways
ORA Pathways
ORA Outliers
Kegg Cytokine Cytokine Receptor Interaction
Runx1 And Foxp3 Control The Development Of Regulatory T Lymphocytes Tregs
Dimension 1
Dimension 2
0.4
0.2
0.0
−0.2
−0.4

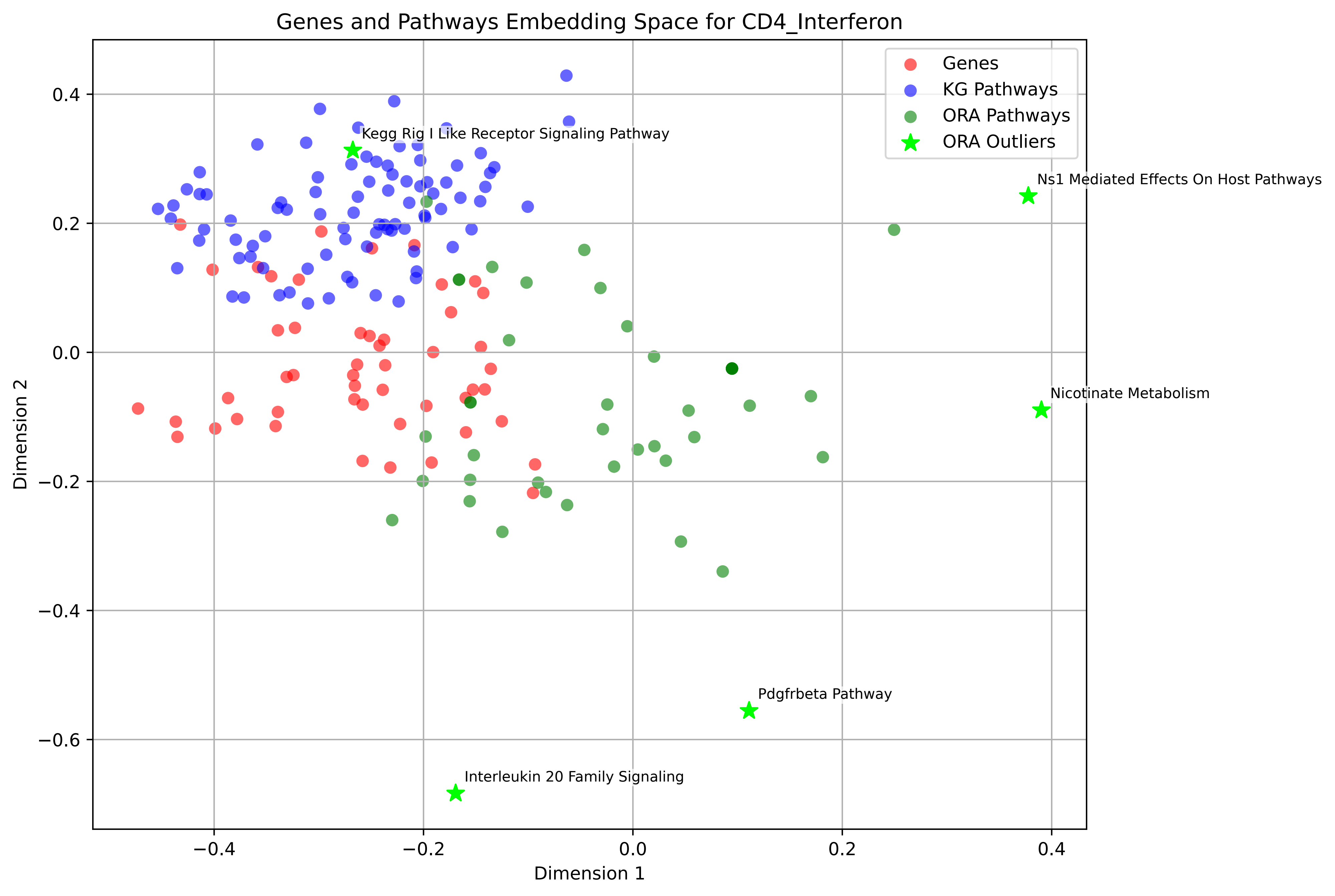

Genes and Pathways Embedding Space for CD4_Interferon
Genes
KG Pathways
ORA Pathways
ORA Outliers
Kegg Rig I Like Receptor Signaling Pathway
Ns1 Mediated Effects On Host Pathways
Nicotinate Metabolism
Pdgfrbeta Pathway
Interleukin 20 Family Signaling
Dimension 1
Dimension 2
0.4
0.2
0.0
−0.2
−0.4
−0.6
−0.4
−0.2
0.0
0.2
0.4

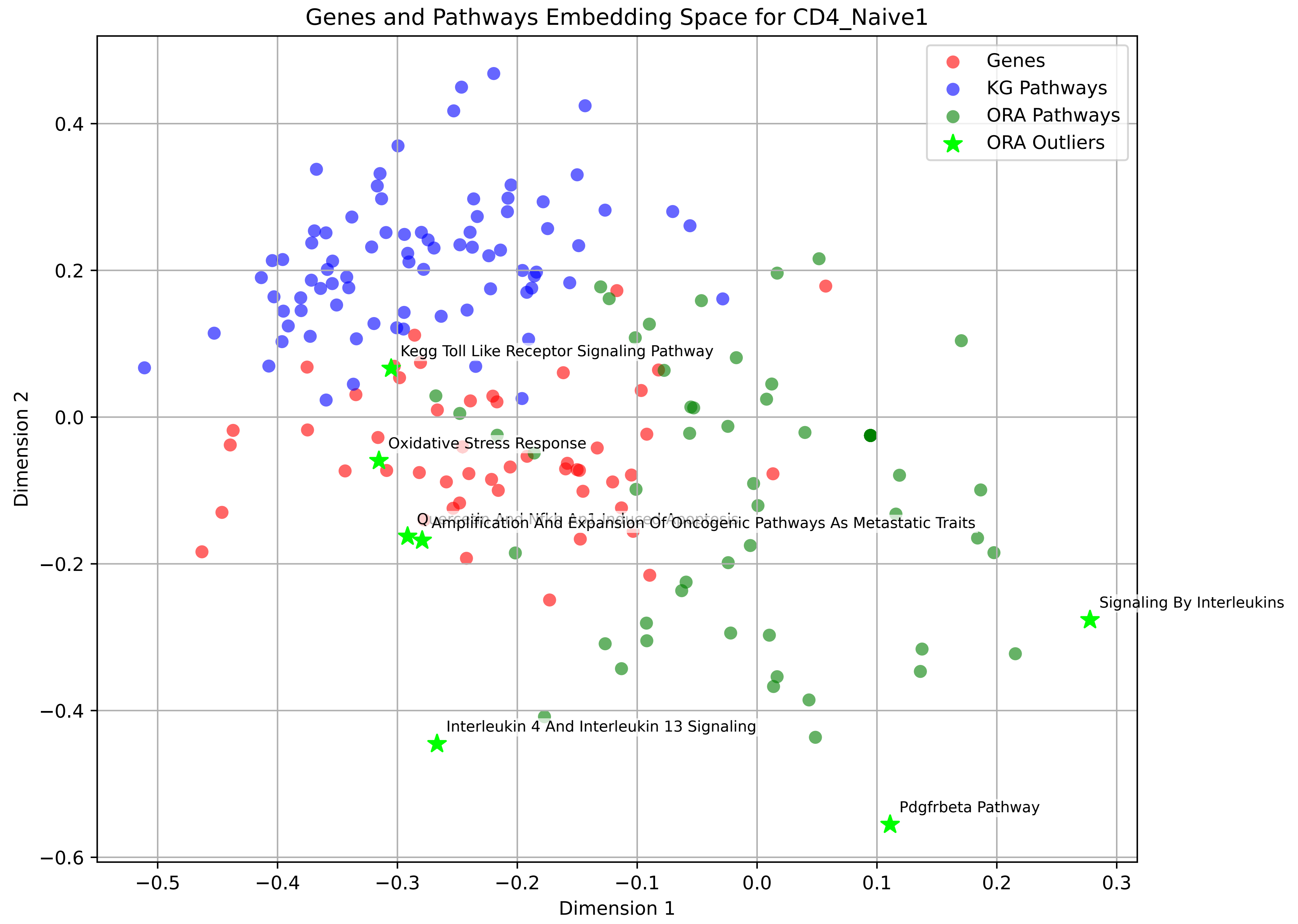

Genes and Pathways Embedding Space for CD4_Naive1
Genes
KG Pathways
ORA Pathways
ORA Outliers
Kegg Toll Like Receptor Signaling Pathway
Oxidative Stress Response
Quercetin And Nfkb Ap1 Induced Apoptosis
Amplification And Expansion Of Oncogenic Pathways As Metastatic Traits
Signaling By Interleukins
Interleukin 4 And Interleukin 13 Signaling
Pdgfrbeta Pathway
0.4
0.2
0.0
−0.2
−0.4
−0.6
−0.5
−0.4
−0.3
−0.2
−0.1
0.0
0.1
0.2
0.3
Dimension 1
Dimension 2

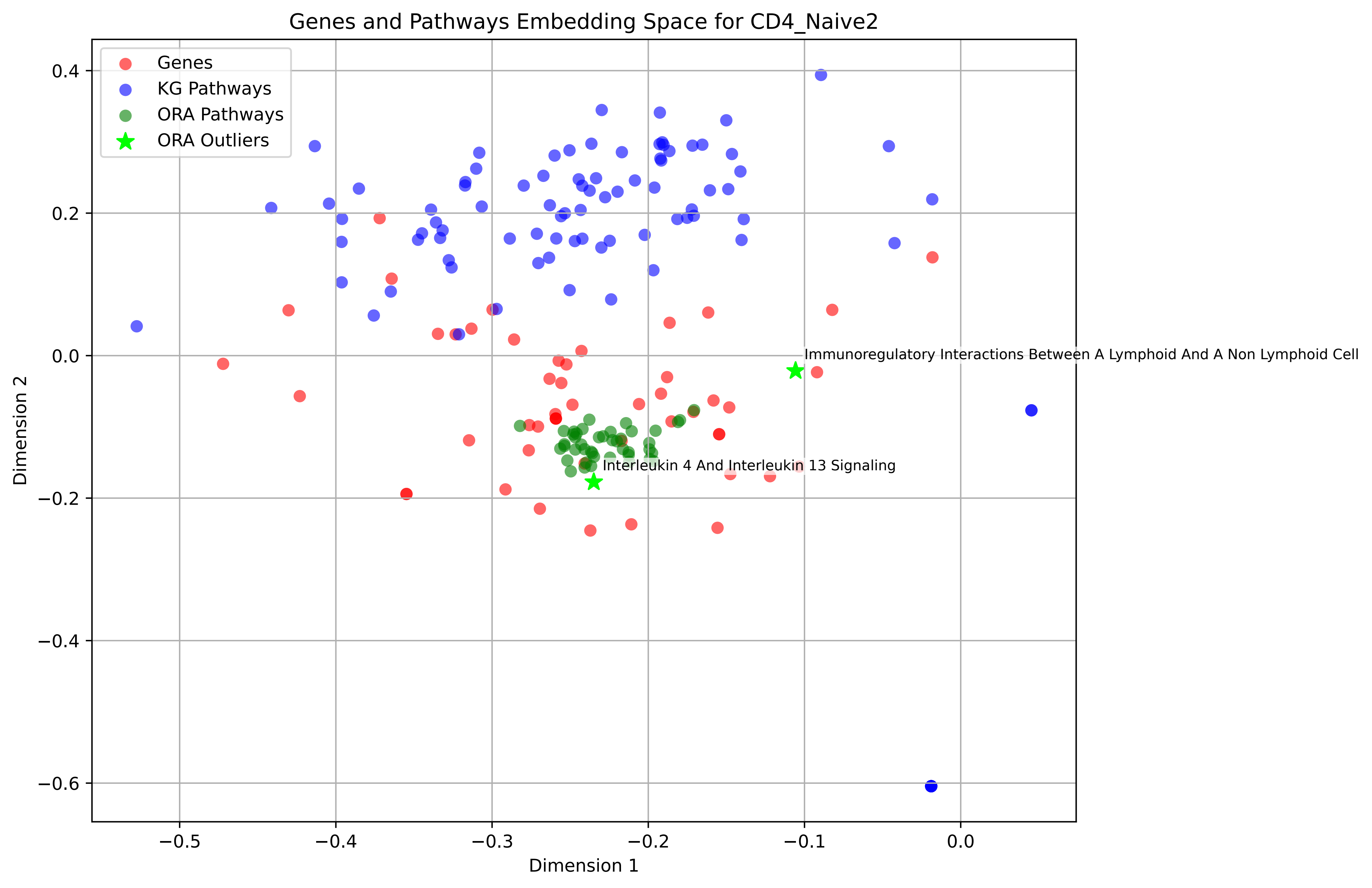

Genes and Pathways Embedding Space for CD4_Naive2
Genes
KG Pathways
ORA Pathways
ORA Outliers
Immunoregulatory Interactions Between A Lymphoid And A Non Lymphoid Cell
Interleukin 4 And Interleukin 13 Signaling
Dimension 1
Dimension 2
0.4
0.2
0.0
−0.2
−0.4
−0.6
−0.5
−0.4
−0.3
−0.2
−0.1
0.0

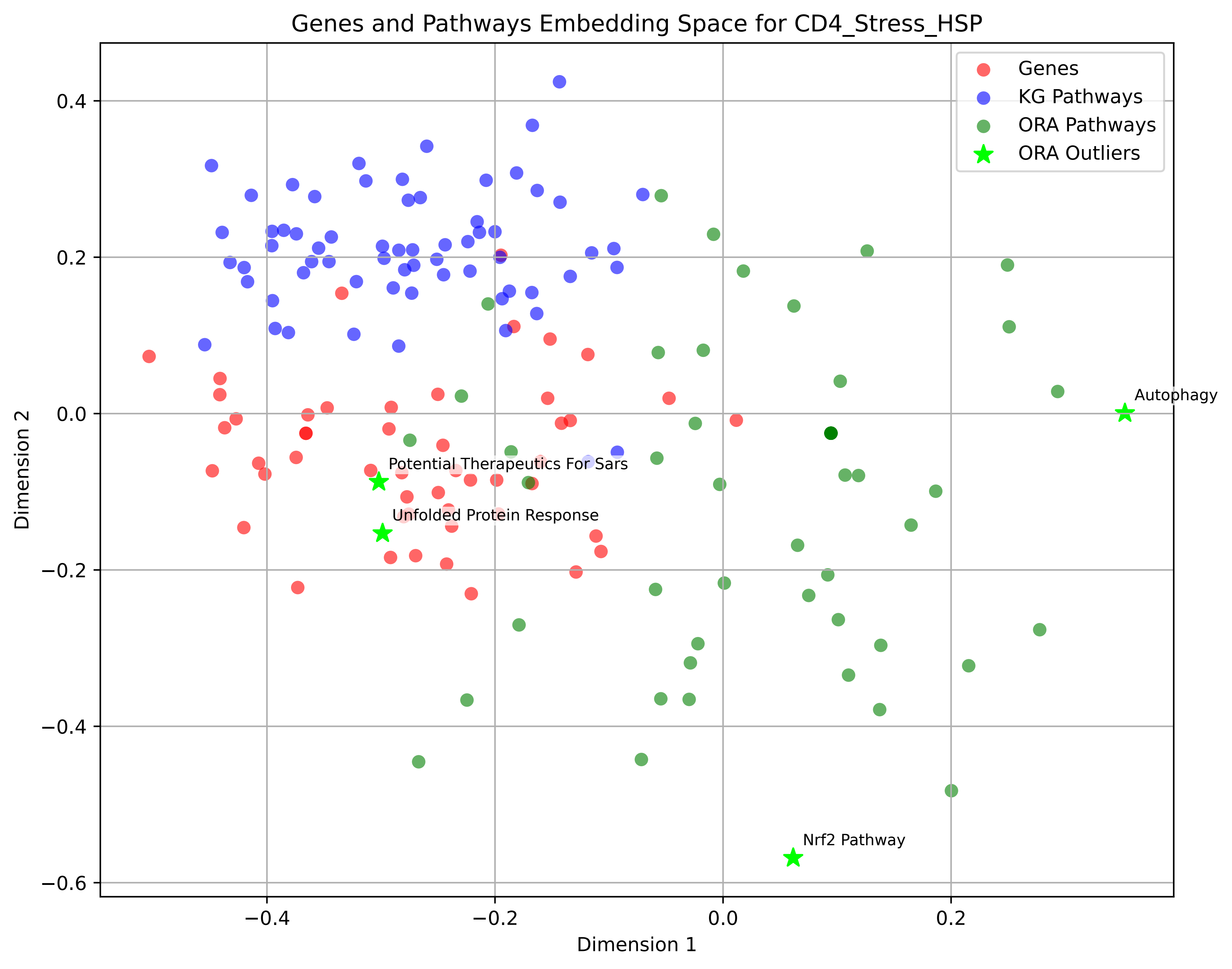
Genes and Pathways Embedding Space for CD4_Stress_HSP
Genes
KG Pathways
ORA Pathways
ORA Outliers
Autophagy
Potential Therapeutics For Sars
Unfolded Protein Response
Nrf2 Pathway
Dimension 1
Dimension 2
0.4
0.2
0.0
−0.2
−0.4
−0.6
−0.4
−0.2
0.0
0.2

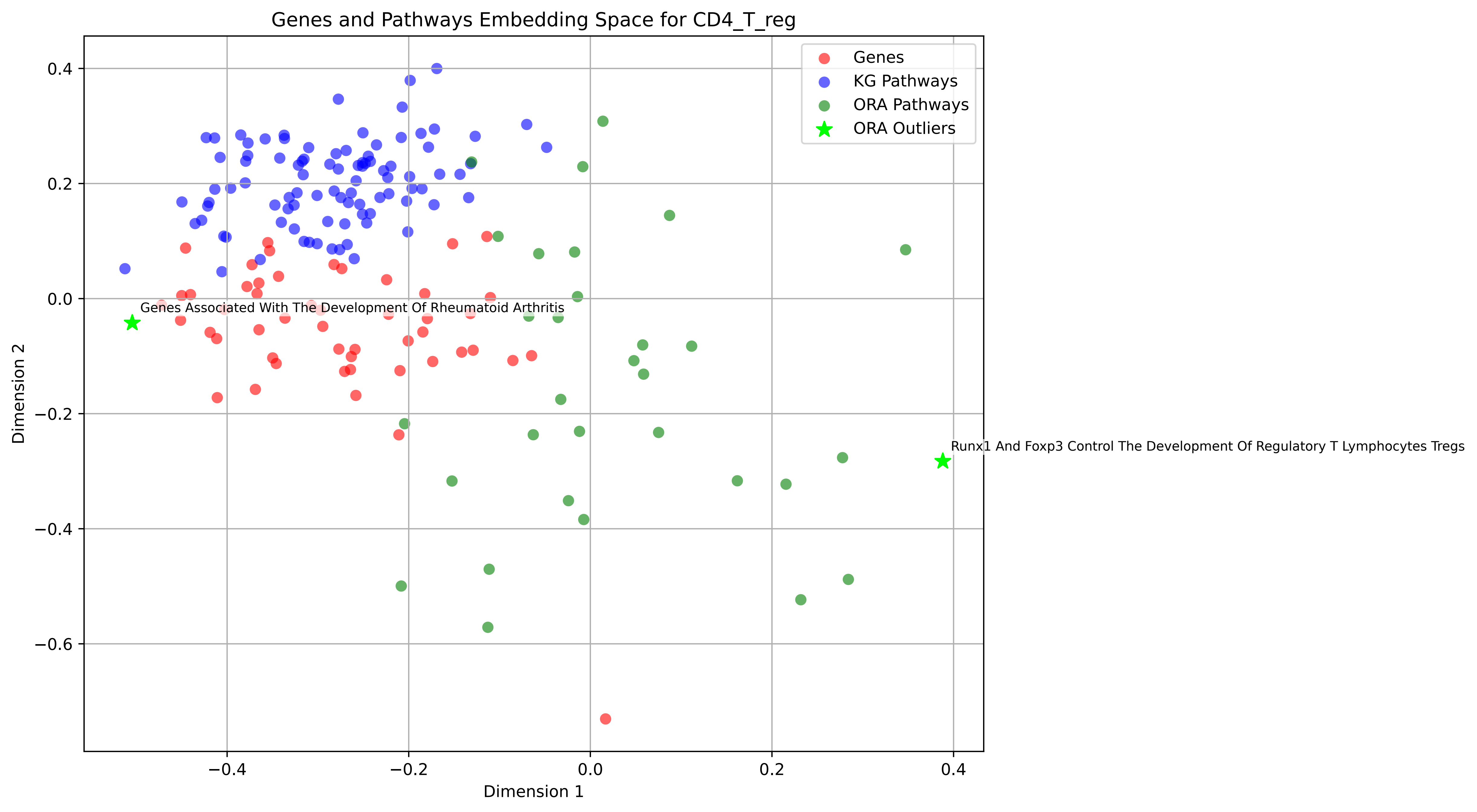
Genes and Pathways Embedding Space for CD4_T_reg
Genes
KG Pathways
ORA Pathways
ORA Outliers
Genes Associated With The Development Of Rheumatoid Arthritis
Runx1 And Foxp3 Control The Development Of Regulatory T Lymphocytes Tregs
Dimension 1
Dimension 2
0.4
0.2
0.0
−0.2
−0.4
−0.6
−0.4
−0.2
0.0
0.2
0.4

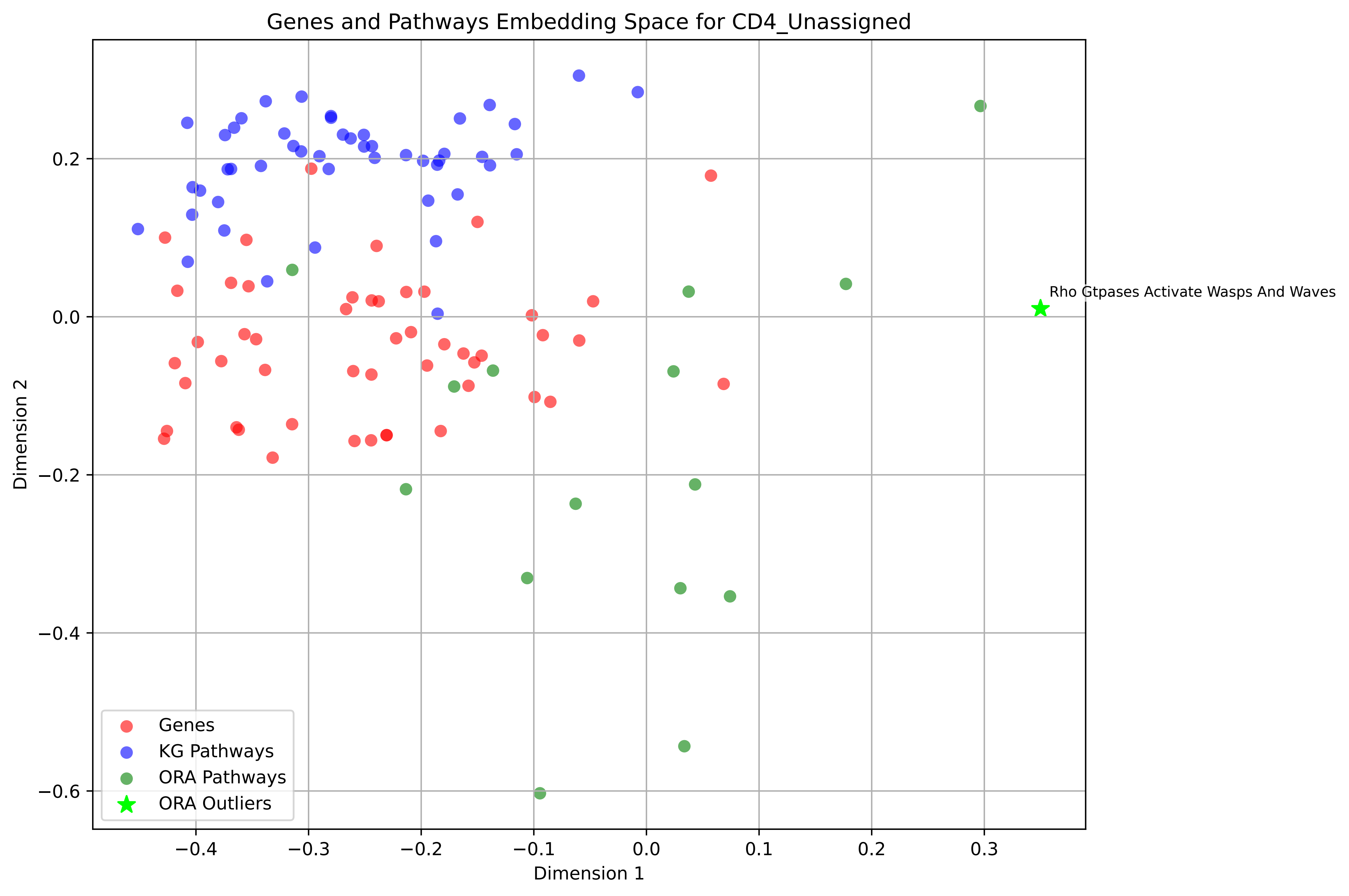

Genes and Pathways Embedding Space for CD4_Unassigned
Rho Gtpases Activate Wasps And Waves
Genes
KG Pathways
ORA Pathways
ORA Outliers
Dimension 1
Dimension 2
−0.4
−0.3
−0.2
−0.1
0.0
0.1
0.2
0.3
0.2
0.0
−0.2
−0.4
−0.6

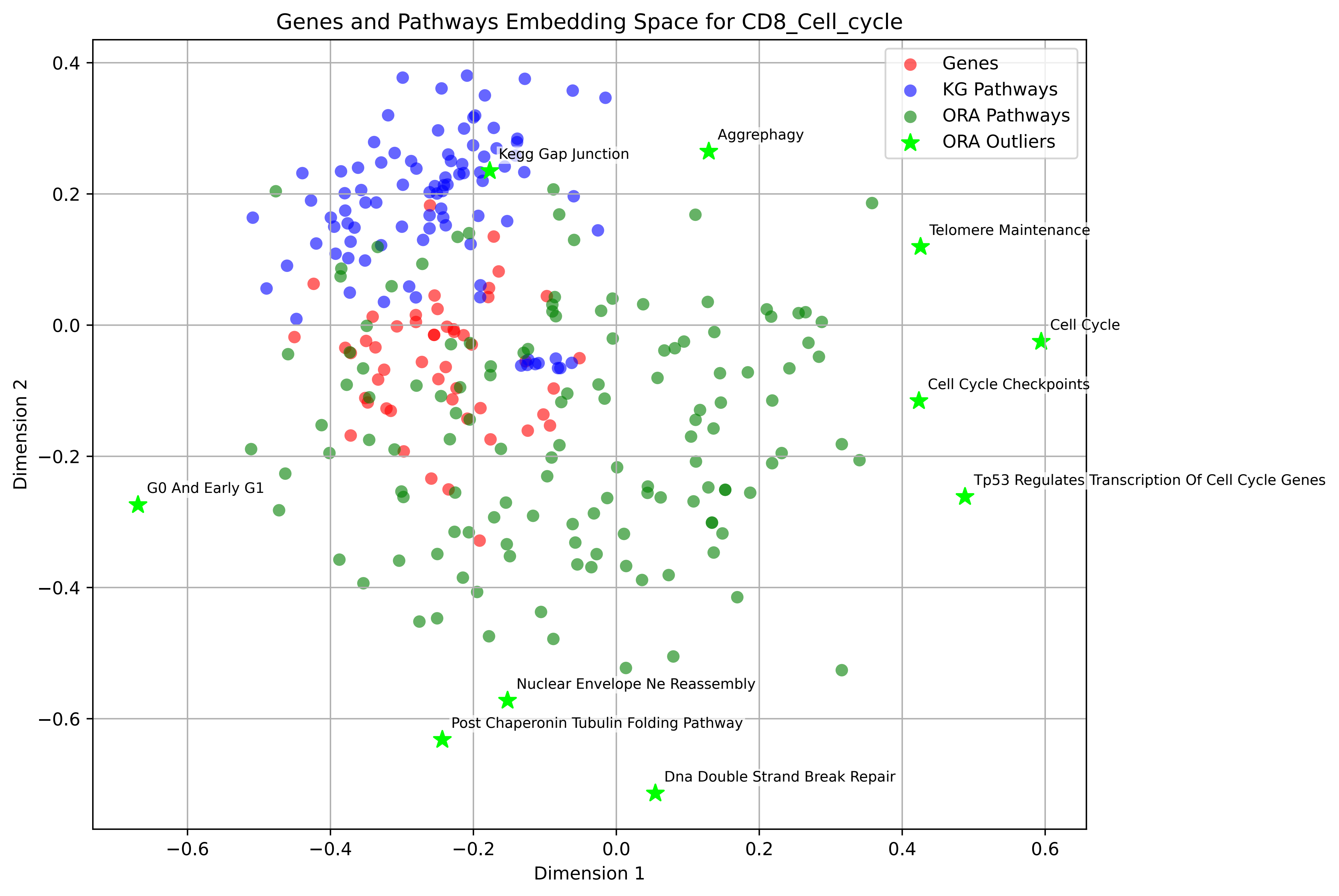
Genes and Pathways Embedding Space for CD8_Cell_cycle
Genes
KG Pathways
ORA Pathways
ORA Outliers
Aggrephagy
Kegg Gap Junction
Telomere Maintenance
Cell Cycle
Cell Cycle Checkpoints
Tp53 Regulates Transcription Of Cell Cycle Genes
G0 And Early G1
Nuclear Envelope Ne Reassembly
Post Chaperonin Tubulin Folding Pathway
Dna Double Strand Break Repair
Dimension 1
Dimension 2
0.4
0.2
0.0
−0.2
−0.4
−0.6
−0.6
−0.4
−0.2
0.0
0.2
0.4
0.6

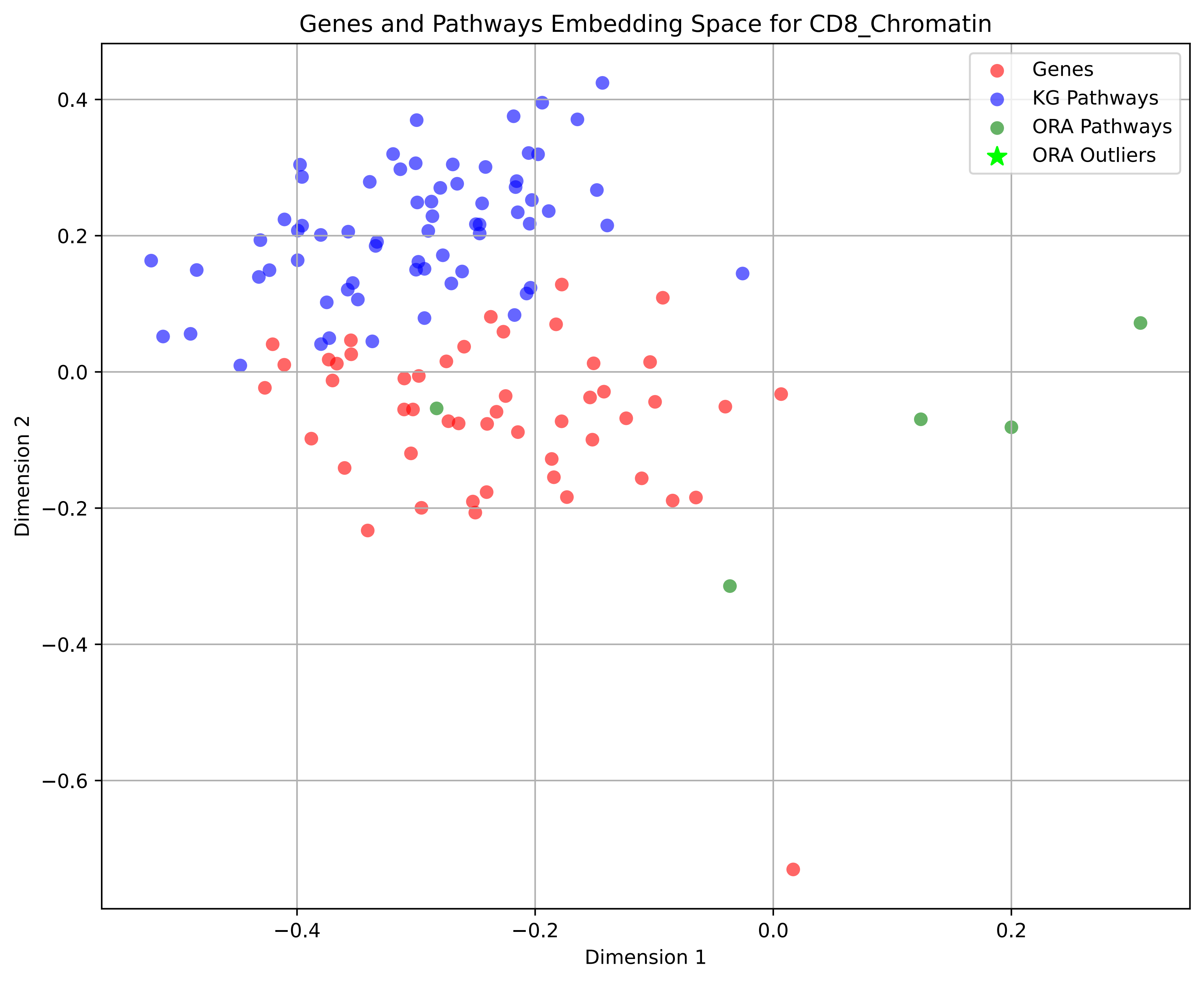
Genes and Pathways Embedding Space for CD8_Chromatin
Genes
KG Pathways
ORA Pathways
ORA Outliers
Dimension 1
Dimension 2
0.4
0.2
0.0
−0.2
−0.4
−0.6
−0.4
−0.2
0.0
0.2

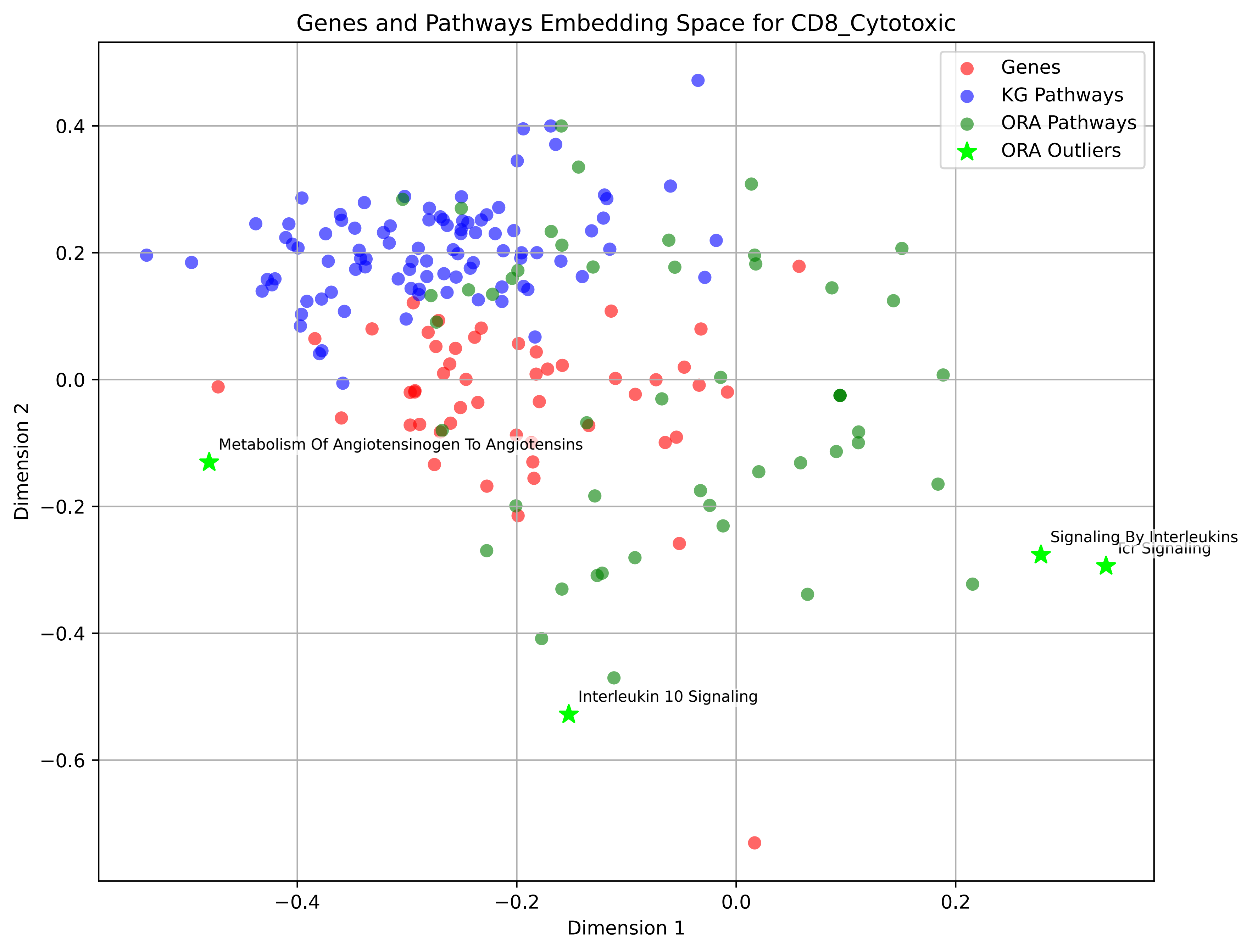

Genes and Pathways Embedding Space for CD8_Cytotoxic
Genes
KG Pathways
ORA Pathways
ORA Outliers
Metabolism Of Angiotensinogen To Angiotensins
Signaling By Interleukins
Tcr Signaling
Interleukin 10 Signaling
0.4
0.2
0.0
−0.2
−0.4
−0.6
−0.4
−0.2
0.0
0.2
Dimension 1
Dimension 2

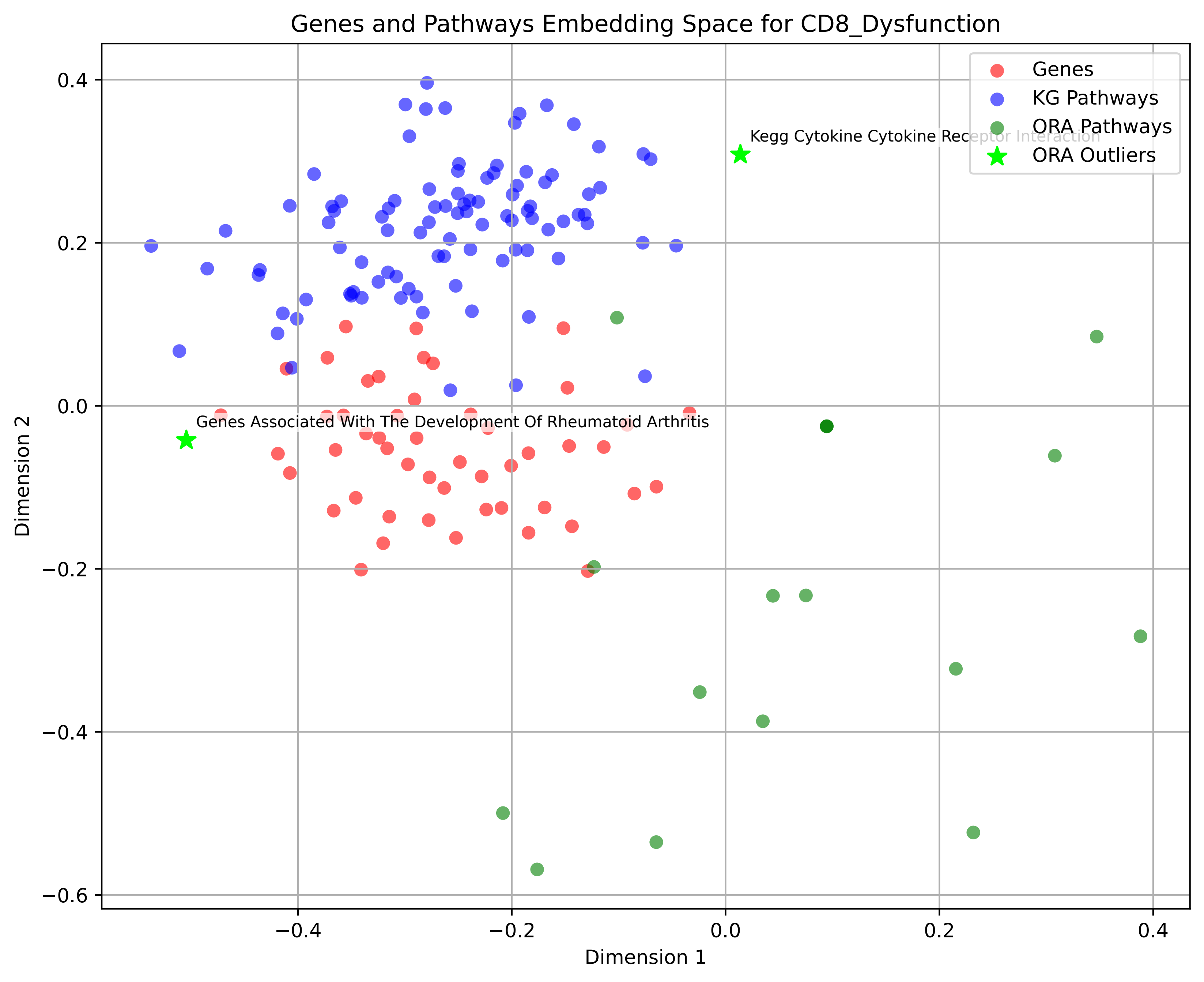

Genes and Pathways Embedding Space for CD8_Dysfunction
Genes
KG Pathways
ORA Pathways
ORA Outliers
Kegg Cytokine Cytokine Receptor Interaction
Genes Associated With The Development Of Rheumatoid Arthritis
Dimension 1
Dimension 2
0.4
0.2
0.0
−0.2
−0.4
−0.6
−0.4
−0.2
0.0
0.2
0.4

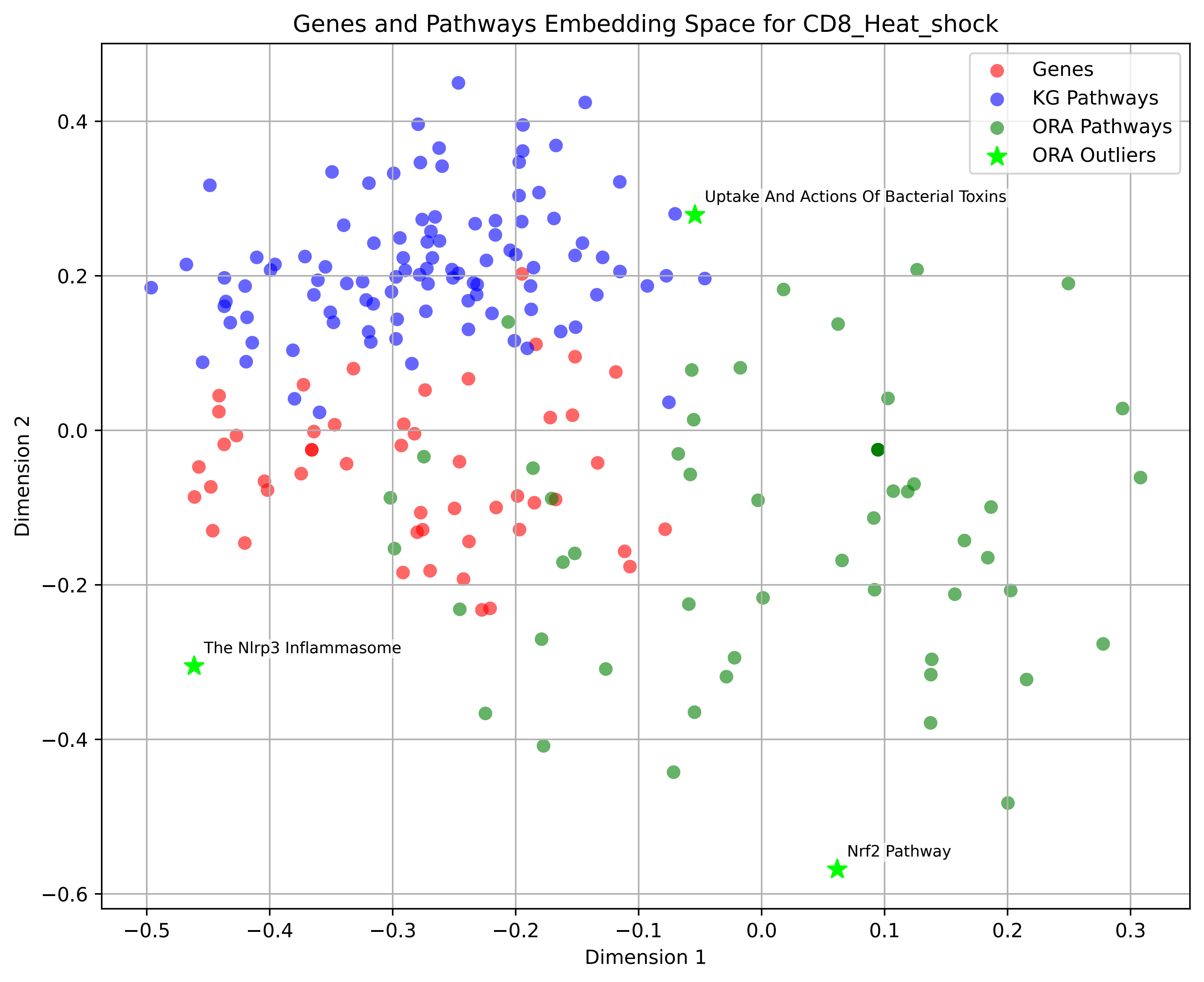

Genes and Pathways Embedding Space for CD8_Heat_shock
Genes
KG Pathways
ORA Pathways
ORA Outliers
Uptake And Actions Of Bacterial Toxins
The Nlrp3 Inflammasome
Nrf2 Pathway
Dimension 1
Dimension 2
0.4
0.2
0.0
−0.2
−0.4
−0.6
−0.5
−0.4
−0.3
−0.2
−0.1
0.0
0.1
0.2
0.3

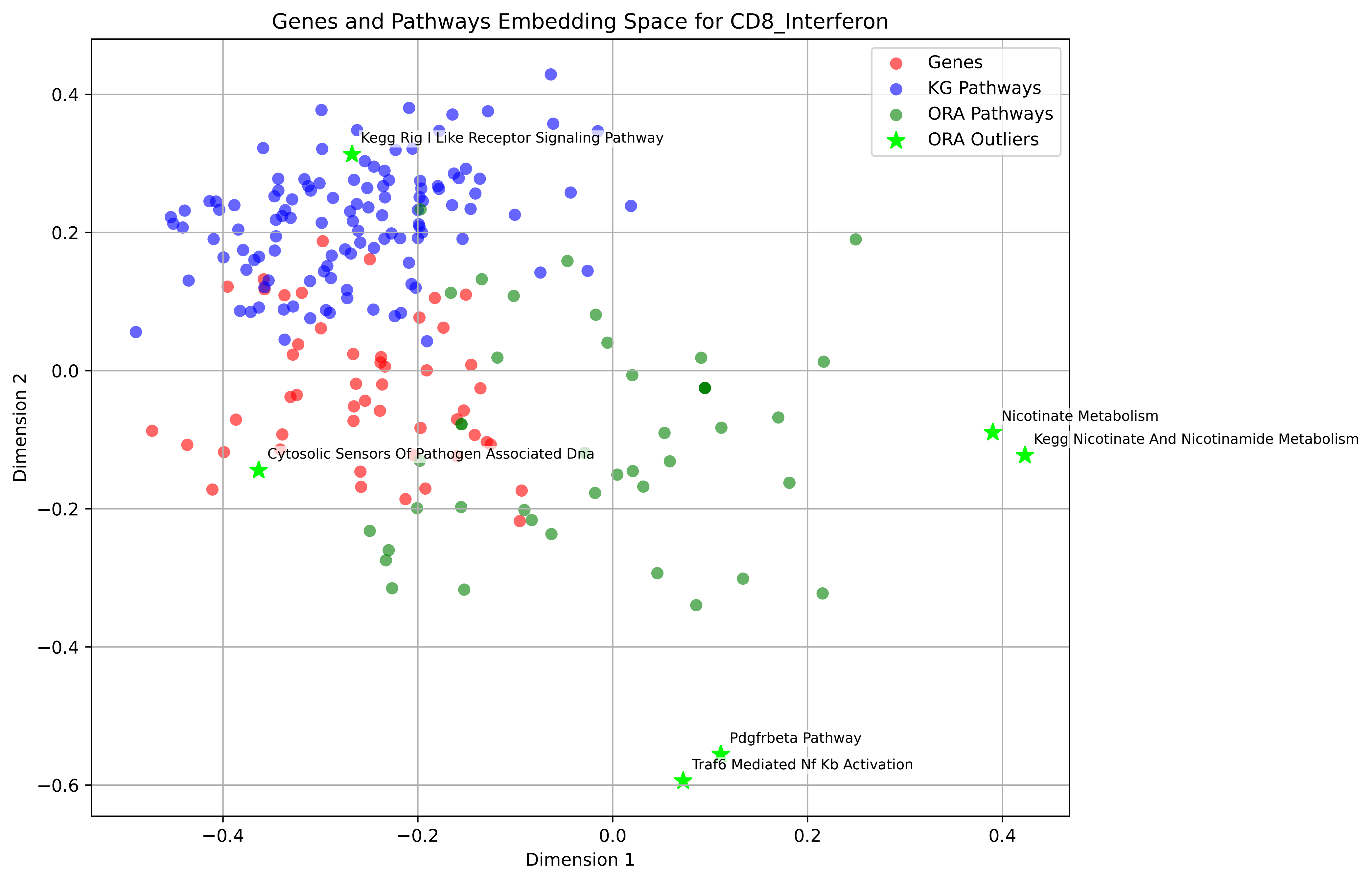

Genes and Pathways Embedding Space for CD8_Interferon
Genes
KG Pathways
ORA Pathways
ORA Outliers
Kegg Rig I Like Receptor Signaling Pathway
Cytosolic Sensors Of Pathogen Associated Dna
Nicotinate Metabolism
Kegg Nicotinate And Nicotinamide Metabolism
Pdgfrbeta Pathway
Traf6 Mediated Nf Kb Activation
Dimension 1
Dimension 2
0.4
0.2
0.0
−0.2
−0.4
−0.6

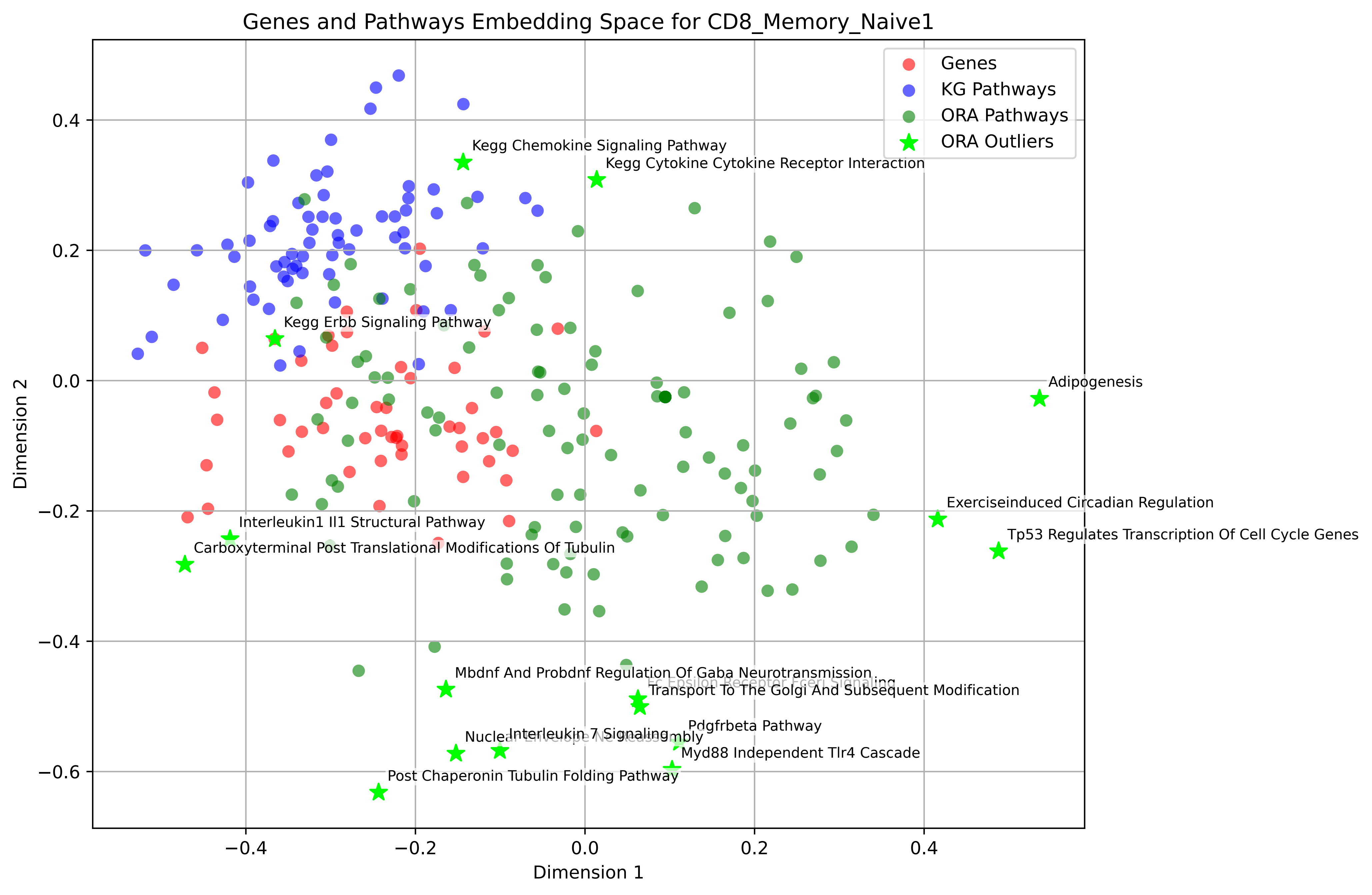

Genes and Pathways Embedding Space for CD8_Memory_Naive1
Genes
KG Pathways
ORA Pathways
ORA Outliers
Kegg Chemokine Signaling Pathway
Kegg Cytokine Cytokine Receptor Interaction
Kegg Erbb Signaling Pathway
Adipogenesis
Exerciseinduced Circadian Regulation
Tp53 Regulates Transcription Of Cell Cycle Genes
Interleukin1 Il1 Structural Pathway
Carboxyterminal Post Translational Modifications Of Tubulin
Mbdnf And Probdnf Regulation Of Gaba Neurotransmission
Fc Epsilon Receptor Fceri Signaling
Transport To The Golgi And Subsequent Modification
Pdgfrbeta Pathway
Interleukin 7 Signaling
Myd88 Independent Tlr4 Cascade
Post Chaperonin Tubulin Folding Pathway
Dimension 1
Dimension 2
0.4
0.2
0.0
−0.2
−0.4
−0.6

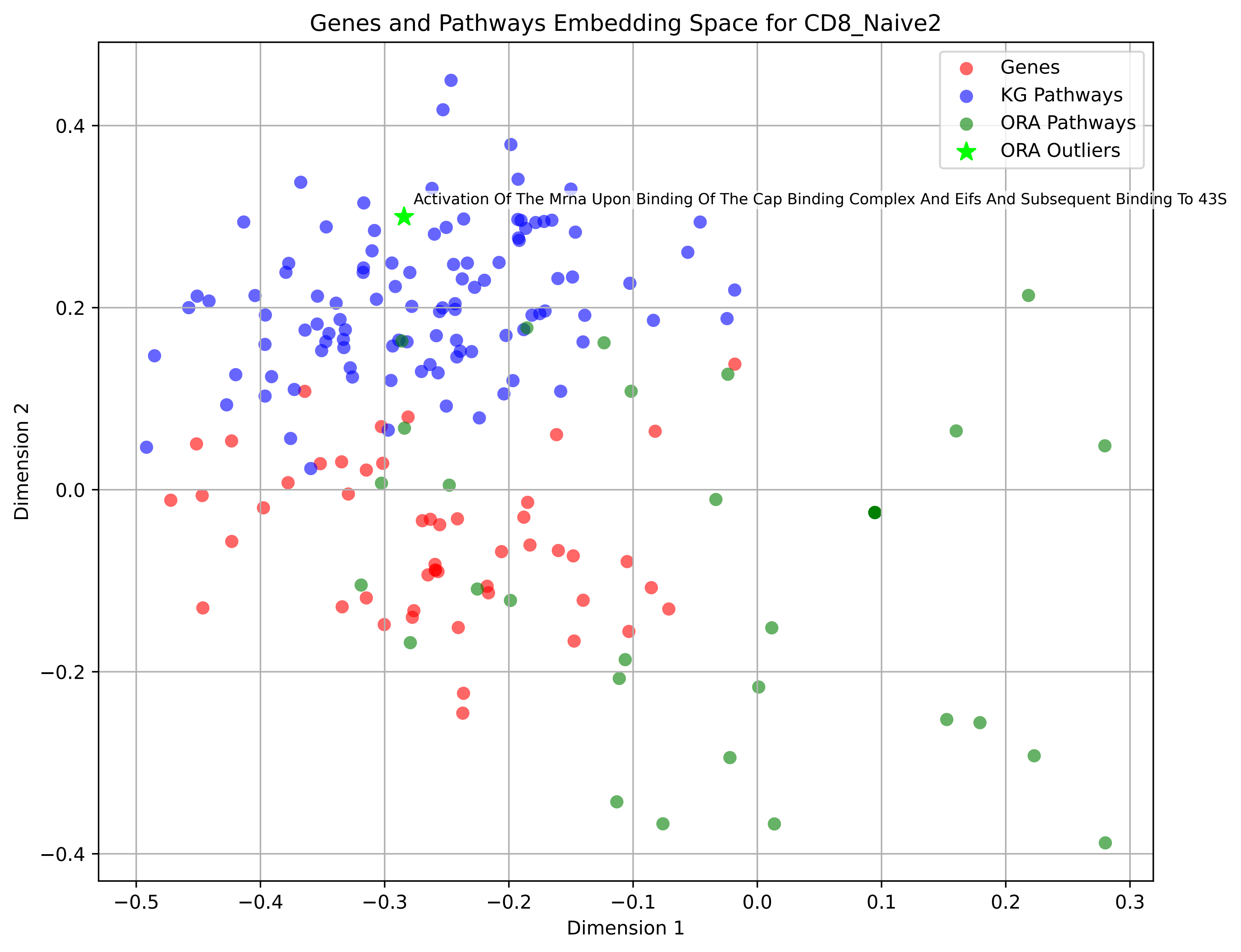
Genes and Pathways Embedding Space for CD8_Naive2
Genes
KG Pathways
ORA Pathways
ORA Outliers
Activation Of The Mrna Upon Binding Of The Cap Binding Complex And Eifs And Subsequent Binding To 43S
Dimension 1
Dimension 2
−0.5
−0.4
−0.3
−0.2
−0.1
0.0
0.1
0.2
0.3
−0.4
−0.2
0.0
0.2
0.4

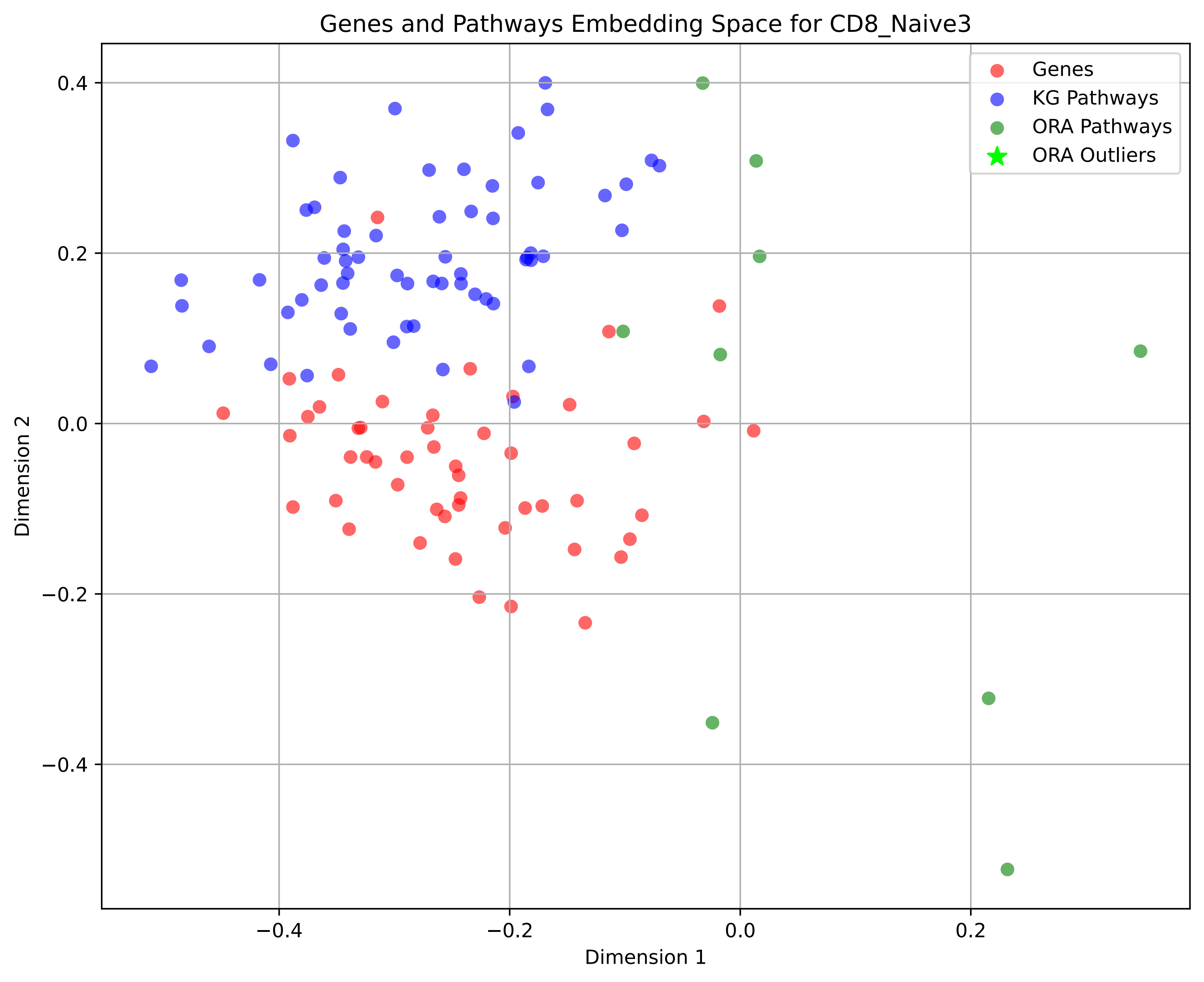

Genes and Pathways Embedding Space for CD8_Naive3
Genes
KG Pathways
ORA Pathways
ORA Outliers
Dimension 1
Dimension 2
0.4
0.2
0.0
−0.2
−0.4
−0.4
−0.2
0.0
0.2

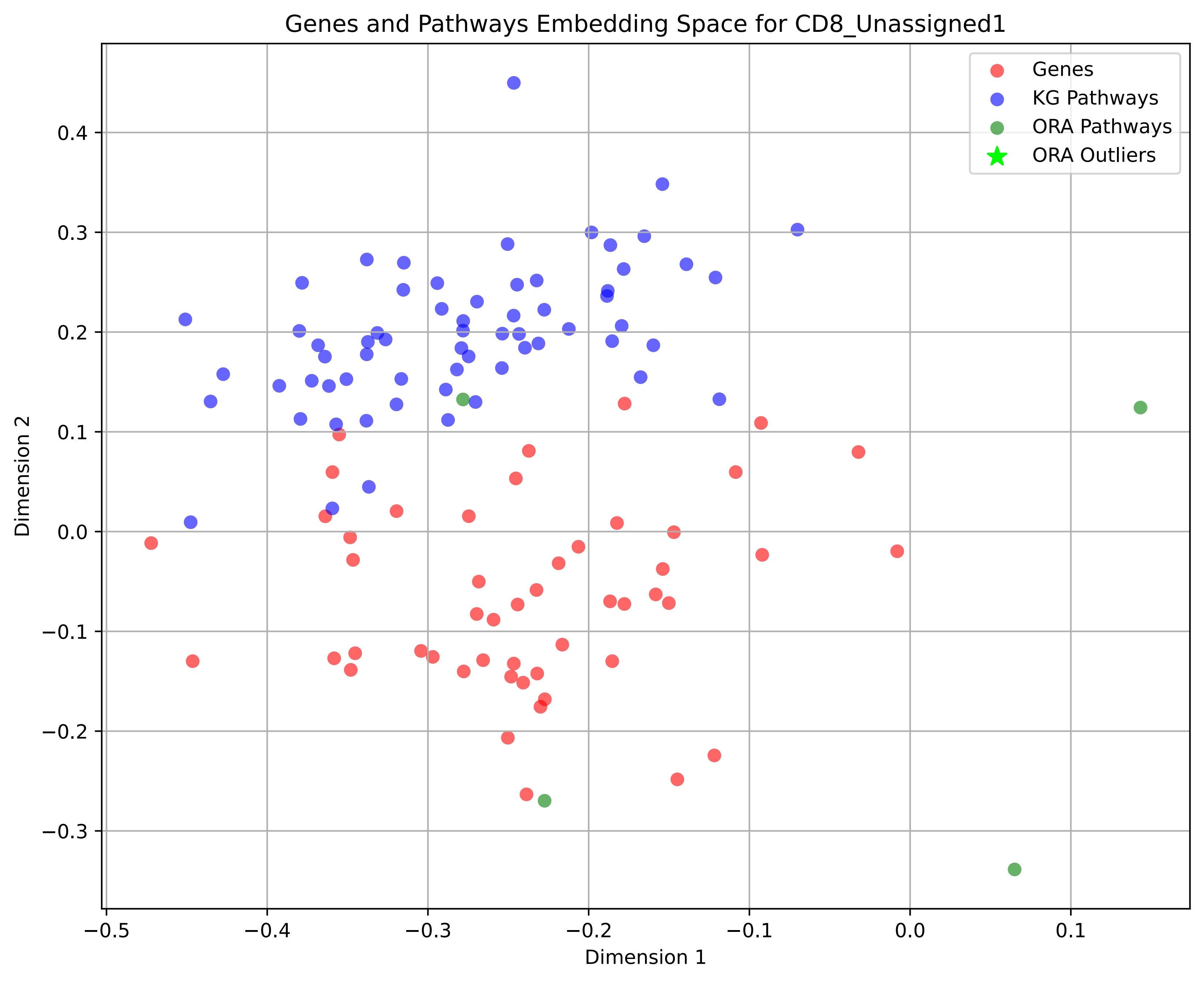
Genes and Pathways Embedding Space for CD8_Unassigned1
Genes
KG Pathways
ORA Pathways
ORA Outliers
Dimension 1
Dimension 2
−0.5
−0.4
−0.3
−0.2
−0.1
0.0
0.1
0.4
0.3
0.2
0.1
0.0
−0.1
−0.2
−0.3

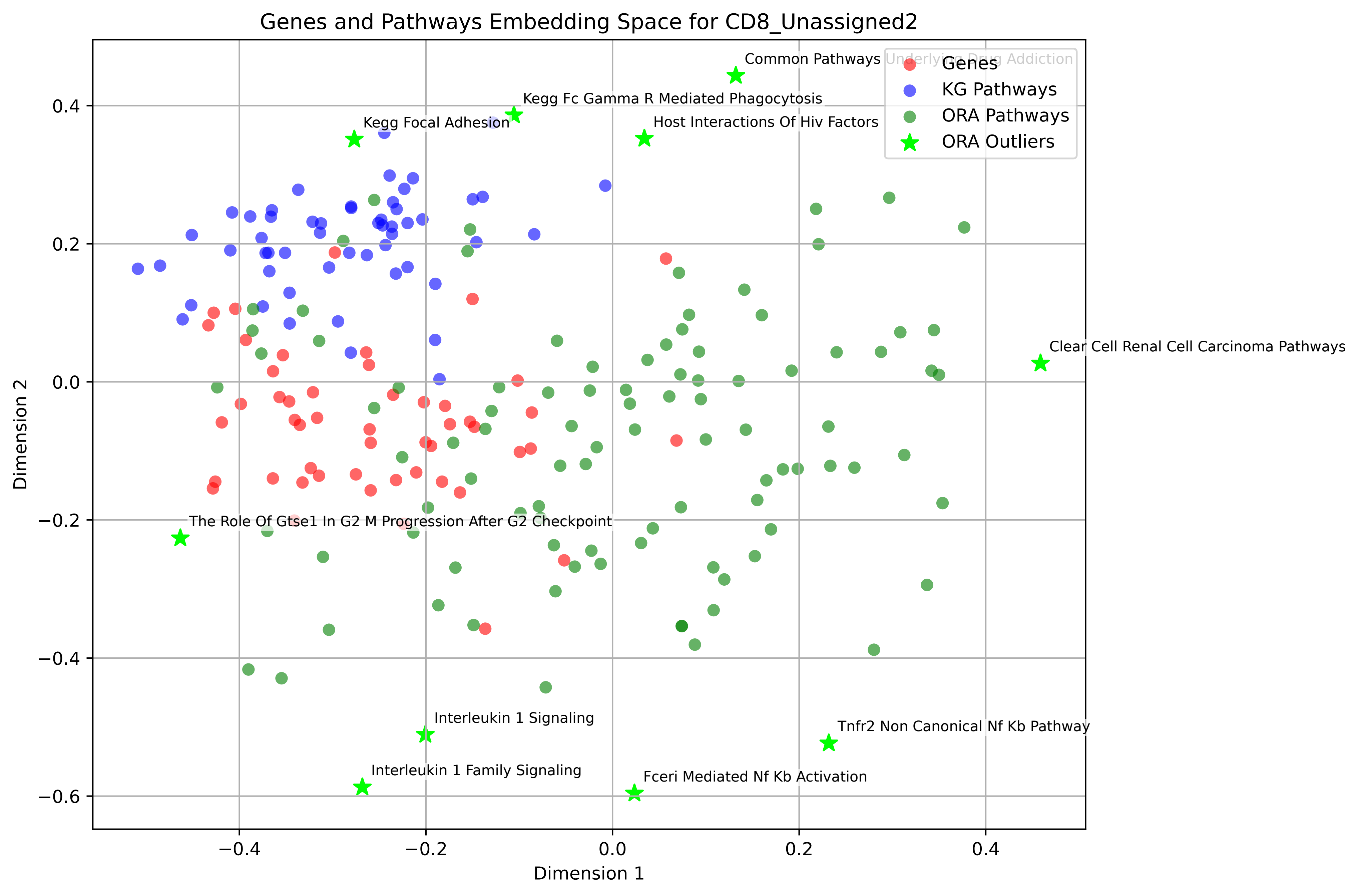

Genes and Pathways Embedding Space for CD8_Unassigned2
Genes
KG Pathways
ORA Pathways
ORA Outliers
Common Pathways Underlying Drug Addiction
Kegg Fc Gamma R Mediated Phagocytosis
Kegg Focal Adhesion
Host Interactions Of Hiv Factors
Clear Cell Renal Cell Carcinoma Pathways
The Role Of Gtse1 In G2 M Progression After G2 Checkpoint
Interleukin 1 Signaling
Tnfr2 Non Canonical Nf Kb Pathway
Interleukin 1 Family Signaling
Fceri Mediated Nf Kb Activation
Dimension 1
Dimension 2
0.4
0.2
0.0
−0.2
−0.4
−0.6
−0.4
−0.2
0.0
0.2
0.4

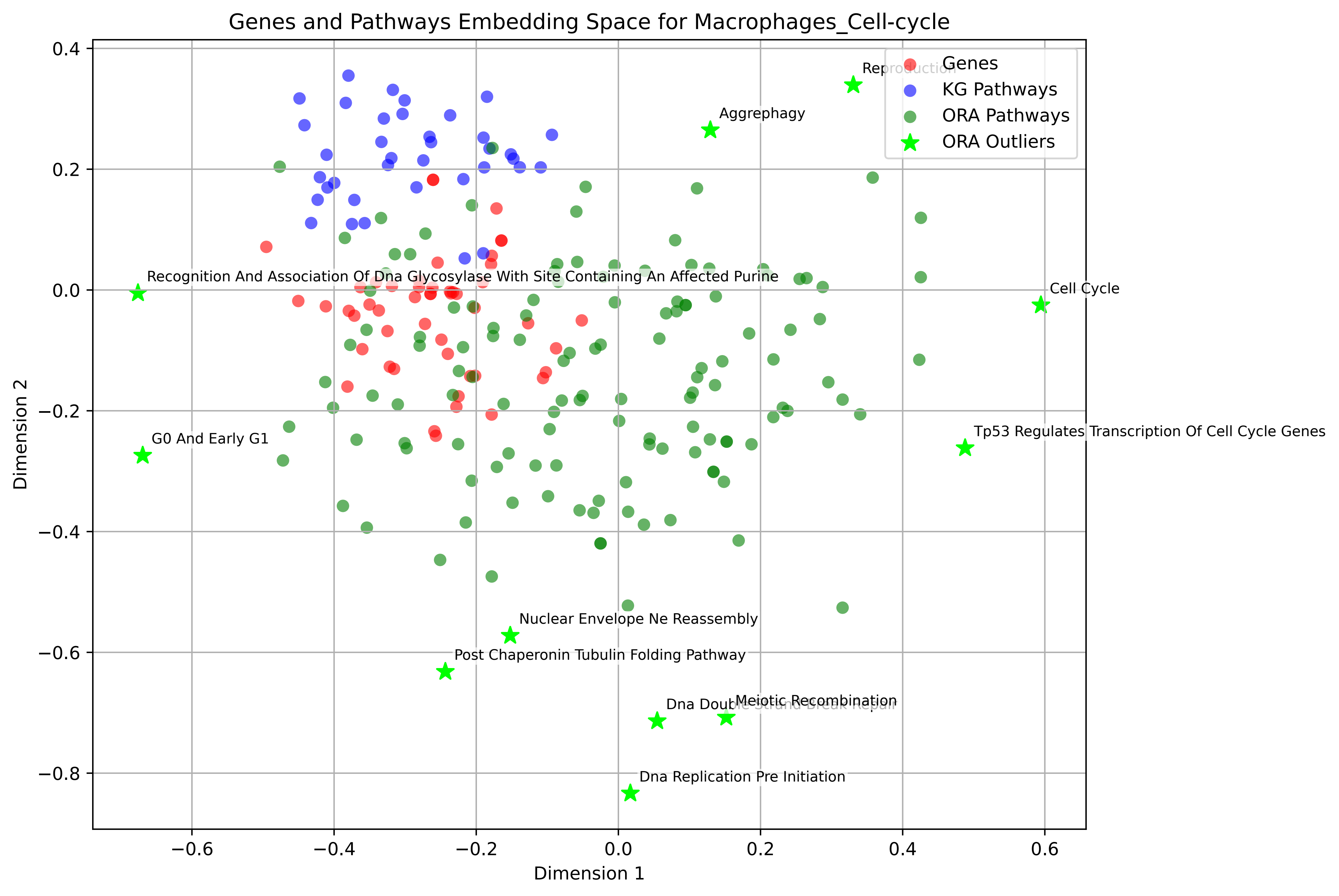

Genes and Pathways Embedding Space for Macrophages_Cell-cycle
Genes
KG Pathways
ORA Pathways
ORA Outliers
Reproduction
Aggrephagy
Recognition And Association Of Dna Glycosylase With Site Containing An Affected Purine
Cell Cycle
G0 And Early G1
Tp53 Regulates Transcription Of Cell Cycle Genes
Nuclear Envelope Ne Reassembly
Post Chaperonin Tubulin Folding Pathway
Meiotic Recombination
Dna Double Strand Break Repair
Dna Replication Pre Initiation
Dimension 1
Dimension 2
0.4
0.2
0.0
−0.2
−0.4
−0.6
−0.8
−0.6
−0.4
−0.2
0.0
0.2
0.4
0.6

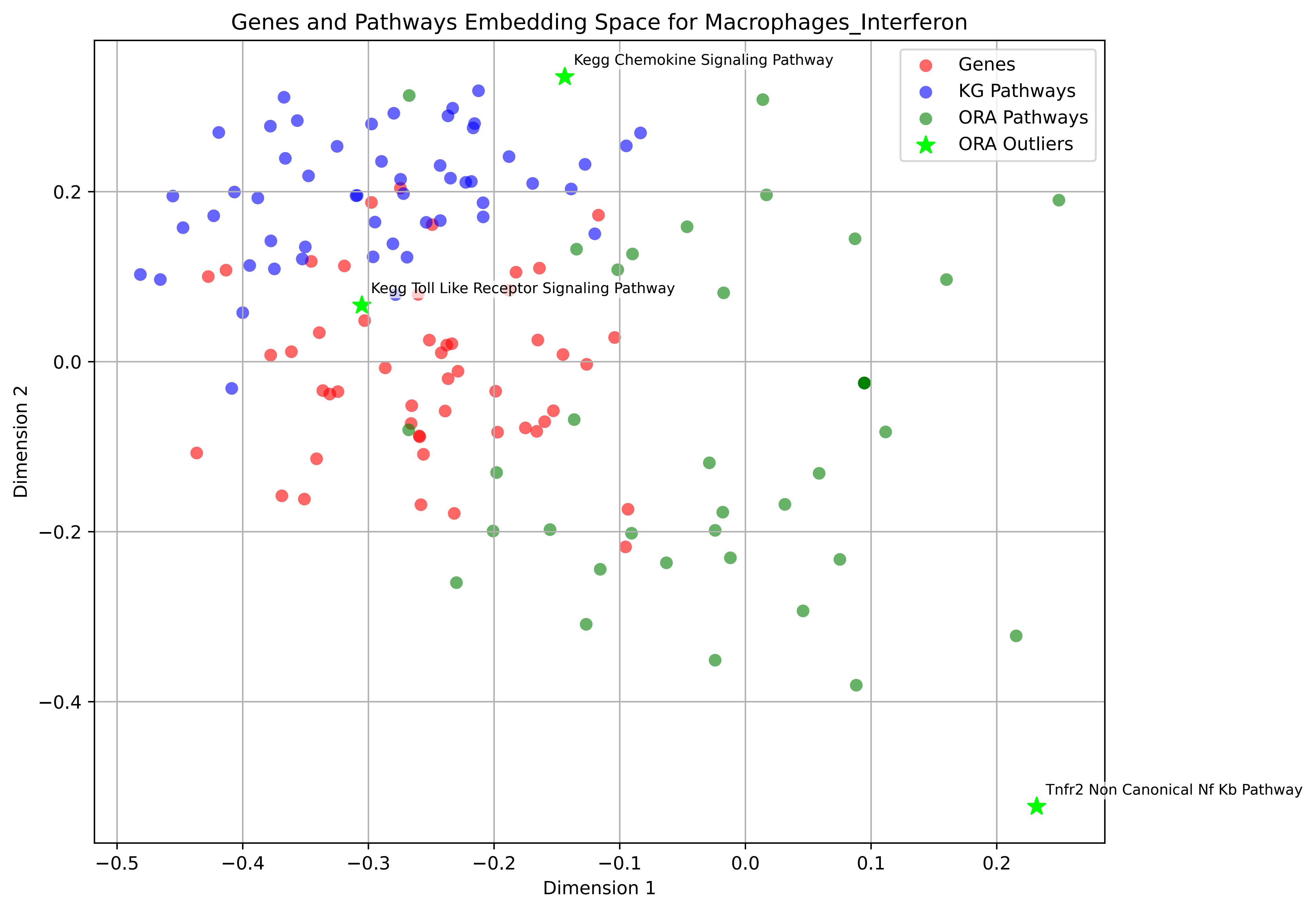

Genes and Pathways Embedding Space for Macrophages_Interferon
Kegg Chemokine Signaling Pathway
Kegg Toll Like Receptor Signaling Pathway
Tnfr2 Non Canonical Nf Kb Pathway
Genes
KG Pathways
ORA Pathways
ORA Outliers
Dimension 1
Dimension 2
−0.5
−0.4
−0.3
−0.2
−0.1
0.0
0.1
0.2
0.2
0.0
−0.2
−0.4

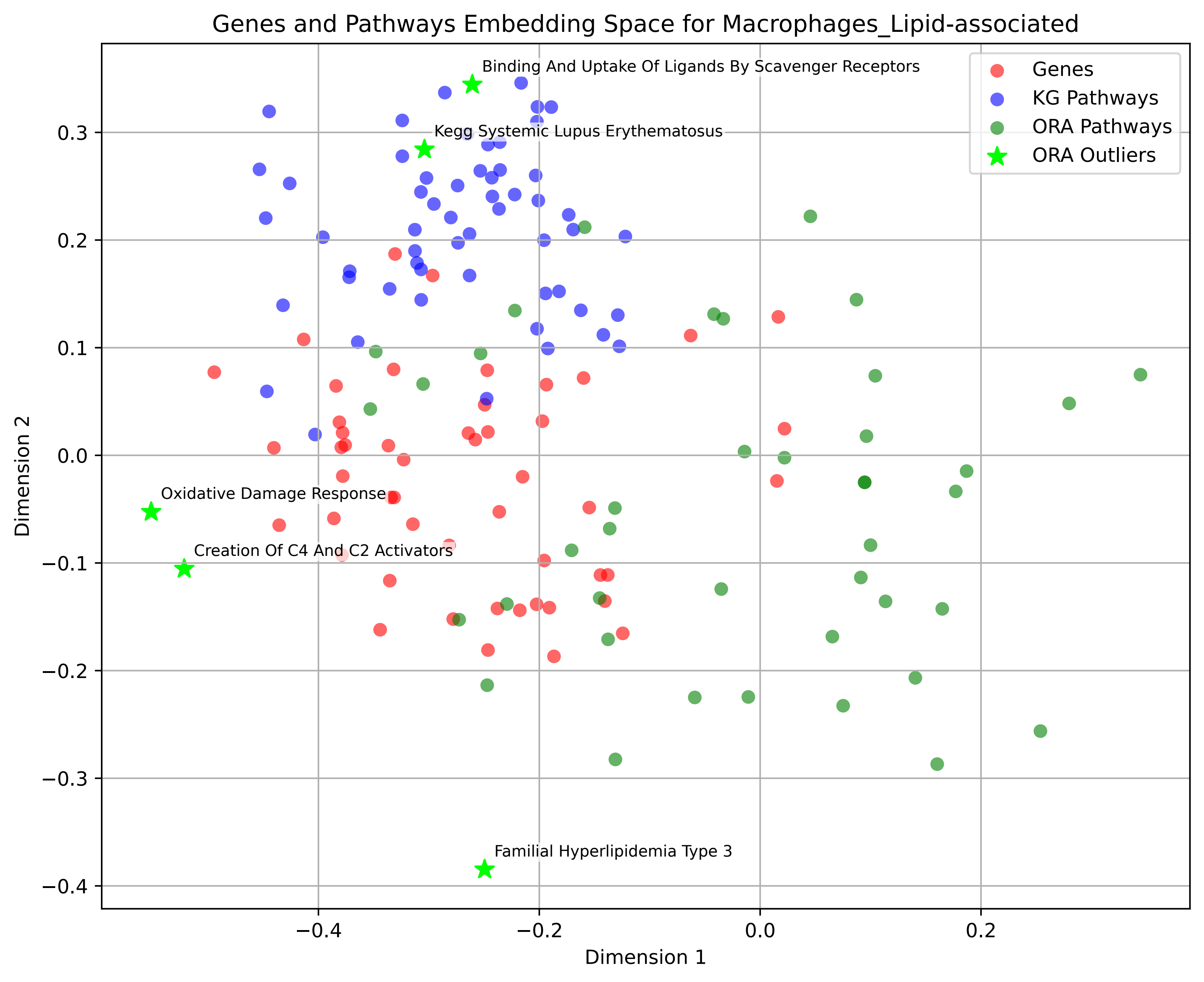
Genes and Pathways Embedding Space for Macrophages_Lipid-associated
Binding And Uptake Of Ligands By Scavenger Receptors
Kegg Systemic Lupus Erythematosus
Oxidative Damage Response
Creation Of C4 And C2 Activators
Familial Hyperlipidemia Type 3
Genes
KG Pathways
ORA Pathways
ORA Outliers
Dimension 1
Dimension 2
0.3
0.2
0.1
0.0
−0.1
−0.2
−0.3
−0.4
−0.4
−0.2
0.0
0.2

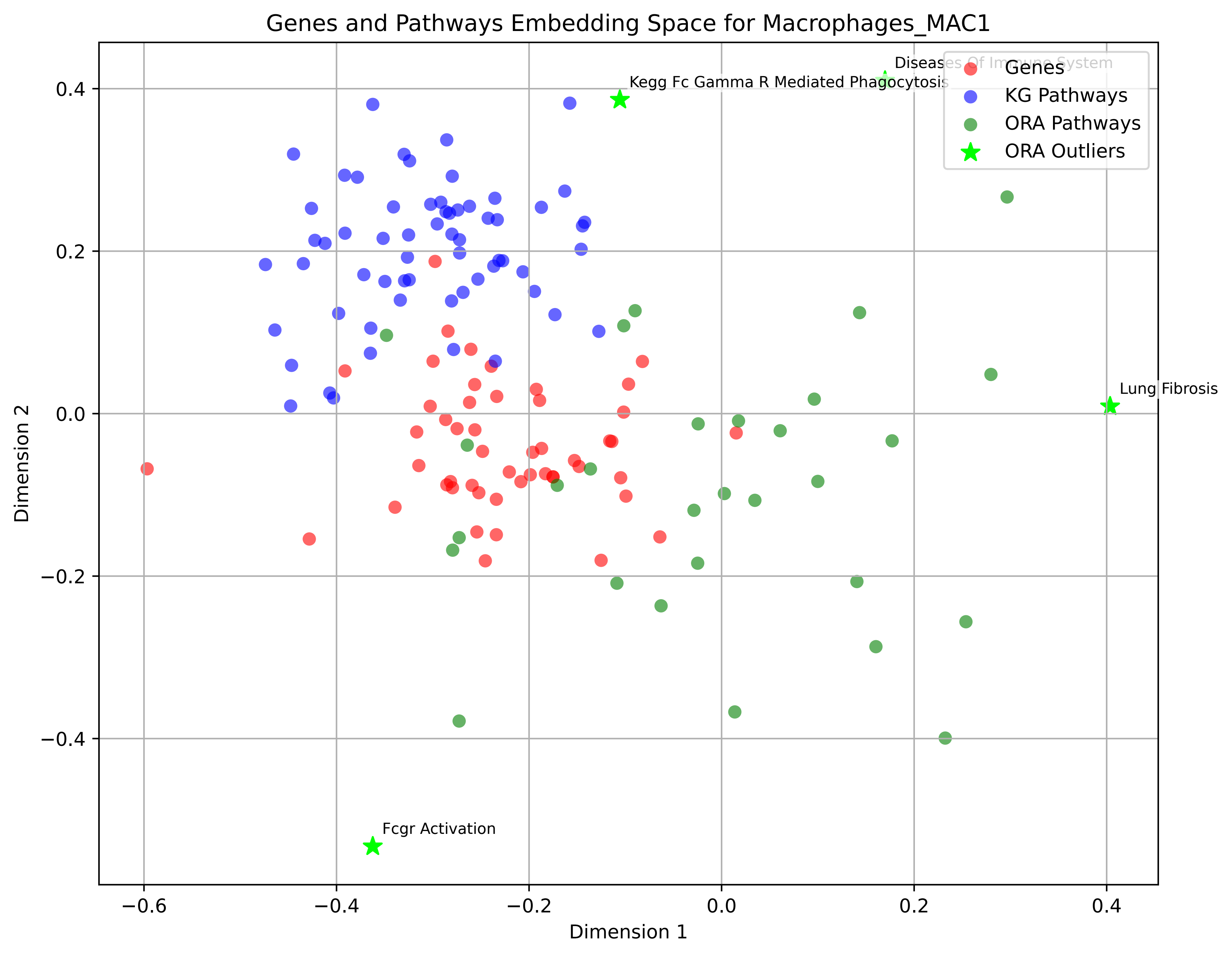

Genes and Pathways Embedding Space for Macrophages_MAC1
Genes
KG Pathways
ORA Pathways
ORA Outliers
Diseases Of Immune System
Kegg Fc Gamma R Mediated Phagocytosis
Lung Fibrosis
Fcgr Activation
Dimension 1
Dimension 2
0.4
0.2
0.0
−0.2
−0.4
−0.6

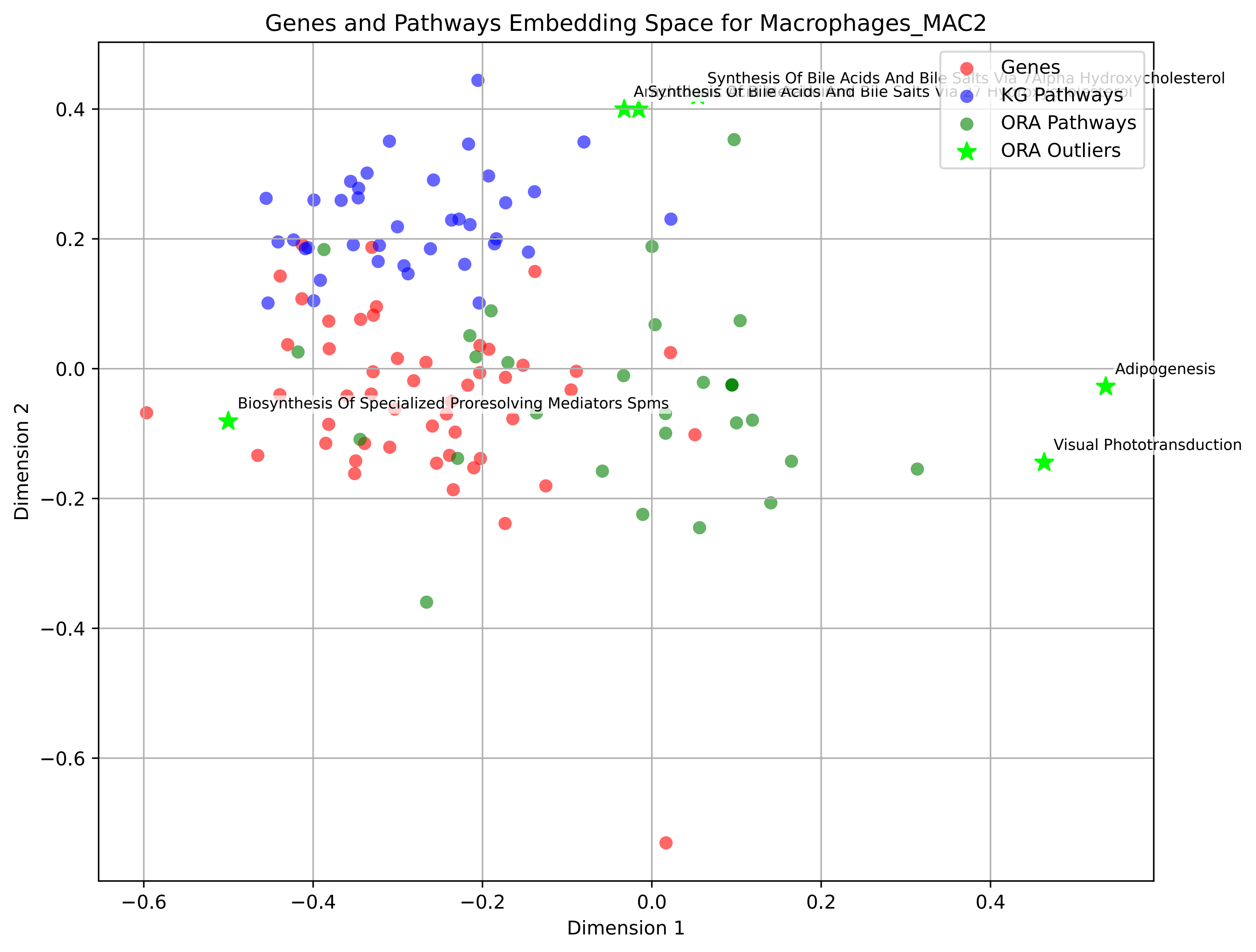

Genes and Pathways Embedding Space for Macrophages_MAC2
Genes
KG Pathways
ORA Pathways
ORA Outliers
Synthesis Of Bile Acids And Bile Salts Via 7Alpha Hydroxycholesterol
Adipogenesis
Visual Phototransduction
Biosynthesis Of Specialized Proresolving Mediators Spms
Dimension 1
Dimension 2
−0.6
−0.4
−0.2
0.0
0.2
0.4

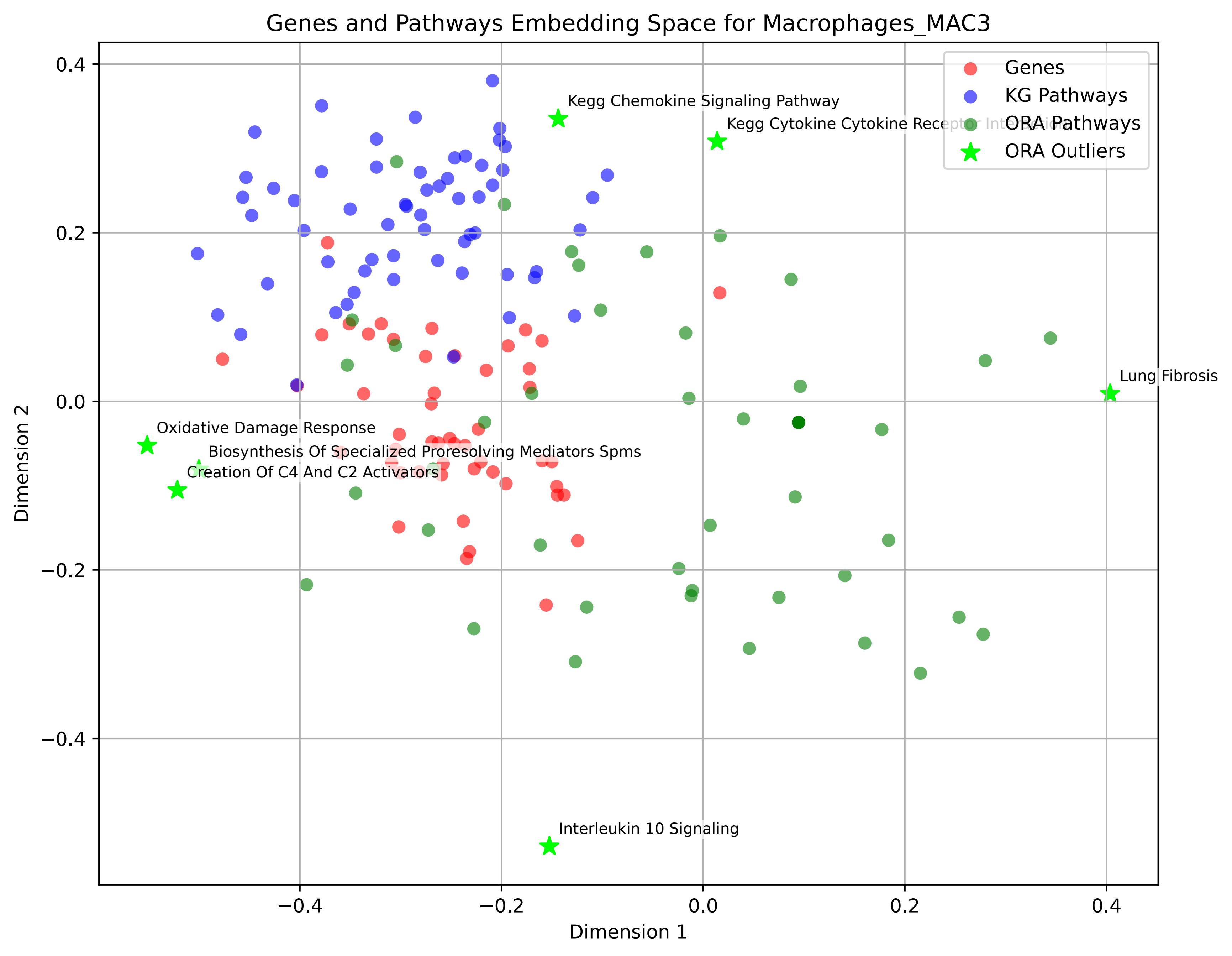

Genes and Pathways Embedding Space for Macrophages_MAC3
Genes
KG Pathways
ORA Pathways
ORA Outliers
Kegg Chemokine Signaling Pathway
Kegg Cytokine Cytokine Receptor Interaction
Lung Fibrosis
Oxidative Damage Response
Biosynthesis Of Specialized Proresolving Mediators Spms
Creation Of C4 And C2 Activators
Interleukin 10 Signaling
Dimension 1
Dimension 2
0.4
0.2
0.0
−0.2
−0.4
−0.4
−0.2
0.0
0.2
0.4

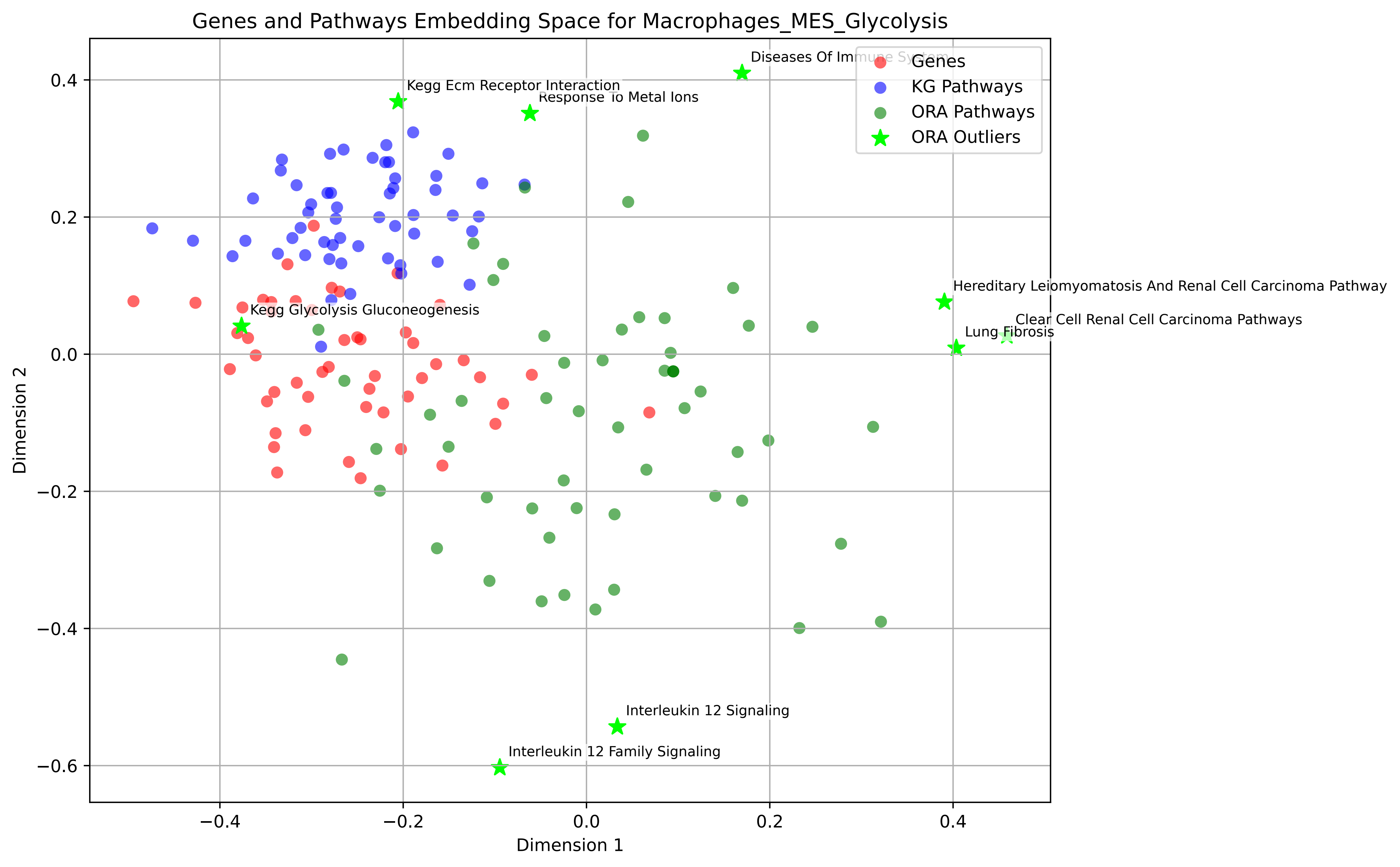

Genes and Pathways Embedding Space for Macrophages_MES_Glycolysis
Genes
KG Pathways
ORA Pathways
ORA Outliers
Diseases Of Immune System
Kegg Ecm Receptor Interaction
Response To Metal Ions
Hereditary Leiomyomatosis And Renal Cell Carcinoma Pathway
Clear Cell Renal Cell Carcinoma Pathways
Lung Fibrosis
Kegg Glycolysis Gluconeogenesis
Interleukin 12 Signaling
Interleukin 12 Family Signaling
Dimension 1
Dimension 2
0.4
0.2
0.0
−0.2
−0.4
−0.6
−0.4
−0.2
0.0
0.2
0.4

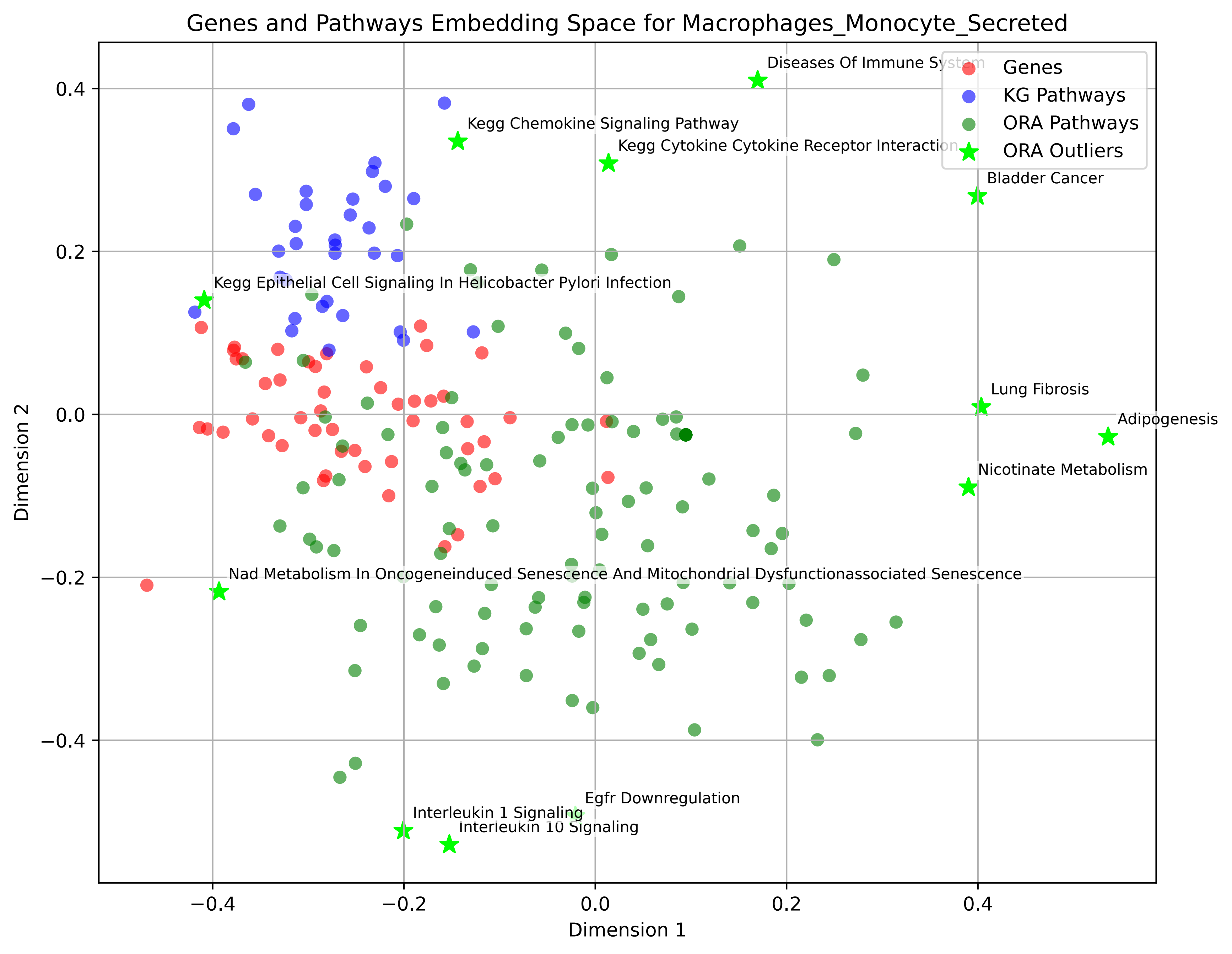

Genes and Pathways Embedding Space for Macrophages_Monocyte_Secreted
Genes
KG Pathways
ORA Pathways
ORA Outliers
Diseases Of Immune System
Kegg Chemokine Signaling Pathway
Kegg Cytokine Cytokine Receptor Interaction
Bladder Cancer
Kegg Epithelial Cell Signaling In Helicobacter Pylori Infection
Lung Fibrosis
Adipogenesis
Nicotinate Metabolism
Nad Metabolism In Oncogeneinduced Senescence And Mitochondrial Dysfunctionassociated Senescence
Egfr Downregulation
Interleukin 1 Signaling
Interleukin 10 Signaling
Dimension 1
Dimension 2
0.4
0.2
0.0
−0.2
−0.4

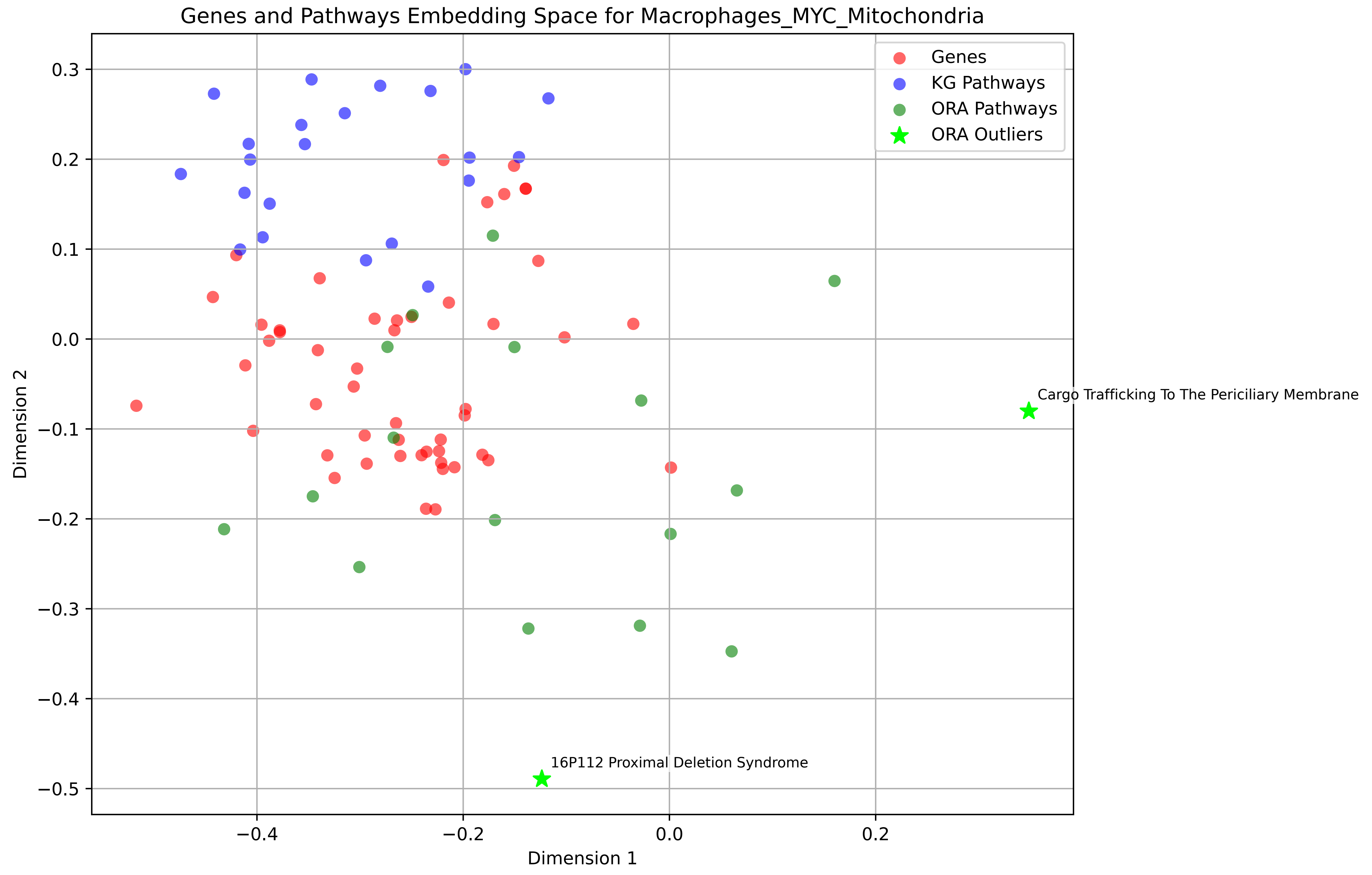
Genes and Pathways Embedding Space for Macrophages_MYC_Mitochondria
Genes
KG Pathways
ORA Pathways
ORA Outliers
Cargo Trafficking To The Periciliary Membrane
16P112 Proximal Deletion Syndrome
Dimension 1
Dimension 2
0.3
0.2
0.1
0.0
−0.1
−0.2
−0.3
−0.4
−0.5
−0.4
−0.2
0.0
0.2

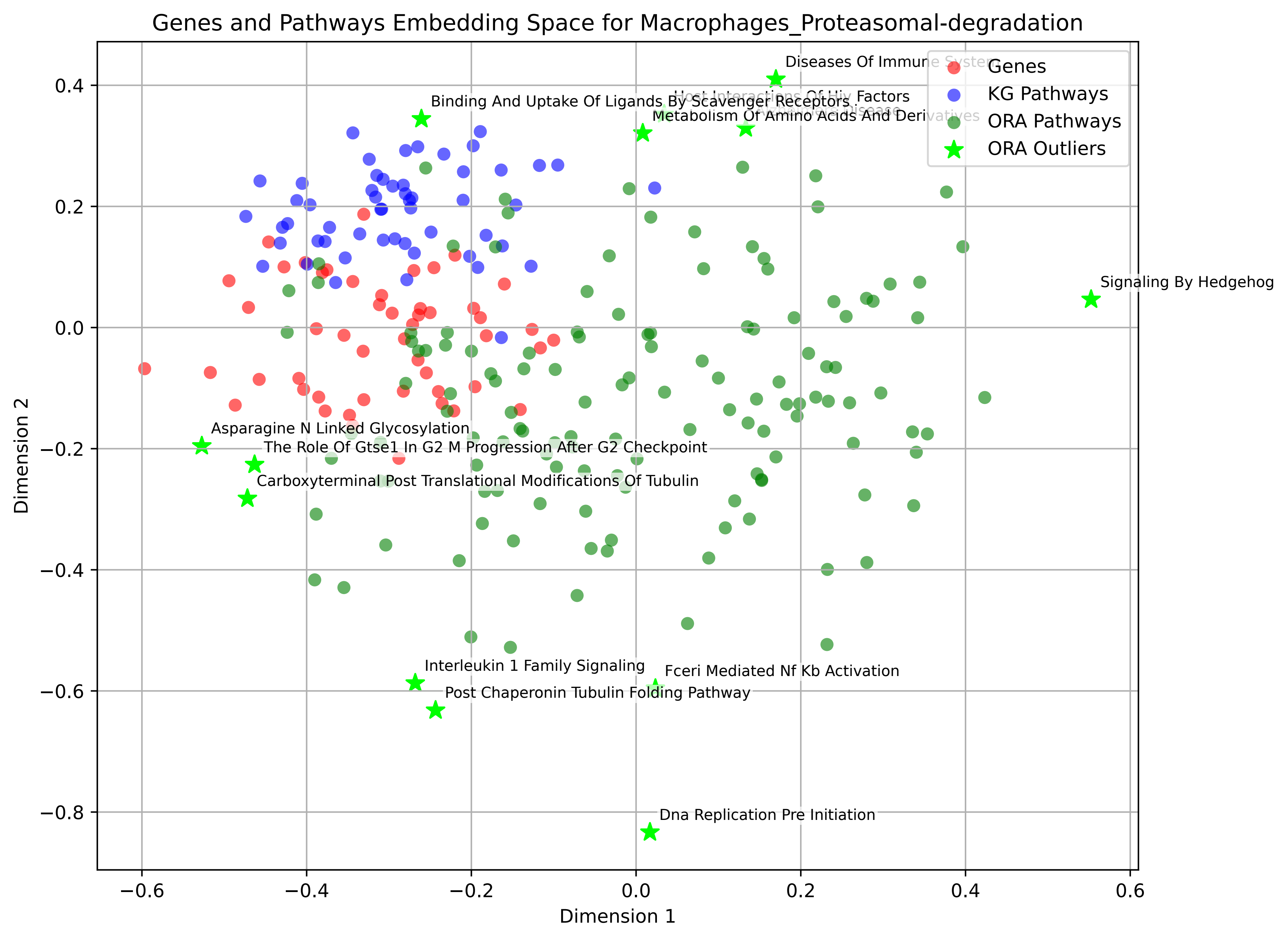

Genes and Pathways Embedding Space for Macrophages_Proteasomal-degradation
Diseases Of Immune System
Host Interactions Of Hiv Factors
Binding And Uptake Of Ligands By Scavenger Receptors
Alzheimers Disease
Metabolism Of Amino Acids And Derivatives
Signaling By Hedgehog
Asparagine N Linked Glycosylation
The Role Of Gtse1 In G2 M Progression After G2 Checkpoint
Carboxyterminal Post Translational Modifications Of Tubulin
Interleukin 1 Family Signaling
Fceri Mediated Nf Kb Activation
Post Chaperonin Tubulin Folding Pathway
Dna Replication Pre Initiation
Genes
KG Pathways
ORA Pathways
ORA Outliers
Dimension 1
Dimension 2

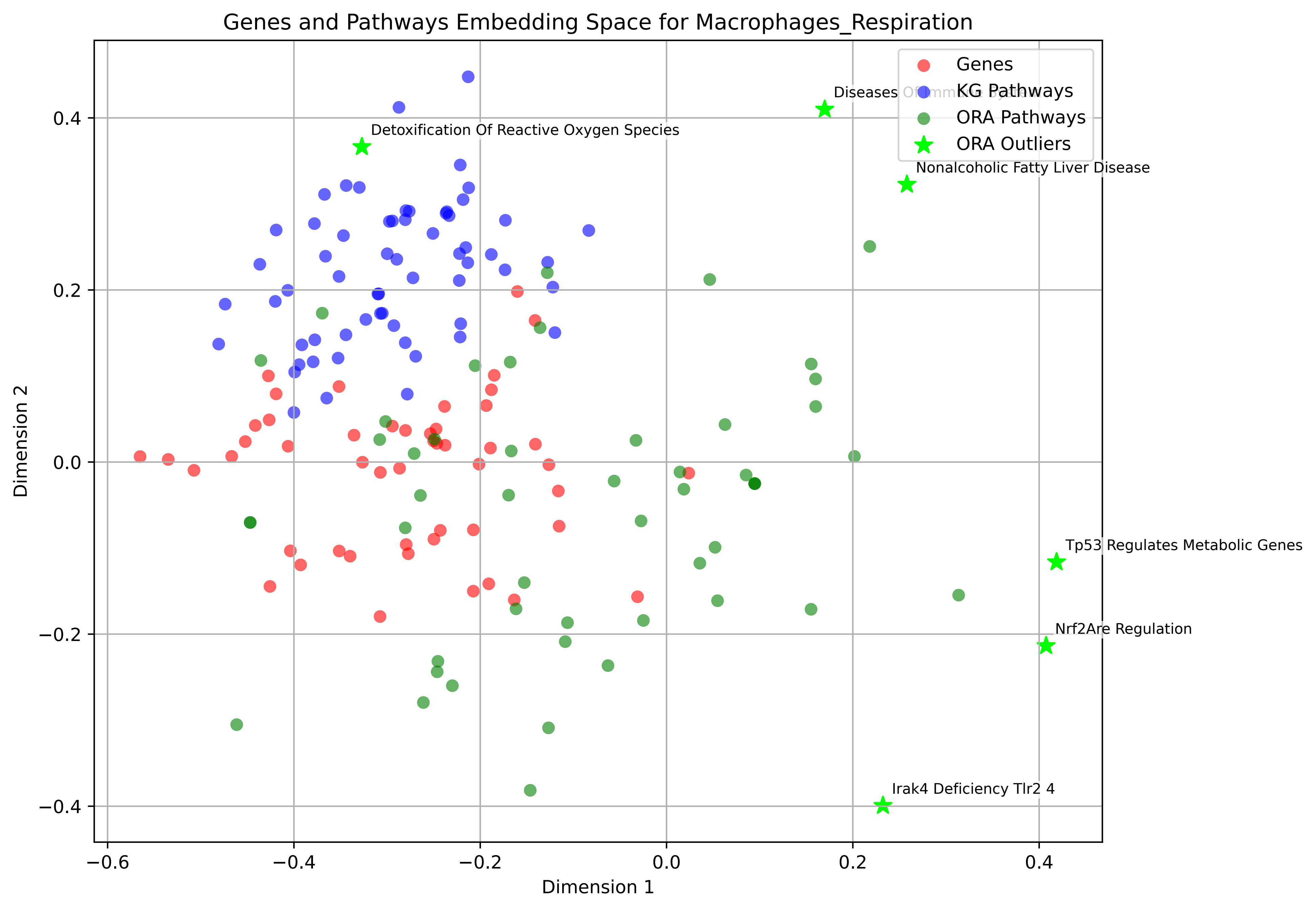
Genes and Pathways Embedding Space for Macrophages_Respiration
Genes
KG Pathways
ORA Pathways
ORA Outliers
Diseases
Detoxification Of Reactive Oxygen Species
Nonalcoholic Fatty Liver Disease
Tp53 Regulates Metabolic Genes
Nrf2Are Regulation
Irak4 Deficiency Tlr2 4
Dimension 1
Dimension 2
−0.6
−0.4
−0.2
0.0
0.2
0.4
−0.4
−0.2
0.0
0.2
0.4

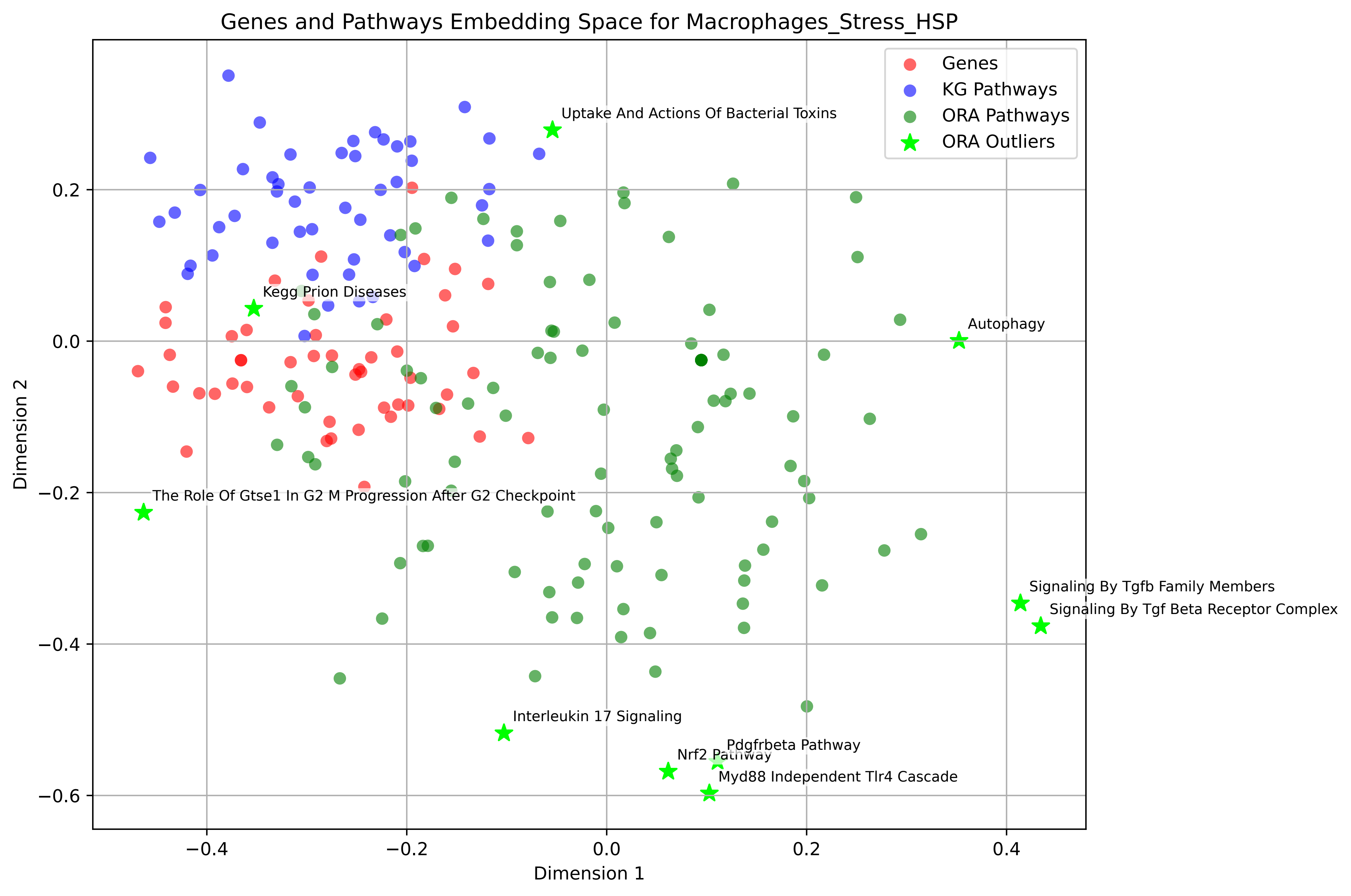

Genes and Pathways Embedding Space for Macrophages_Stress_HSP
Genes
KG Pathways
ORA Pathways
ORA Outliers
Uptake And Actions Of Bacterial Toxins
Kegg Prion Diseases
Autophagy
The Role Of Gtse1 In G2 M Progression After G2 Checkpoint
Signaling By Tgfb Family Members
Signaling By Tgf Beta Receptor Complex
Interleukin 17 Signaling
Pdgfrbeta Pathway
Nrf2 Pathway
Myd88 Independent Tlr4 Cascade
Dimension 1
Dimension 2
−0.4
−0.2
0.0
0.2
0.4
−0.6

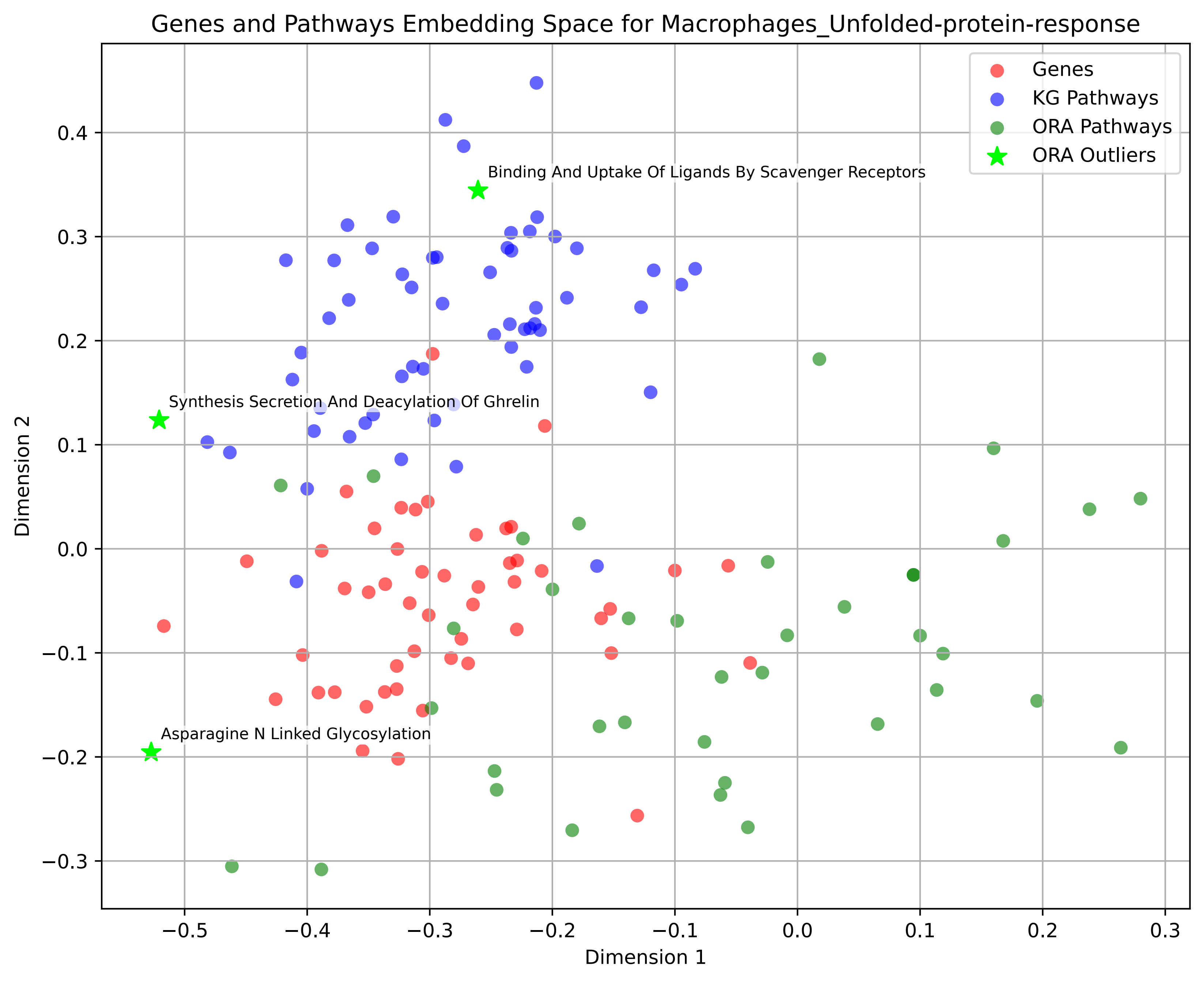

Genes and Pathways Embedding Space for Macrophages_Unfolded-protein-response
Genes
KG Pathways
ORA Pathways
ORA Outliers
Binding And Uptake Of Ligands By Scavenger Receptors
Synthesis Secretion And Deacylation Of Ghrelin
Asparagine N Linked Glycosylation
Dimension 1
Dimension 2
0.4
0.3
0.2
0.1
0.0
−0.1
−0.2
−0.3
−0.5
−0.4
−0.3
−0.2
−0.1
0.0
0.1
0.2
0.3

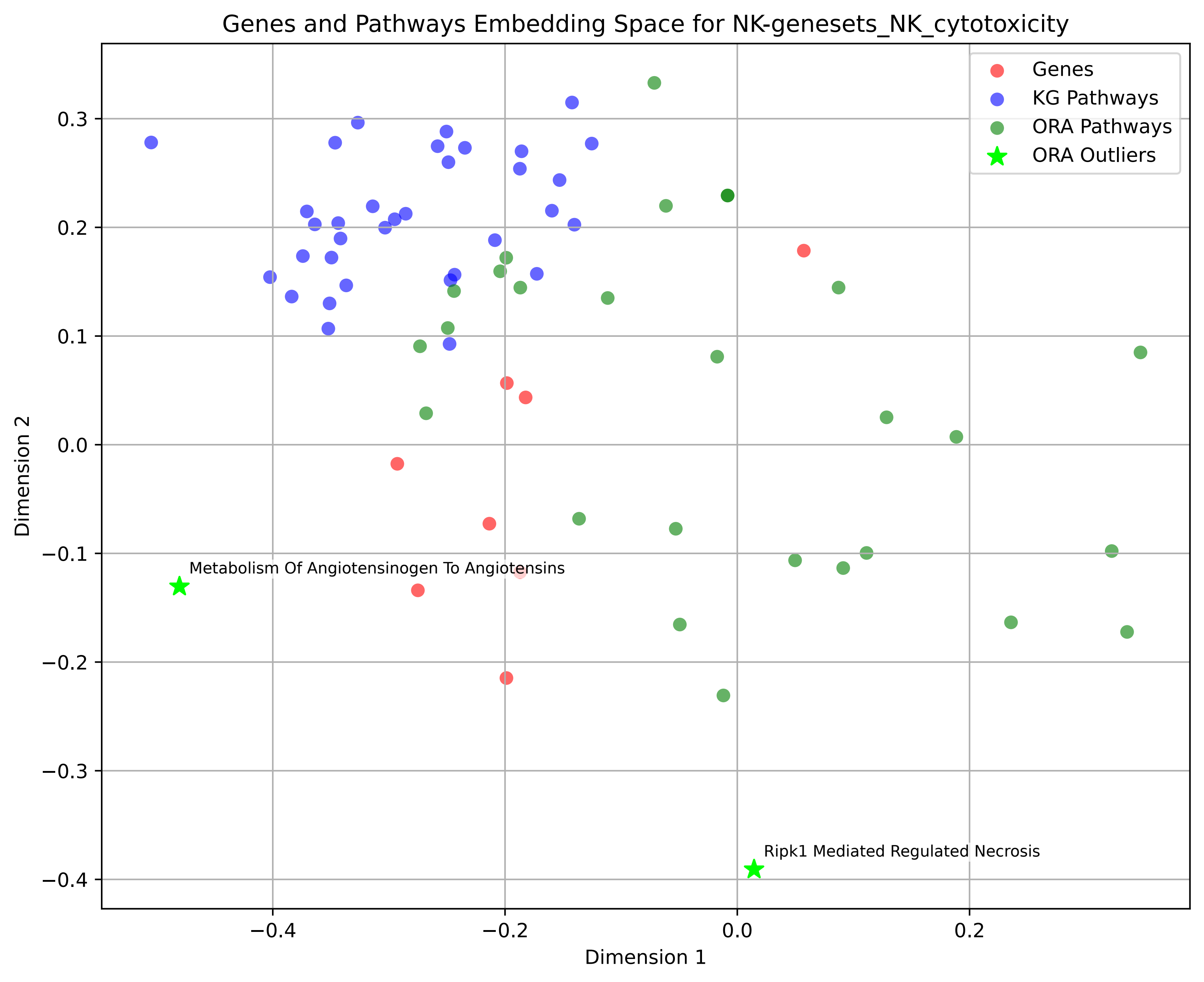
Genes and Pathways Embedding Space for NK-genesets_NK_cytotoxicity
Genes
KG Pathways
ORA Pathways
ORA Outliers
Metabolism Of Angiotensinogen To Angiotensins
Ripk1 Mediated Regulated Necrosis
Dimension 1
Dimension 2
0.3
0.2
0.1
0.0
−0.1
−0.2
−0.3
−0.4
−0.4
−0.2
0.0
0.2

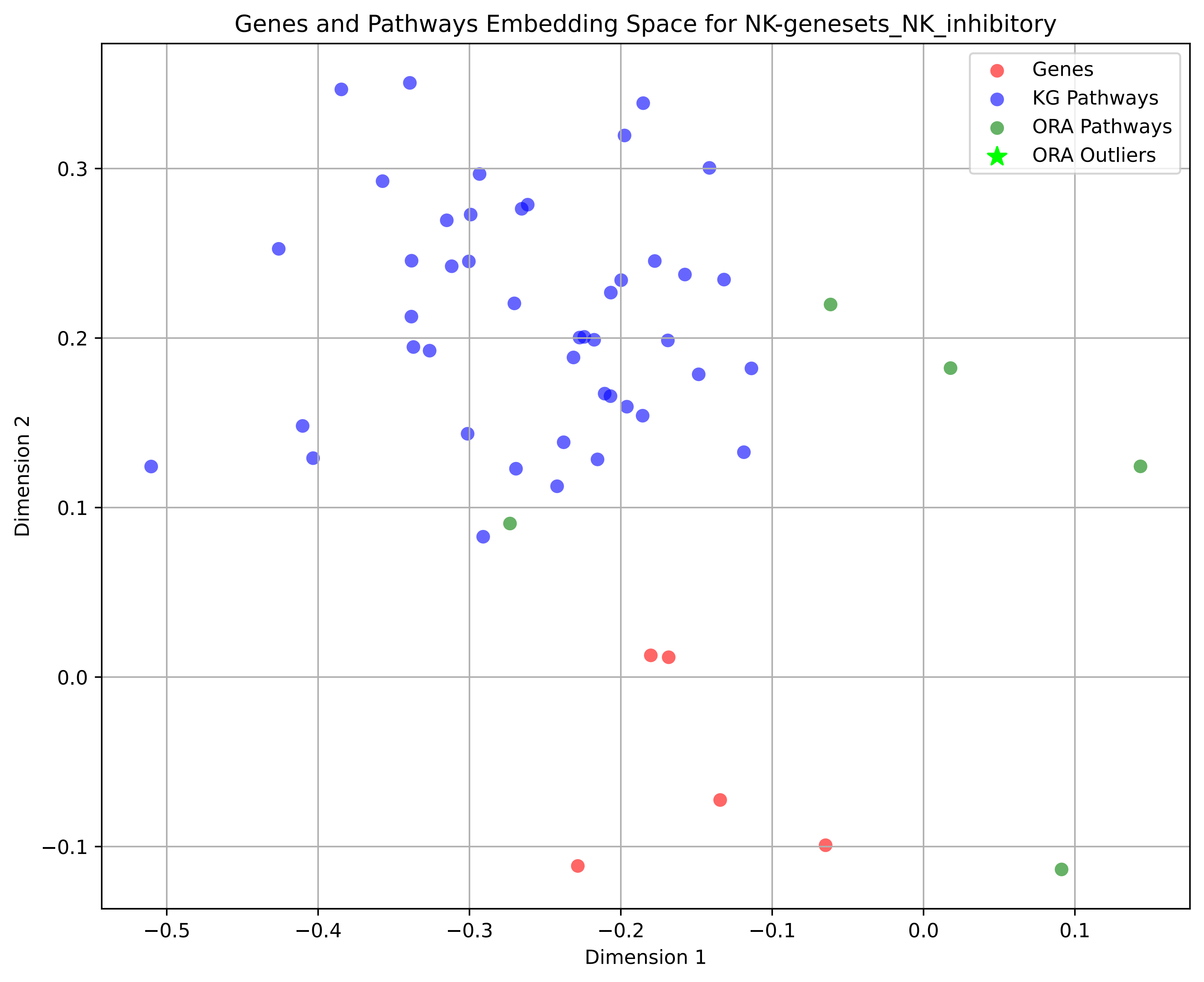
Genes and Pathways Embedding Space for NK-genesets_NK_inhibitory
Genes
KG Pathways
ORA Pathways
ORA Outliers
Dimension 1
Dimension 2
−0.5
−0.4
−0.3
−0.2
−0.1
0.0
0.1
0.3
0.2
0.1
0.0
−0.1

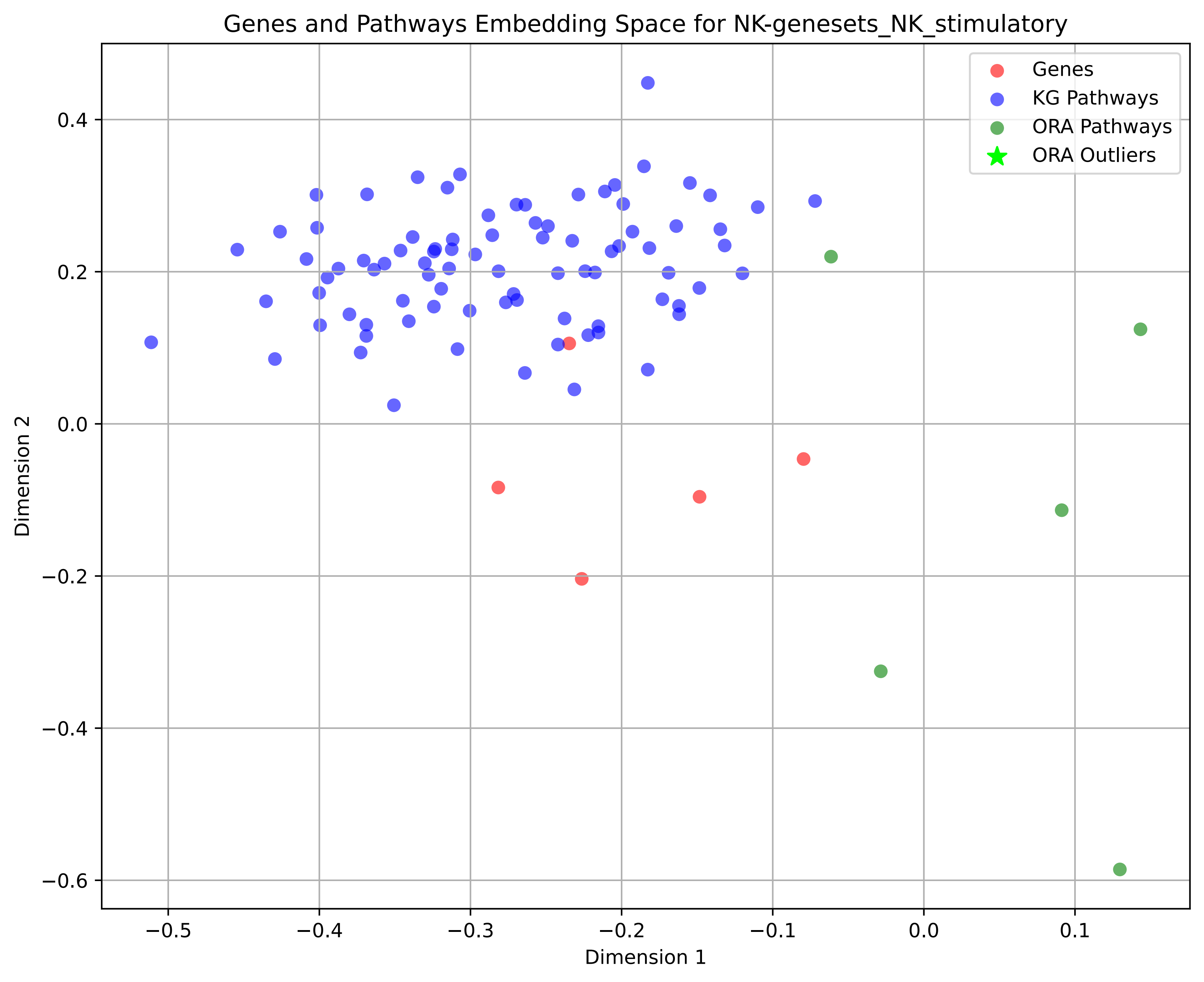

Genes and Pathways Embedding Space for NK-genesets_NK_stimulatory
Genes
KG Pathways
ORA Pathways
ORA Outliers
Dimension 1
Dimension 2
0.4
0.2
0.0
−0.2
−0.4
−0.6
−0.5
−0.4
−0.3
−0.2
−0.1
0.0
0.1

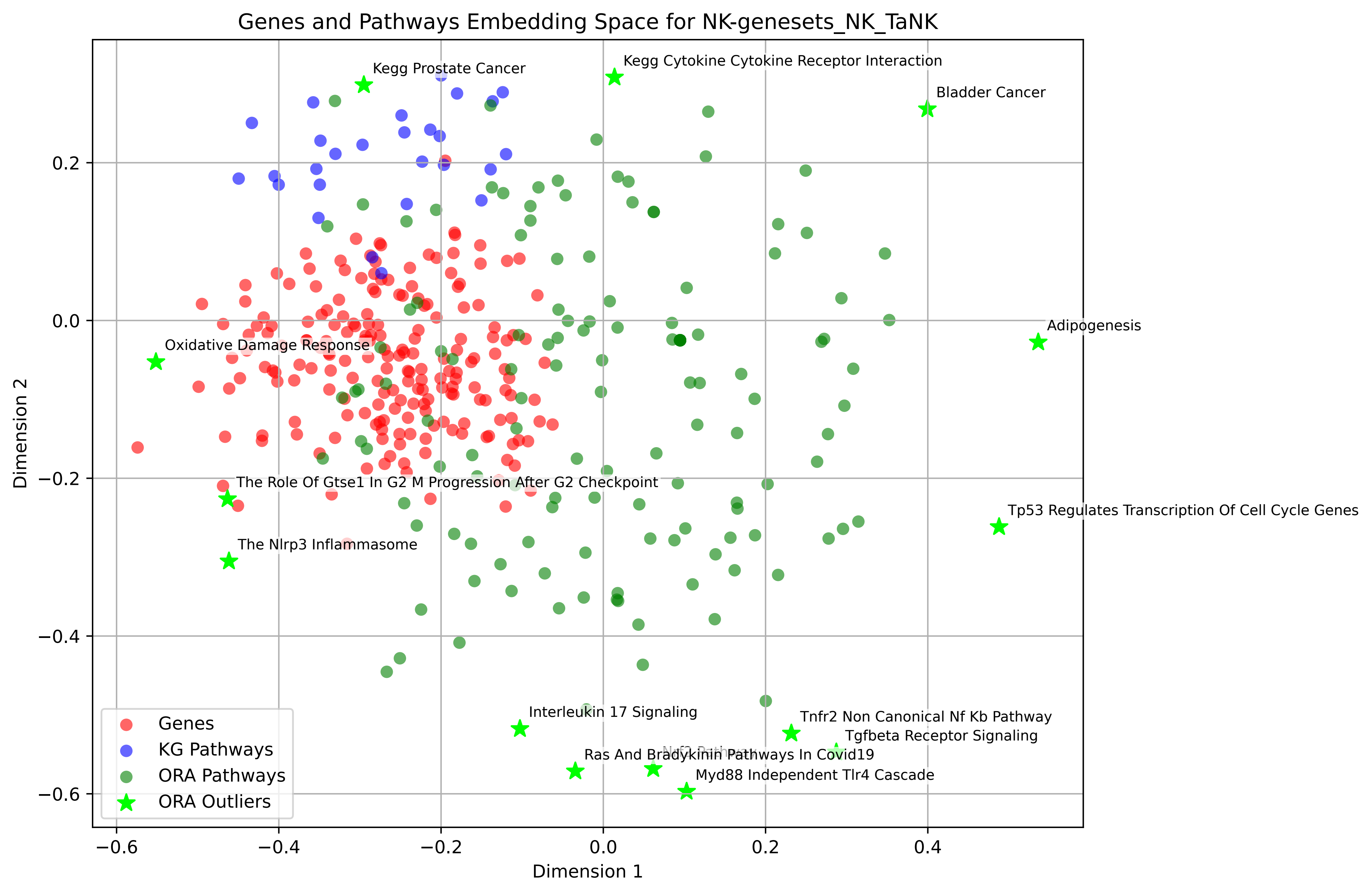
Genes and Pathways Embedding Space for NK-genesets_NK_TaNK
Dimension 1
Dimension 2
−0.6
−0.4
−0.2
0.0
0.2
0.4
0.2
0.0
−0.2
−0.4
−0.6
Kegg Prostate Cancer
Kegg Cytokine Cytokine Receptor Interaction
Bladder Cancer
Adipogenesis
Oxidative Damage Response
The Role Of Gtse1 In G2 M Progression After G2 Checkpoint
Tp53 Regulates Transcription Of Cell Cycle Genes
The Nlrp3 Inflammasome
Interleukin 17 Signaling
Tnfr2 Non Canonical Nf Kb Pathway
Tgfbeta Receptor Signaling
Ras And Bradykinin Pathways In Covid19
Myd88 Independent Tlr4 Cascade
Genes
KG Pathways
ORA Pathways
ORA Outliers

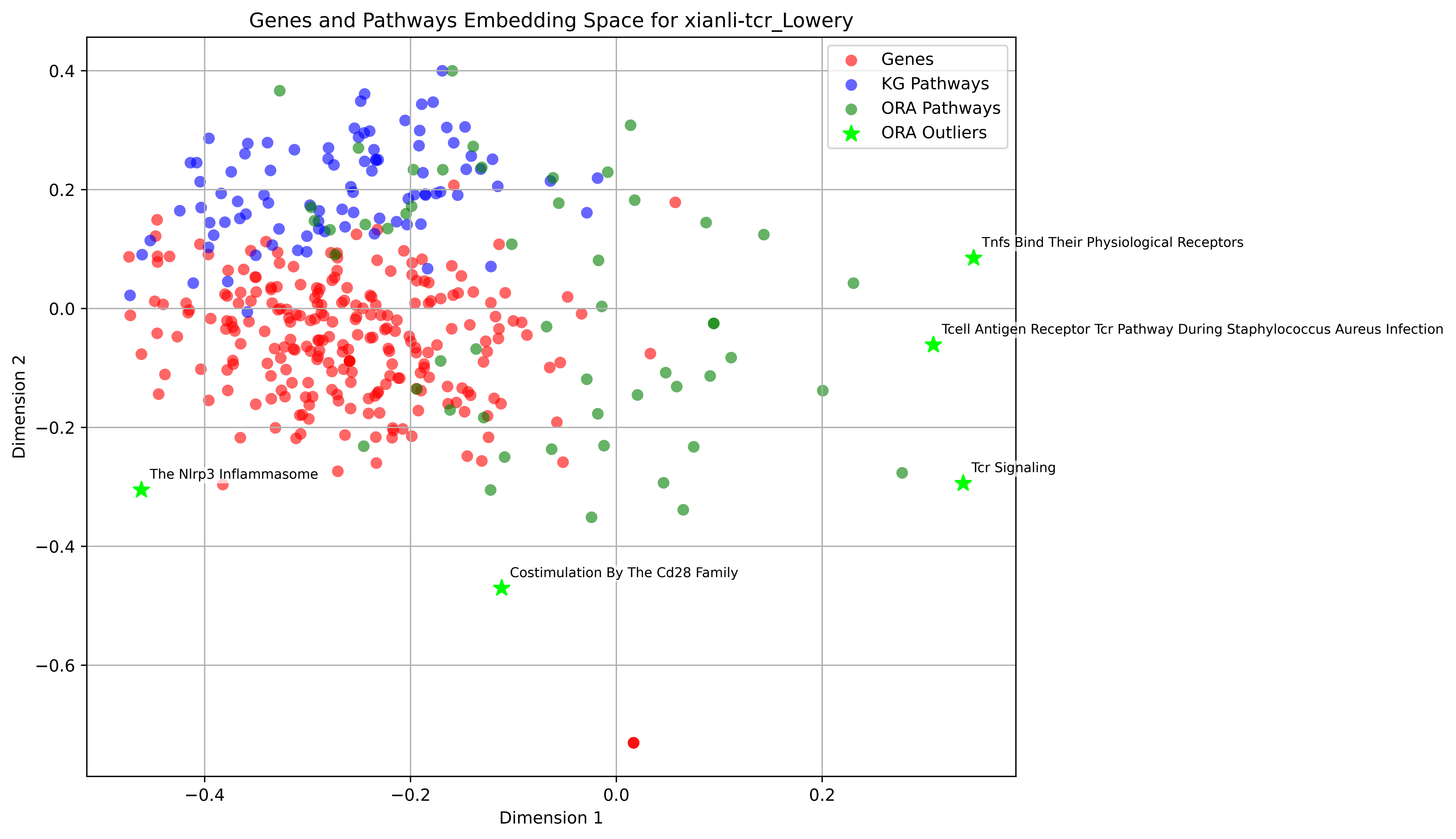

Genes and Pathways Embedding Space for xianli-tcr_Lowery
Genes
KG Pathways
ORA Pathways
ORA Outliers
Tnfs Bind Their Physiological Receptors
Tcell Antigen Receptor Tcr Pathway During Staphylococcus Aureus Infection
Tcr Signaling
The Nlrp3 Inflammasome
Costimulation By The Cd28 Family
Dimension 1
Dimension 2
0.4
0.2
0.0
−0.2
−0.4
−0.6
−0.4
−0.2
0.0
0.2

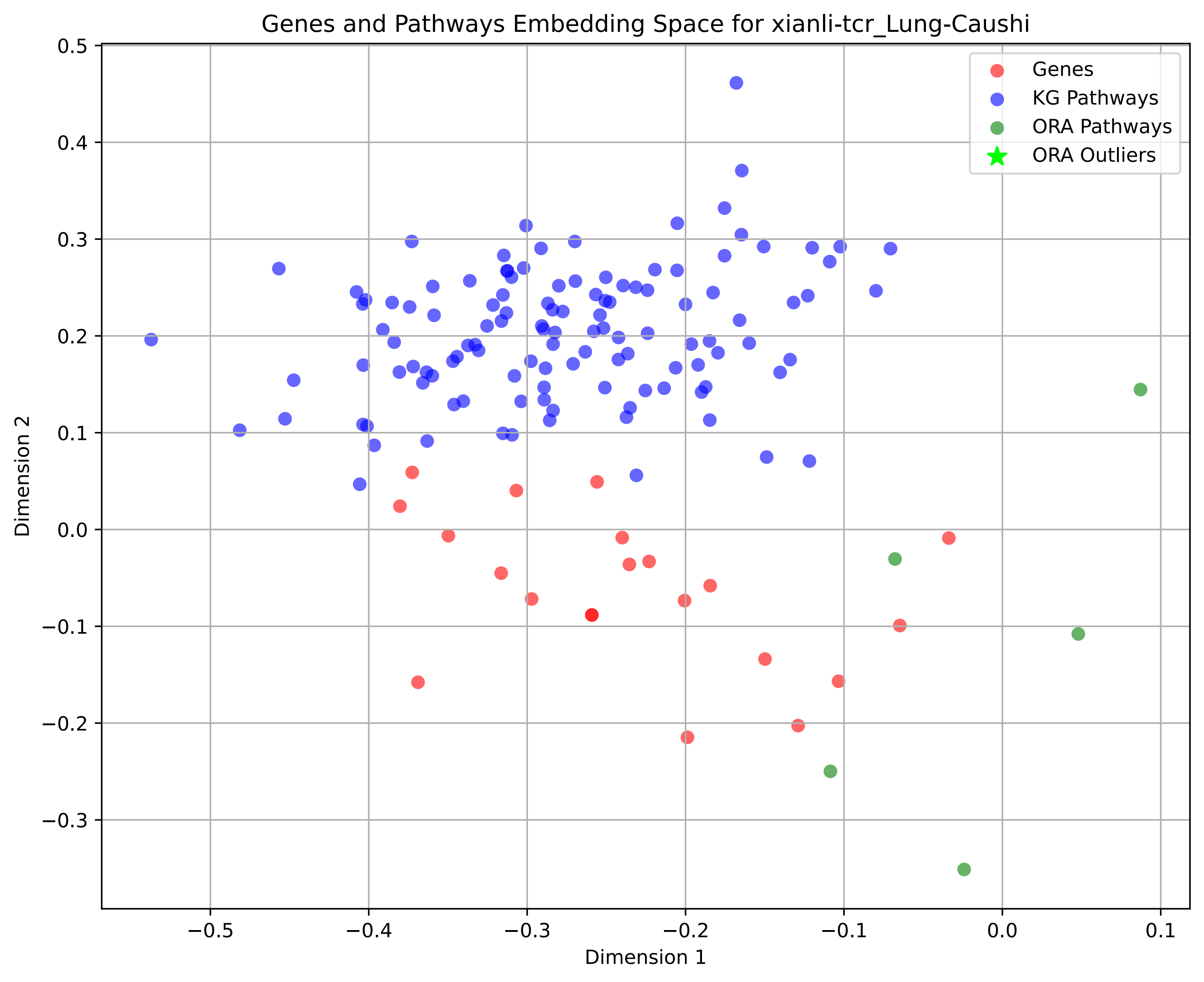
Genes and Pathways Embedding Space for xianli-tcr_Lung-Caushi
Genes
KG Pathways
ORA Pathways
ORA Outliers
Dimension 1
Dimension 2
0.5
0.4
0.3
0.2
0.1
0.0
−0.1
−0.2
−0.3
−0.5
−0.4
−0.3
−0.2
−0.1
0.0
0.1

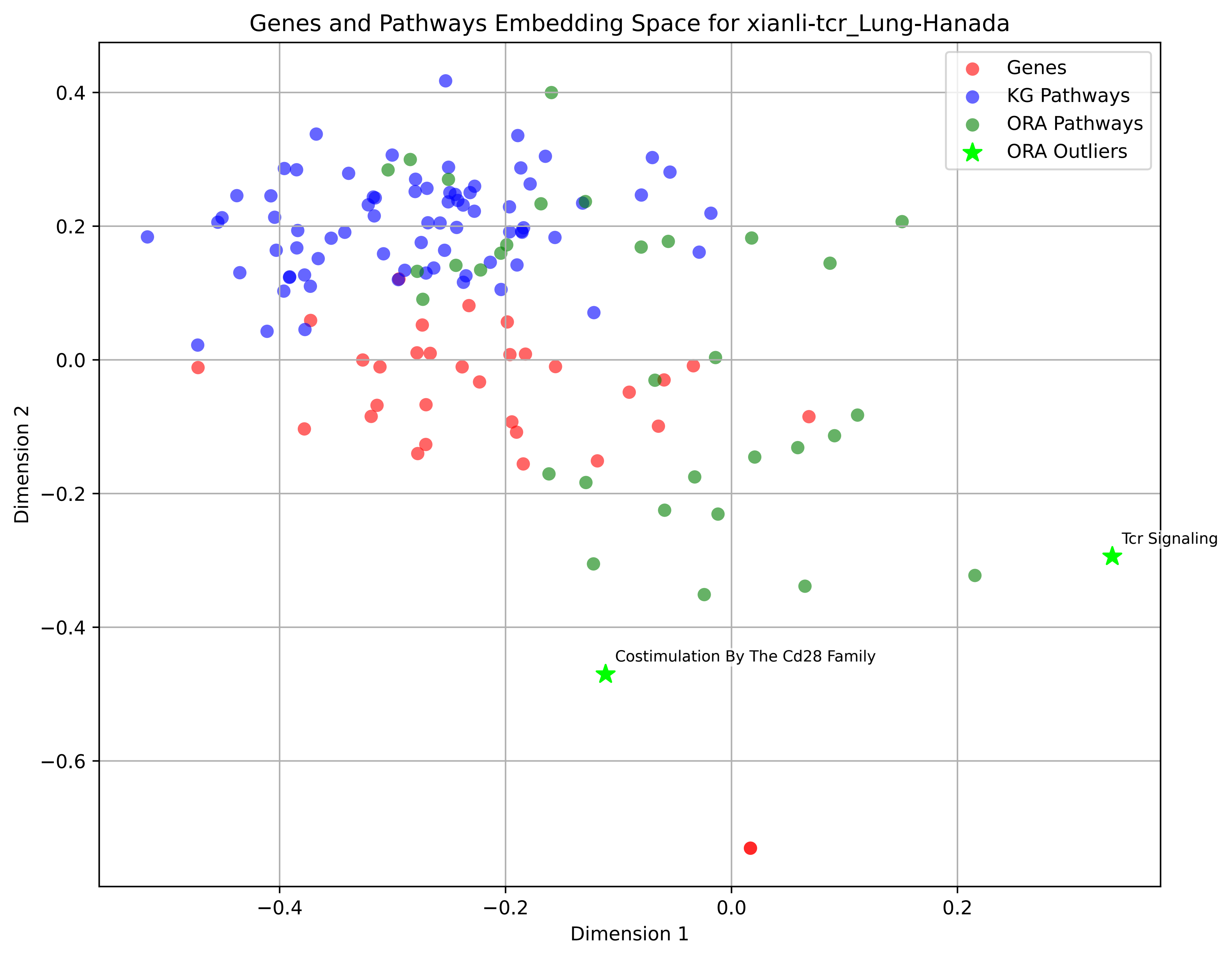

Genes and Pathways Embedding Space for xianli-tcr_Lung-Hanada
Genes
KG Pathways
ORA Pathways
ORA Outliers
Tcr Signaling
Costimulation By The Cd28 Family
Dimension 1
Dimension 2
−0.4
−0.2
0.0
0.2
0.4
−0.6

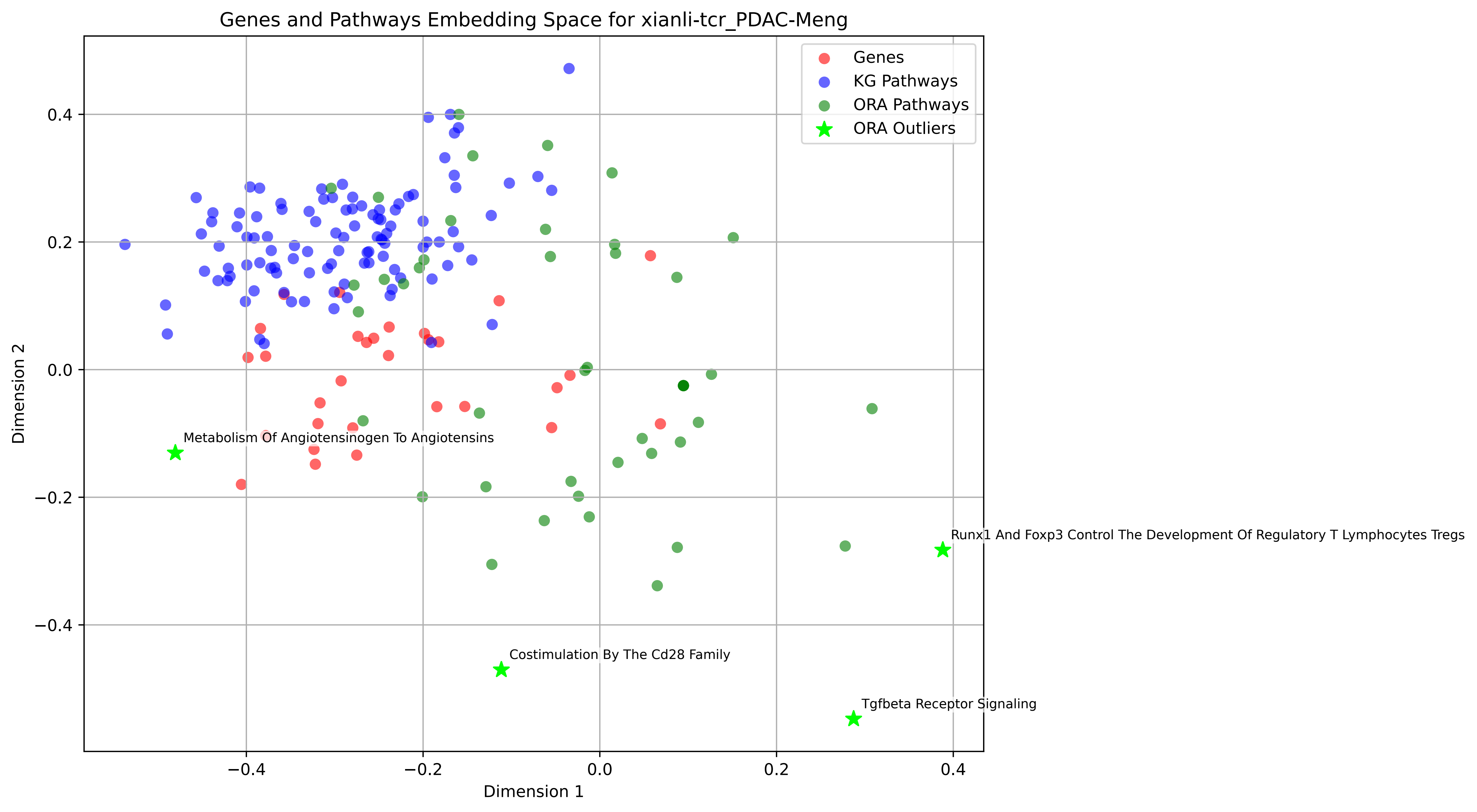

Genes and Pathways Embedding Space for xianli-tcr_PDAC-Meng
Genes
KG Pathways
ORA Pathways
ORA Outliers
Metabolism Of Angiotensinogen To Angiotensins
Runx1 And Foxp3 Control The Development Of Regulatory T Lymphocytes Tregs
Costimulation By The Cd28 Family
Tgfbeta Receptor Signaling
Dimension 1
Dimension 2
0.4
0.2
0.0
−0.2
−0.4
